# Integrated cycles for urban biomass as a strategy to promote a CO_2_-neutral society – a feasibility study

**DOI:** 10.1101/2021.04.22.440727

**Authors:** Nicole Meinusch, Susanne Kramer, Oliver Körner, Jürgen Wiese, Ingolf Seick, Anita Beblek, Regine Berges, Bernhard Illenberger, Marco Illenberger, Jennifer Uebbing, Maximilian Wolf, Gunter Saake, Dirk Benndorf, Udo Reichl, Robert Heyer

## Abstract

Progressive global warming is one of the biggest challenges civilization is facing today. The establishment of a carbon dioxide (CO_2_)-neutral society based on sustainable value creation cycles is required to stop this development. The Integrated Cycles for Urban Biomass (ICU) concept is a new concept towards a CO_2_-neutral society. The integration of closed biomass cycles into residential buildings enable efficient resource utilization and avoid transport of biowaste. In this scenario, biowaste is degraded on-site into biogas that is converted into heat and electricity. The liquid fermentation residues are upgraded by nitrification processes (e.g., by a soiling^®^-process, EP3684909A1) to refined fertilizer, which can be used subsequently in house-internal gardens to produce fresh food for residents.

Whereas this scenario sounds promising, comprehensive evaluations of produced amounts of biogas and food, saved CO_2_ and costs as well as social-cultural aspects are lacking. To assess these points, a feasibility study was performed, which estimated the material and energy flows based on simulations of the biogas process and food production.

The calculations show that a residential complex with 100 persons can generate 21 % of the annual power (electrical and heat) consumption from the accumulated biowaste. The nitrogen (N) in the liquid fermentation residues enables the production of up to 6.3 t of fresh mass of lettuce per year in a 70 m^2^ professional hydroponic production area. The amount of produced lettuce corresponds to the amount of calories required to feed four persons for one year. Additionally, due to the reduction of biowaste transport and the in-house food and fertilizer production, 6 468 kg CO_2_-equivalent (CO_2_-eq) per year are saved compared to a conventional building. While the ICU concept is technically feasible, its costs are still 1.5 times higher than the revenues. However, the model predictions show that the ICU concept becomes economically feasible in case food prices further increase and ICU is implemented at larger scale, e.g.; at the district level. Finally, this study demonstrates that the ICU implementation can be a worthwile contribution towards a sustainable CO_2_-neutral society and enable to decrease the demand for agricultural land.

## 1. Introduction

One of the most demanding challenges for the future is progressive global warming caused by excessive carbon dioxide (CO_2_) emissions and other greenhouse gases. To stop global warming, our society must reduce the CO_2_ emission and make our entire lifestyle CO_2_-neutral. While many concepts for sustainable electrical energy production already exist, CO_2_-neutral agriculture and biomass circulation concepts are lacking. And, since half of the world population now lives in cities, these concepts have to be also applicable to urban areas. For example, in some urban districts (e.g. the Jenfelder Au in Hamburg, Germany (Hertel et al., 2015)) black water is used on-site to produce heat and electricity by anaerobic digestion (AD). Furthermore, roof-top gardens enable the production of food in the cities (Barreca, 2016). Fuldauer *et al.* (Fuldauer, 2018) demonstrated that it is even feasible to connect a small-scale anaerobic digestion plant (ADP) with a hydroponic or algae cultivation system to close the biomass cycle. The benefits of closed urban biomass cycles are an efficient utilization of the resources and the avoidance of transport (Jouhara, 2017).

This study investigates the potentials and limitations of a concept for urban biomass circulation regarding energy and food production, carbon dioxide equivalent (CO_2_-eq) savings, costs, and social-cultural aspects in Germany.

The concept, called Integrated Cycles for Urban Biomass (ICU), demands an in-house ADP to degrade biowaste from residential buildings to biogas and digestate. The biogas generated is converted on-site to heat and electricity through a combined heat and power plant (CHP). The remaining fermenter liquid is upgraded by a soiling^®^-process (EP3684909A1) and a nitrification process to refined fertilizer. Finally, the liquid fertilizer is used to produce fruits, vegetables, and ornamental plants using either in-house integrated hydroponic systems, soil-based agriculture or roof-top gardens. Finally, the residents can consume the food while the accruing plant residues are fed into the ADP to close the biomass cycle again (Fig. 1). Whereas soil-based agriculture is more robust, the use of hydroponic systems for the production of vegetables enables faster growth, higher product quality and needs less space (Sapkota, 2019).

**Fig. 1:**
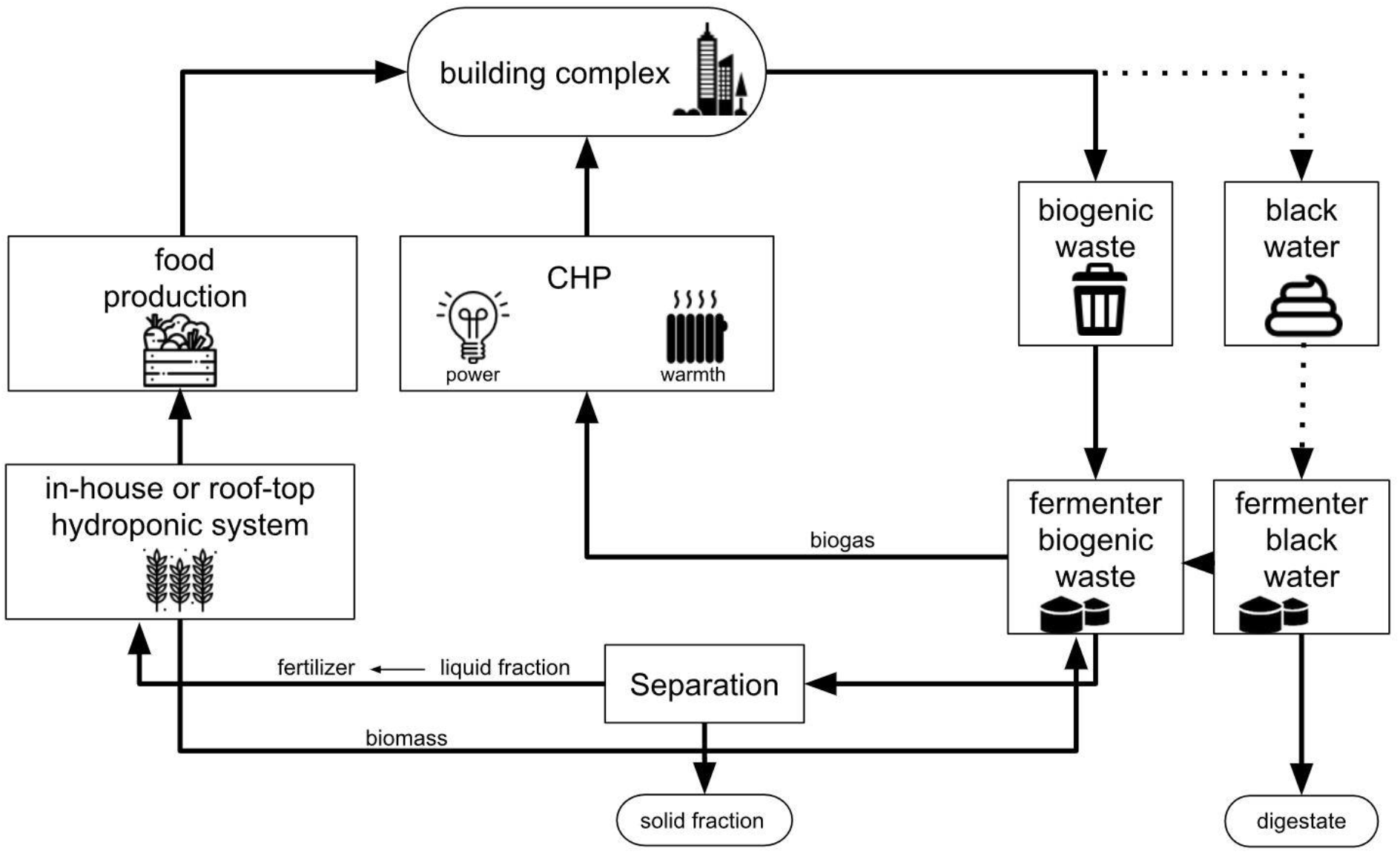
Process chart of ICU concept. ◻: Integrated Cycles for Urban Biomass (IUC) processes. **O**: Input of biomass through the building complex into the process; outputs like solid fraction and digestate. “^….”^: additional blackwater for biogas production.

A key challenge to close the biomass cycle between anaerobic digestion (AD) and agriculture is transforming the digestate into fertilizer. Digestates contain high amounts of ammonium (NH_4_). But, while NH_4_ can be used as a nitrogen (N) source by plants, high NH_4_ contents potentially increase N-losses by emission and can inhibit plant growth, especially in hydroponics. Therefore, NH_4_ has to be oxidized via nitrite (NO_2_) to nitrate (NO_3_) (e.g. by the soiling^®^-process). In particular, for hydroponic-based crop production, fertilizer quality is of high importance as, among others, its buffer capacity is very low (compared to soil). For hydroponics, under optimal conditions, synthetic or inorganic-based fertilizers are commonly applied. However, organic-based nutrient solutions such as fermenter digestates are also suitable, while the nitrification step needs to be implemented for deriving plant-available N-forms (Krishnasamy et al. 2012, Shinohara et al. 2011, Stoknes et al. 2016). With the adjustment of the correct dilution ratio and nutrient concentrations of the organic fertilizers, similar or even higher yields compared to a commercial nutrient solution are possible (Liedl et al. 2004, Wang et al. 2019). Finally, based on a Life Cycle Assesment (LCA), the reduction of CO_2_-eq can be calculated (Lombardi et al., 2003).

## 2. Methods

To assess the ICU concept regarding energy and food production, the conversion of biowaste to heat and electricity using AD (section: 2.1.) and agriculture (section: 2.2 and 2.3) were simulated. CO_2_-eq reductions as an indicator for the global warming potential of parts of the ICU process were evaluated by LCA (section: 2.4). In addition, the costs for the implementation of the ICU concept in real buildings were estimated (section: 2.5). Finally, social-cultural aspects of the implementation were reviewed for Germany (section: 2.6).

### 2.1 Modelling in-house biowaste degradation for energy production

Biogas production was used as a key process to model biowaste conversion to heat and electricity. The entire process was simulated using pre-implemented building blocks from the software SIMBA#Biogas (https://www.ifak.eu/de/produkte/simba-biogas, ifak, 2020) (Fig. 2) assuming a building with 100 residents.

**Fig 2.**
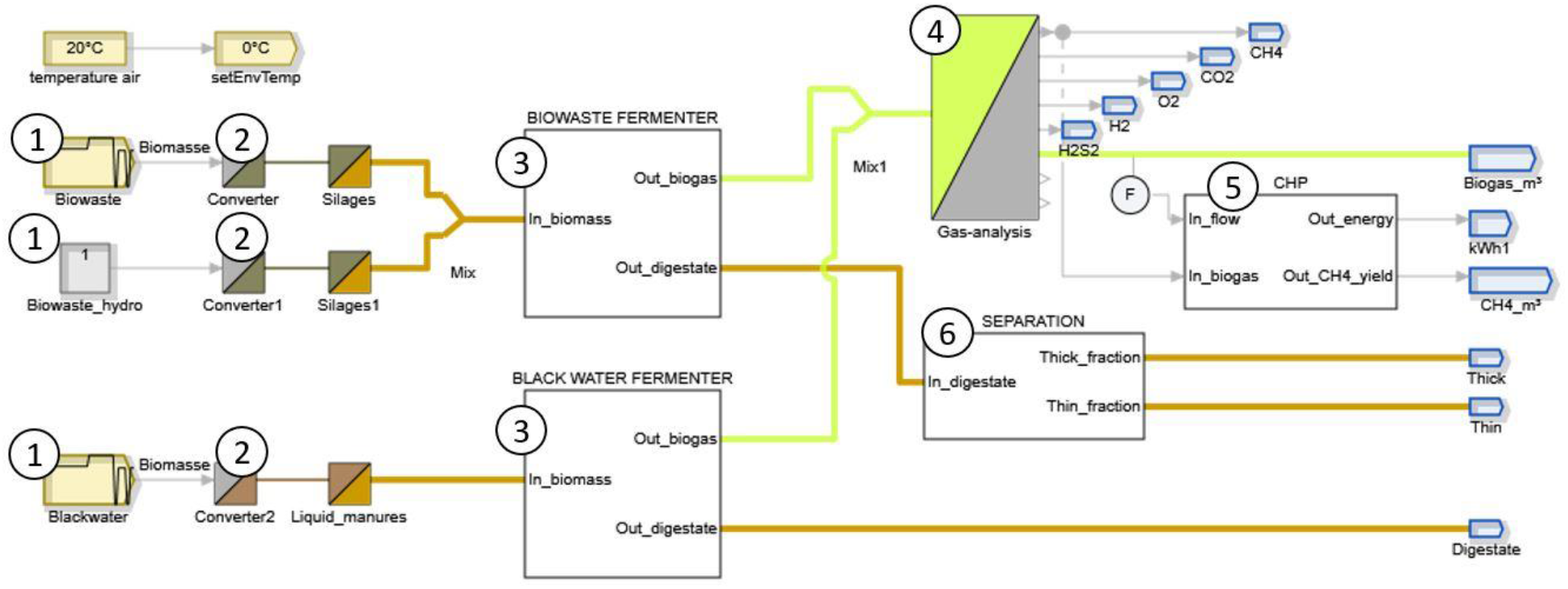
Simba model for in-house degradation of biowaste to heat and electricity. 1. Biomass input of biogenic waste and blackwater with 70 °C pre-treatment. 2. The Converter block determines the biomass composition. 3. The Fermenter block determines the biogas and digestate output. 4. The Gas analysis determines the biogas composition. 5. The CHP block converts biogas into energy with a calorific value of 10.5 kJ/kg 6. The Separation block determines thick (solid) and thin (liquid) fractions.

The “Biowaste block” (input) assumes homogenization at 70 °C to ensure the necessary sanitization conditions. Therefore, the biomass will be pretreated in a homogenization tank for 2-3 days before fermentation (Luste et al., 2010). On average, residents are supposed to produce 33.6 kg of biogenic waste and 720 l of blackwater per day. Since the amount of biowaste fluctuates over the year, an increase of 22 % for four months and a feeding of 30.7 kg for five days a week was applied to monitor the dynamic behavior (Hanc, 2011, supplementary file 1). The “Converter block” takes into account the biomass composition based on literature values (supplementary file 1, Malakahmad et al., 2008, Hanc et al. 2011). The “Biowaste Fermenter block” represents the conversion of kitchen and hydroponic biowaste into biogas and digestate. The simulations use the Anaerobic Digestion Model Number 1 da (ADM1da), which is an extension of the Anaerobic Digestion Model Number 1 (ADM1) (Karlsson et al., 2017). The ADM1da comprises 32 differential and algebraic equations. They represent all relevant steps of the biomass degradation and physicochemical process parameters. Operation of the ADP at 55 °C was assumed. The “Gas analysis block” defines the biogas composition. The “CHP block” is used to determine the methane (CH_4_) yield into electrical energy and heat. Here, an electrical efficiency of 38 % and a thermal efficiency of 45 % was used (Liebetrau et al. 2020, Scheftelowitz et al. 2013). The electrical efficiency increases with the purity of the AD product gas (Liebetrau et al. 2019). The “Separation block” is used to split the digestate into a thin (liquid) and a thick (solid) fraction. The liquid effluent is further processed and nitrified by the soiling^®^-process into refined digestate. Soiling^®^-nutrient recycling fertilizer is composed of mineral N plus macro- and micronutrients. Note: only the N amount can be taken into account with Simba; for further calculations the macro- and micronutrients were neglected.

For the “Blackwater block” (input) a second fermenter is considered as there is currently no approval for a fertilizer containing anthropogenic raw material in Germany according to the so-called Düngemittelverordnung (DümV, 2012). Therefore, blackwater fermentation is only considered for energy production but not for fertilizer production. Furthermore, the “Blackwater Fermenter block” is modeled by three different reactor block configurations to identify the most efficient one. The first scenario considers biogas production inside a continuously stirred-tank reactor (CSTR). The second scenario takes into account five CSTR blocks connected in series to simulate a plug flow bioreactor (PFR). The third scenario describes a two-stage reactor (2sR). Here, a small CSTR is used for hydrolysis and fermentation of biowaste, whereas acidogenesis and methanogenesis occur in a bigger second CSTR. The specific parameter settings for all scenarios are in the supplementary file 2a, 2b, 2c, 2d.

The ADP considered in this work was assumed to produce biogas with a calorific value of 10.5 kJ/kg Thus, neither combined cycle power plants (CCPP) nor CHP can be applied on-site because of their lower efficiency. In practice, many operators of small-scale biogas plants favour a satellite CHP over on-site power production. A satellite CHP is supplied with biogas from multiple small-scale biogas plants via a local micro gas grid (Scheftelowitz et al. 2013). The assumption is that a large number of small-scale biogas plants are nearby; an option that could also be applied here. Suitable for the on-site power production of small-scale biogas plants are fuel cells, microturbines (MT), and engines (igniting beam engine, gas engine). Fuel cells can reach high electrical efficiencies and run quietly. However, they are comparatively large, expensive and their operation requires a high gas purity, which would make additional biogas upgrading necessary. Therefore, the use of a fuel cell was not considered in this study. MT, on the other hand, can operate with a wide range of CH_4_ concentrations (30-100 %) (Scheftelowitz et al. 2013). MTs reach an electrical efficiency of 25-33 % with a thermal efficiency of ~49 % (Lugmayr, 2010), and operate silently and environmentally compatible (Hasemann, 2015). MTs are commonly applied on a 30-550 kW scale (Scheftelowitz et al. 2013, Lingstädt et al., 2018). Commercial 1 kW scale turbines are under development (ENBW, 2021). However, at the current state, small-scale implementations are inefficient (11 % electrical efficiency) at costs of around 6 000 € per turbine (Haseman, 2018, Agelidou et al. 2019). Alternatively, engines (igniting beam engine, gas engine) can reach a higher electrical efficiency than MTs of 30-40 % and a thermal efficiency of ~47 %. Compared to MTs, engines are louder, produce noxious side products and require more maintenance. In particular igniting beam engines, which require the addition of pilot oil for combustion, produce noxious side products and soot, which inhibits the efficient use of excess heat (Lugmayr, 2010). The preferable alternative are gas engines, which operate without pilot oil but require CH_4_ concentrations above 45 % (Lugmayr, 2010).

In summary, gas engines were considered the most attractive option for on-site production of electrical energy from AD product gas in this study. However, if other small-scale biogas plants were available, a satellite CHP could be the more efficient alternative.

### 2.2 Crop production systems

The amount of total N (N_tot_) (N_tot_ = N_org_ + NH_4_-N) is the main input to calculate the liquid effluent. An optimal, highly efficient system with biologic activity efficiency of 1.0 was assumed, i.e., 100 % of N_tot_ was transformed into plant-available nitrate-nitrogen (NO_3_-N) by the soiling^®^-process. For cultivation planning (system sizing), the ratio of fresh biomass production to available N was considered. As model crop lettuce (*Lactuca sativa ssp.*) was used. A fresh matter N content of 0.18 % was assumed (Feller et al. 2019) with a fixed dry matter fraction of 0.048 %.

Four possible methods were considered for lettuce cultivation: Scenario 1 and 2 are open-air plant cultivation systems with raised beds or vertical hydroponics, respectively. The residents drive these scenarios on the roof-top with a cultivation period from April to October (vegetation period of Berlin). Both scenarios are complex as they involve the participation of community members (that are outside of the scope of the present simulations). Scenario 3 and 4 are protected cultivations with hydroponic greenhouses or plant factories, respectively. Both have to be operated year-round by trained staff and can be located on the roof-top or in the basement of buildings. Here a pure bio-technical assessment using deterministic explanatory simulation models was applied.

A numerical simulator for controlled environments and greenhouses was used that is a further development of earlier published greenhouse simulators (Körner and Hansen, 2012; Körner et al., 2008). The simulator was programmed using MATLAB (MathWorks Inc., USA). It was connected to a replica of commercially available climate controllers, including a setpoint generator that calculated climate setpoints for heating, ventilation, light and CO_2_ concentration. The simulator was fitted to a standard Venlo-type greenhouse structure or a vertical farming hydroponics-controlled environment (scenario 3 and 4). The simulator’s crop-basis is a photosynthesis-driven growth model with microclimate predictions for water and nutrient uptake according to the Penman-Monteith equation (Körner et al., 2007). Nutrient uptake was calculated assuming that the diluted nutrients in the irrigation system are optimally taken up by the crop. As such, a perfect pH, electrical conductivity (EC), and a root environment with optimal nutrient solution composition with an optimal availability of all nutrients were assumed. In accordance with Goddek and Körner (2019), all element-specific chemical, biological or physical resistances were set to zero.

For technical layout, supplementary lighting was applied with LED lamps installed either under the roof above the crop with an installed capacity of 80 W m^−2^ power and an output of 192 μmol m^−2^ s^−1^ or at an installed capacity of 110 W m^−2^ power and an output of 264 μmol m^−2^ s^−1^ in scenario 3 or 4, respectively. The light was controlled dynamically with setpoints generated using a daily light integral (DLI) of either 12 mol m^−2^ d^−1^ or 20 mol m^−2^ d^−1^ for greenhouse or vertical farming, respectively (Körner et al., 2006). In both scenarios, CO_2_ in the air was set to 700 μmol mol^−1^ and supplied according to the demand (max. at 15 g m^2^ h^−1^) during lightening when greenhouse vents were closed and at all times in the vertical farming scenario. In the greenhouse scenarios, heat exchange for cooling was calculated with passive roof ventilation while active cooling and dehumidification were used in the vertical farming-controlled environment scenarios (active cooler based on ANSI/AHRI standards 1200 (Anonymous, 2013). Dehumidification was implemented with a commercially available dehumidification unit of the type ventilated latent heat energy converter. Further model parameters are summarized in the supplementary file 3. The simulator calculated macro- and microclimate in a time-step of 5 min, integrated hourly using controlled actuators (e.g., heating, ventilation, cooling, CO_2_, light) that were re-adjusted as described by Körner and Van Straten (2008). The simulations’ output included hourly biological and physical variables related to lettuce production, such as microclimate conditions, photosynthesis, yield, and resource consumption (electrical power, heating energy, water, CO_2_). Input to the simulation program included, among others, physical location (latidude (LAT), longitude (LON)), humidity set point (%), set points for heating and ventilation (°C), crop planting density (plants m^−2^) and temperature-sum related harvest time. Input climate data were hourly data sets for Berlin (Germany, LAT 52.5N, LON 13.4) from 2009 to 2018 (Meteoblue; www.meteoblue.com). Calculations were performed for all scenarios for single years of each of the 10-year horizons.

Simulations were performed targeting nutrient and water uptake, yield, and energy demand for heat and lighting for either a greenhouse with a size of 70 m^2^, or for a vertical farm with a four-layer system (17.5 m^2^ each, in a room of 30 m^2^ area and 2.50 m height). As commercially viable climate control in small greenhouses is challenging to maintain, a 500 m^2^ greenhouse as minimum commercial size was modeled in addition. All simulations were done for year-round production of hydroponically grown lettuce with a fixed planting density of 36 plants m^−2^ as in commercial practice (e.g., Brechner et al., 2013).

### 2.3. Estimation of the CO_2_ saving potential using life cycle assessment with openLCA

The ICU concept offers the opportunity to save CO_2_ due to reduced transport of biowaste and food (Finkbeiner et al., 2006). To quantify the amount of saved CO_2_, a Life Cycle Assesment (LCA) with the open-source software openLCA (version 1.10.3) was conducted. This software considers the total energy consumption by all components at various levels. The used database was ecoinvent35_Cut (Wernet, 2016). The system environment was divided into five phases: extraction of raw materials and energy sources, manufacture, use, transport and disposal (Fig. 3.A) (McDonough, 2010). The boundaries for the ecological assessment are shown in Fig. 3.A (grey dots). An average distance of 30 km for the transport of biowaste to the ADP in the conventional scenario was assumed. In the ICU concept, transport was neglected (Fig. 3.B). All flows and process data are found in the supplementary file 4. The system contains specific elements providing the functional unit of 1 kg biomass for the complete life cycle. The input flow is 11 t with bio-degradable garden and park waste, food and kitchen waste from households, restaurants, caterers and retail stores, comparable waste from food processing plants as well as forestry or agricultural residues, and manure. It does not contain sewage sludge or other biodegradable waste such as natural textiles, paper or processed wood.

**Fig. 3:**
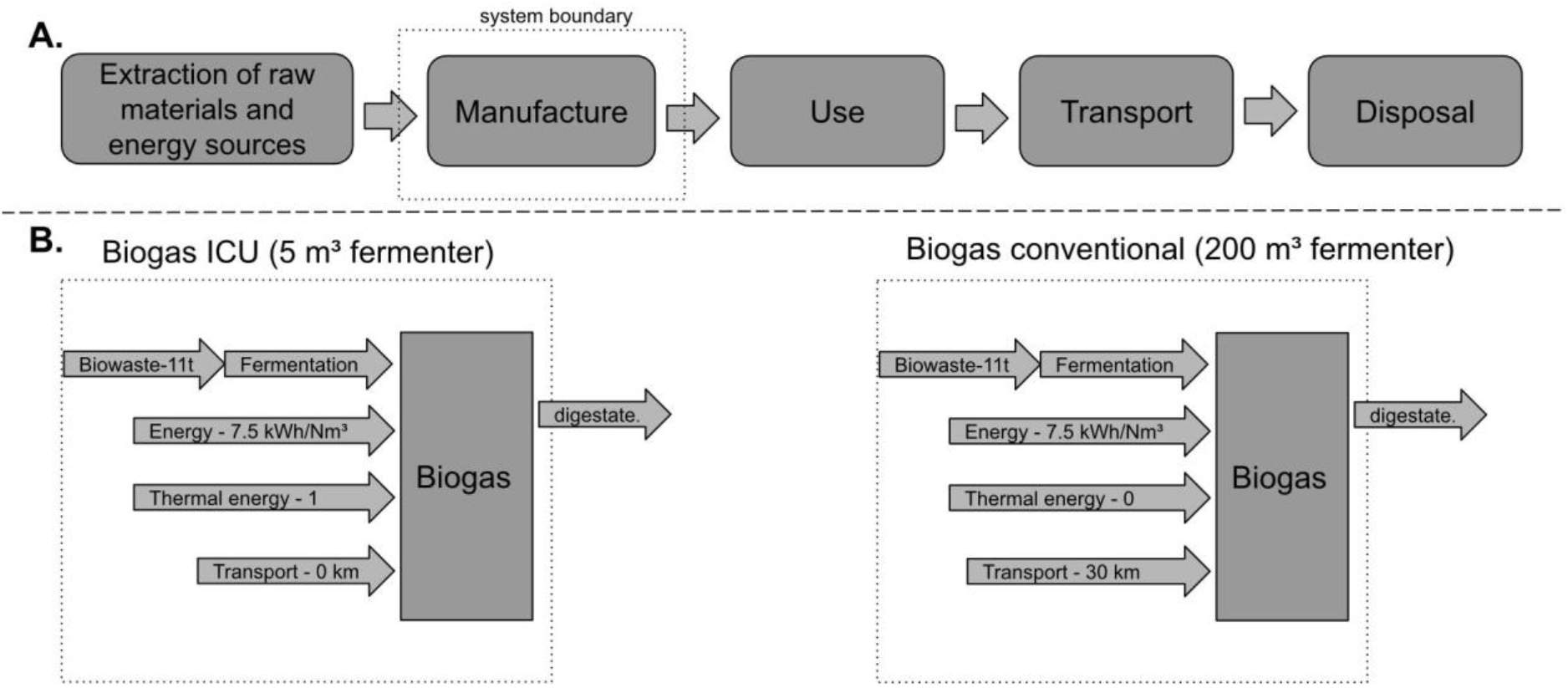
System boundaries of the Life Cycle Assessment study. A: System environments of the product life-cycles based on DIN 15804 in four steps and system boundary (functional unit 1 m^3^ biogas). B: System boundary for the ICU concept in comparison to a building with transport of biowaste to a conventional biogas plant. (Energy: https://biogas.fnr.de/daten-und-fakten/faustzahlen)

### 2.4 Cost calculation

The costs of implementing the ICU concept were estimated by a life cycle cost analysis (LCCA, equation 1) for the example of a building with 100 residents. Therefore, the costs for acquiring and operating the fermenters, the soiling, the hydroponic systems, the technical staff and the building’s space were considered taking into account the net present value (NPV).

Investment costs (C) were depreciated for a period of 20 years. Replacement costs were assumed with 10 % of the investment costs after 20 years and maintenance costs (A+M) with annually 5 % of the investment costs. Energy costs (E) were omitted since the system produces the required energy on its own. Also, the resale value (R) of the installations was assumed as “0” € as it was expected that the building’s value remains at least stable. Additionally, ADP operation and plant cultivation require an experienced worker requiring at least 25 € per hour.

The production of in-house biogas generates energy in form of electricity and heat. This energy is reused inside the ICU building. If the generated energy were sold, the price for 1 kWh heat and 1 kWh electricity would be 0.024 € (Andor et al., 2018) and 0.13 € (§43 EEG 2017), respectively. A summary of results obtained is found in table 3 (section 3.4).

Against these costs, the value of the produced energy and food was taken into account. The benefit of the reduction in the disposal of biowaste and wastewater was neglected to avoid further complication of the calculation. Furthermore, the installation of vacuum toilets and separate black and grey water tubes is also cost-intensive. However, the cost of the installations compensates with the benefit of a reduced wastewater volume.

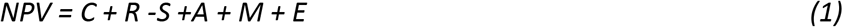

C = investment costs
R = replacement costs
S = resale value at the end of study period
A = sum of annually recurring operating, maintenance and repair costs
M = non-annually recurring operating, maintenance and repair cost
E = energy costs

### 2.5 Overview of important social-cultural aspects required for the implementation

To assess social-cultural aspects for the implementation of the ICU concept in Germany, a literature survey was performed addressing the following questions:

- How great is the interest of the residents in urban agriculture and sustainable lifestyles?
- How great is the willingness of real estate owners to implement an ICU concept?
- How important is it for the government to achieve a carbon-neutral society?
- Which legal paragraphs have to be considered for implementing an ICU concept?
- Which additional social-cultural aspects might be relevant for implementing an ICU concept?

While for most of these questions results and data from literature already exist, the question on the real estate owners’ willingness to implement an ICU concept has not been addressed, so far. Therefore, an online survey to collect this data was performed. To obtain a comprehensive picture of the attitude towards this new concept, 235 real estate owners, about 15 from every Federal State, were selected. All owners received a short online questionnaire containing ten questions (supplementary file 5) to rate to which extend different aspects of the ICU concept and its implementation are important to them. In the end, only 14 answers were received.

## 3. Results and Discussion

This feasibility study evaluates the amount of energy (section: 3.1) and food (section: 3.3), which could be produced by implementing an ICU concept for a building with 100 residents. Before the final simulation, the ADPs’ fermenter size and configuration were evaluated and the best scenario for house-internal food production was selected. A precondition for plant growth was converting NH_4_ in the digestate to NO_3_ (section: 3.2).

Based on the ICU-concept’s best scenario, the costs were calculated (chapter: 3.4) and the potential CO_2_-savings (section: 3.5). Finally, social-cultural aspects were reviewed, including the laws required for implementing the concepts (section: 3.6) and potential addons for the ICU-oncept (section: 3.7).

### 3.1.1 Utilization of biowaste by optimized anaerobic fermenters enable to cover 21 % of the annual energy demand of the building

As the first step, the size and performance of CSTR, PFR, and 2sR ADP for processing of biowaste (Fig. 4) and black water (Fig. 5) were compared based on the energy content of the biogas. The volume ratio between the hydrolysis and the main fermenter of the 2sR was 1:50 as determined in the supplementary table 6. For the PFR the sum of all five fermenters connected in series was assumend for the simulation.

**Fig. 4:**
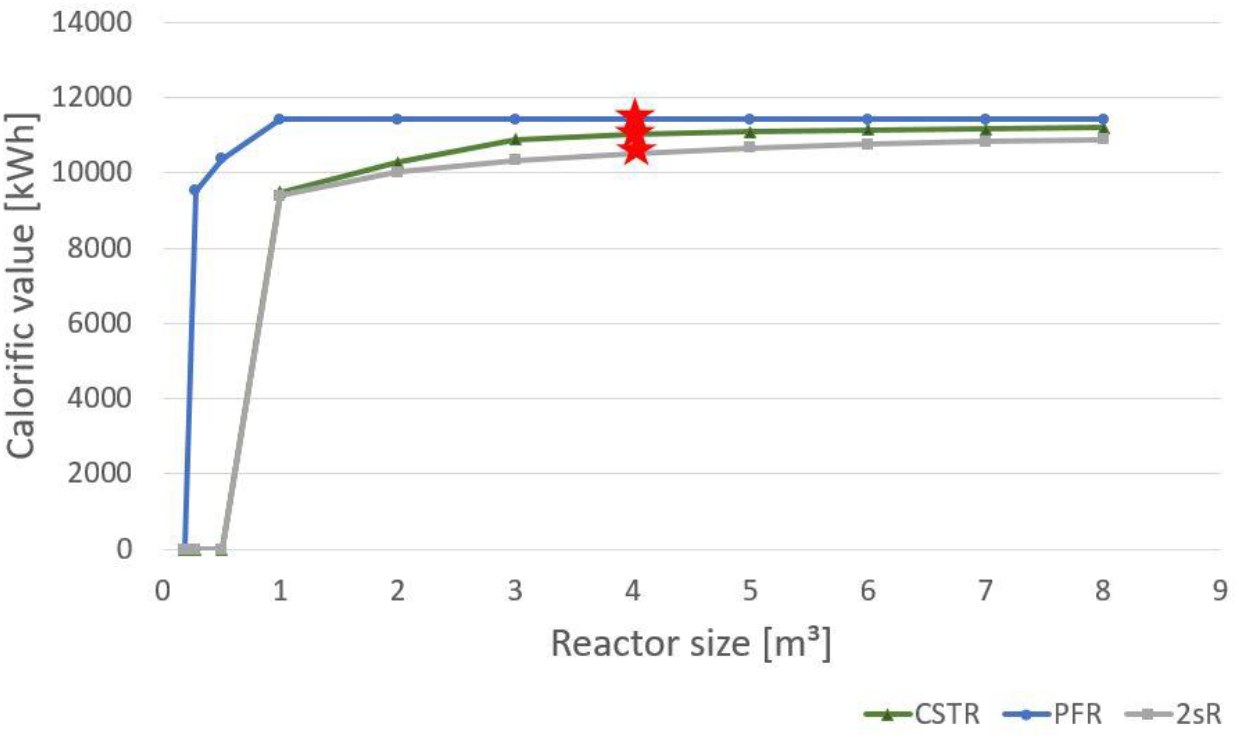
Evaluation of reactor scenarios with biowaste fermenters. Energy content of the biogas produced annually (without losses). Only biowaste input. For PFR reactor the sum of all 5 fermenters and for 2sR is hydrolyse + main fermenter are considered. Star shows the chosen reactor size.

**Fig. 5:**
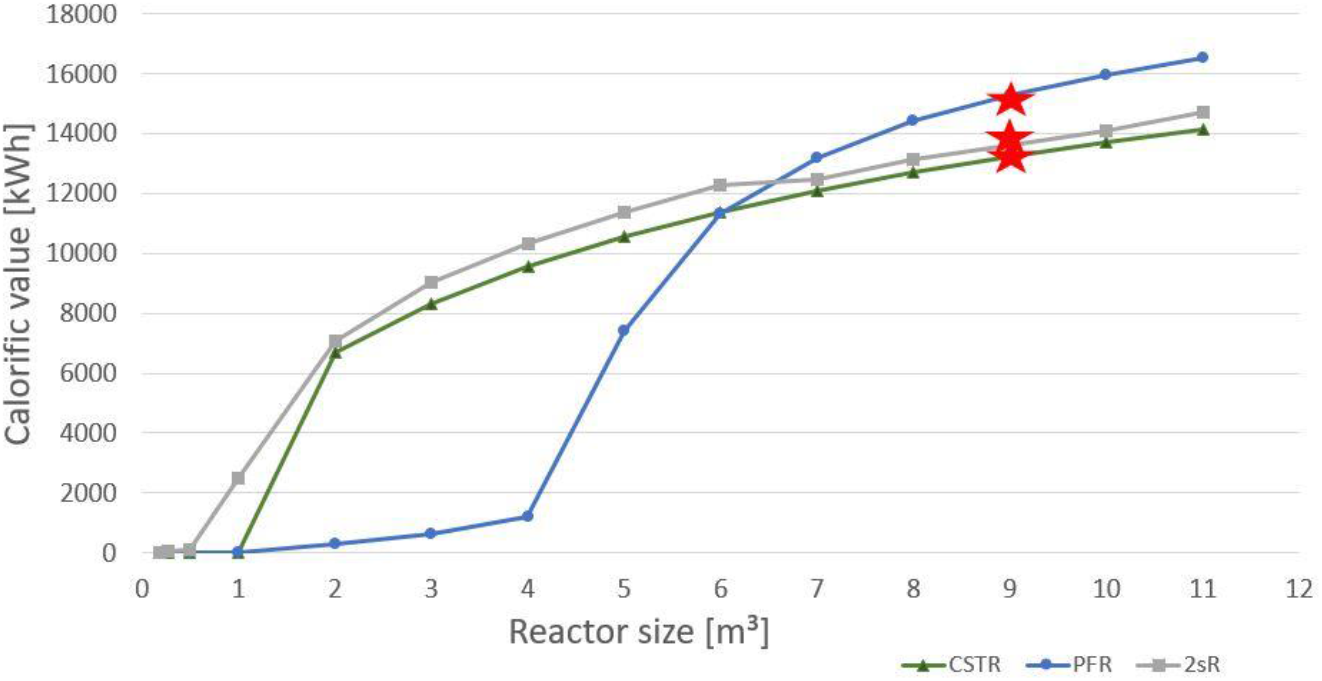
Evaluation of reactor scenarios with additional blackwater fermenter. Energy content of the biogas produced annually (without losses). Blackwater input. For PFR reactor the sum of all 5 fermenters and for 2sR is hydrolyse + main fermenter are considered. Star shows the chosen reactor size.

**Fig. 6:**
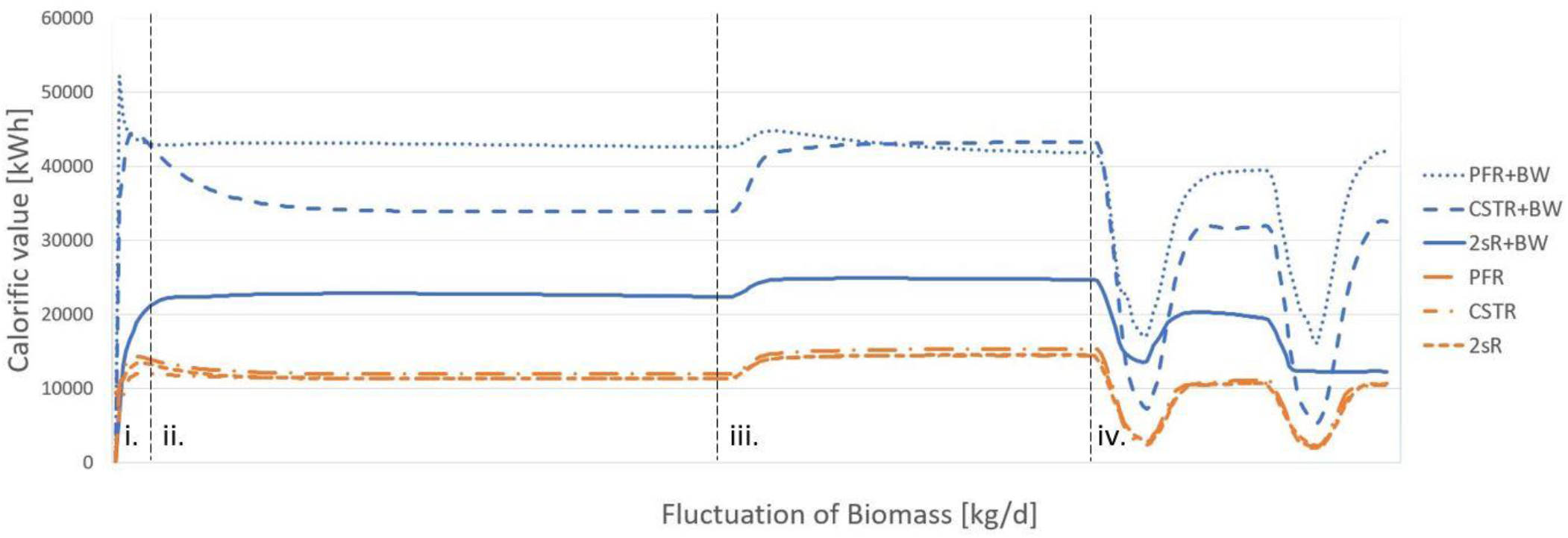
Dynamic behavior of the ICU model. Feed fluctuations: Energy production from biowaste (green) and additional black water (BW) (brown) for 2sR, CSTR and PFR fermenters. i: Initiation phase of the model. ii. Feeding rate of 33.6 kg/d iii. Feeding rate of 41.1 kg/d iv. Feeding rate of 30.7 kg/d for five days a week.

The first scenario was the calculation of biowaste input in one fermenter with an average amount of 33.6 kg biowaste per day (Fig. 4). Depending on the reactor size 1 kW energy may be produced. The second scenario calculates the addtional black water fermentation in a second fermenter with an average of 720 L black water per day (Fig. 5). Black water may improve the daily energy yield to 3 kW energy, fitting to the values of studies with similar substrates (Wriege-Bechtold et al. 2015).

For the first scenario the simulation of PFR produces about 5.5 % more energy than the two-step and 22 % more than the CSTR fermenter. These magnitudes between the fermenter types were also shown by Bensman et al. (Bensmann, 2013). Additionally, a PFR is more robust against contaminants like plastic material in biowaste. The shape of the power to fermenter size curve is sigmoid, reflecting that too small fermenter sizes lead to acidification, whereas too large fermenter adds no further benefit (Fig. 4). As optimal biowaste fermenter sizes were chosen a 4 m^3^ CSTR-fermenter, five fermenters connected in series with each 1 m^3^ for the PFR-fermenter and also 4 m^3^ for the main fermenter of the 2sR (Fig. 4). The PFR-fermenter was selected as optimal because with 11.434 kWh calorific value annually it was able to produce the most energy. This amount of energy corresponds to 9.5 % of the annual energy demand of 100 persons (Frondel et al., 2015). Since the reactors with their control units require less than 20 m^2^, installation in the technical center of a building is technically feasible. Production of heat and electricity would require an additional CHP unit of about 10 m^2^ size. Alternatively, the biogas can be used to cook and climatize the building. This scenario requires that the building have gas heating/heating pumps instead of an oil or electric system.

In comparision the second scenario with an additional black water produce 25,855 kWh energy. This scenario is ecological more efficient to the fist one because it can cover 21 % of the yearly energy production (Fig. 5). As optimal reactor size for the CSTR a 9 m^3^ fermenter, for PFR five fermenters connected in series with each 1.8 m3, and for 2sR 9 m^3^ were choosen.

Because of the higher energy content, the additional fermentation of black water should be considered for the ICU-concept. For black water usage, the separation of grey and black water is needed. This required a two-pipe system and separation or vacuum toilets (e.g. Jenfelder Au, Gao et al., 2006). The implementation is technically demanding but allows the reuse of greywater, which would further reduce water consumption..

### 3.1.2 Dynamic behavior of the anaerobic digester

For the first simulation a constant supply of biowaste and black water with the chance that fluctuations occur was assumed. To assess the dynamic behavior of the system, after initial conditions (i), a shift in the feeding rate from (ii.) constant 33.6 kg/d to (iii.) an increase of biomass of 18 % for four months (iii.) to (iv.) a feeding rate of 30.7 kg/d for only five days a week was considered. The simulation of all three fermenter types (2sR, CSTR and PFR) shows smooth transitions for the different changes in the feeding strategy indicating an overall stable process behavior. This suggests that the ICU concept could be easily integrated into buildings. However, systems for handling fluctuations in gas production such as gas storage tanks or gas torches should be considered in case of technical problems (data not shown). Furthermore, it is known that ICU systems are prone to long periods of biomass overloading (Bensmann et al. 2016) and require a relatively long period for the start-up.

### 3.1.3. The soiling^®^-process represents an efficient approach to convert digestate to fertilizer

Digestates contain high amounts of NH_3_-N but not NO_3_-N required for plant growth in soilless cultivations. NH_3_ conversion to NO_3_ can be achieved by composting, by the soil microbiome and by nitrification fermenters. One efficient nitrification fermenter is the Soiling^®^-module from Jassen Kunststoffzentrum GmbH (EP 3684909A1). Varification of Jassen GmbH shows that before the soiling^®^-treatment, the NH_3_ amount was 200 mg/l. After 9 days of aeration with 25 m3/d oxygen, the NH_3_ amount decreased to 15 mg/l. This means 92.5 % of the NH_3_-N is nitrified into NO_3_-N (calculation in supplementary file 9). This is superior to other nitrification systems, like the nitrification system of Wang et al., 2017, which has an efficiency of about 87.2 %. Therefore, the soiling^®^-system was used for all further calculations. In addition, the project partner Jassen Kunststoffzentrum GmbH could provide exact numbers for the conversion of the digestate taken from ADP to the fertilizer. Here, a digestate yield of 2.63 l fertilizer per day containing 0.00594 kg/NO_3_-N per liter. In total, this requires 535.7 kJ energy for each kg N. For comparison: the production of fertilizer using artificial nitrogen fixation process (Haber-Bosch process) requires already 10.800 kJ per kg N for the production of NH_4_, which has still to be converted to nitrate. A precondition for the nitrification step is a separation of the liquid and solid components of the digestate. This separation could be achieved by a screw press that is easy to handle and has a low energy consumption (about 0.5 kJ). The remaining solid fraction could also used be upgraded by composting but this was not further considered here. To estimate how much N can be produced as fertilizer from a hydroponic system, the N flow according to the Simba model was considered (Fig. 7).

**Fig. 7:**
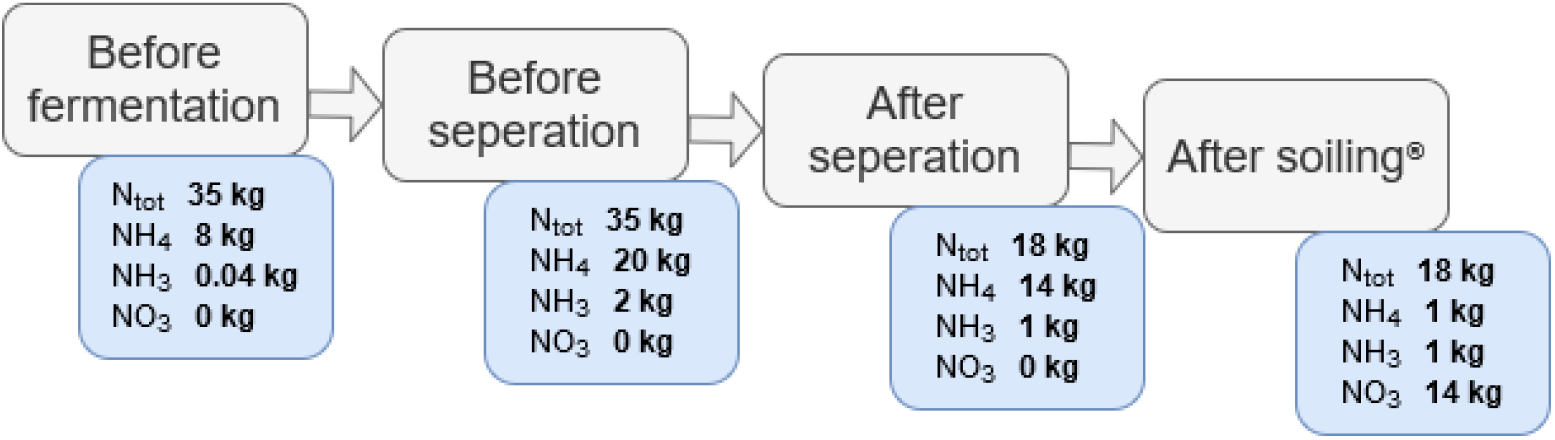
N flow according to the Simba model. N_tot_ = total nitrogen, NH_4_ = ammonium, NH_3_ = ammonia, NO_3_ = nitrate. Quantities relate to one year.

### 3.2.1 Strategies to integrate plant growth into buildings

Our feasibility study considered four scenarios for house-internal gardens (plant production) and compared them regarding productivity, nutrient utilization, energy demand, required skills and social inclusion. For scenario 1 and 2, crop is produced in open roof-top gardens, while protected cultivation in greenhouses or plant factories, also called vertical farming (Carotti et al., 2021), is assumed for scenario 3 and 4. Thus, the scenarios increase from low level to high-level control from scenario 1 to scenario 4 (Fig. 8). Plant factories allow to cultivate crops in multiple layers with a high productivity and uniformity (Graamans et al., 2018). These systems are completely isolated from the exterior climate with the control of light, temperature, relative humidity and CO_2_ concentration (Carotti et al., 2021). Especially by controlling the light quality, the yield and the nutritional value of lettuce can be increased (Cammarisano et al., 2020). However, with increased control from scenario 1 to scenario 4, energy consumption increases through the supply of mechanical heat in greenhouses (Bailey and Seginer, 1989) and electric light in plant factories (Graamans et al., 2018; Harbick and Albright, 2016).

**Fig. 8.**
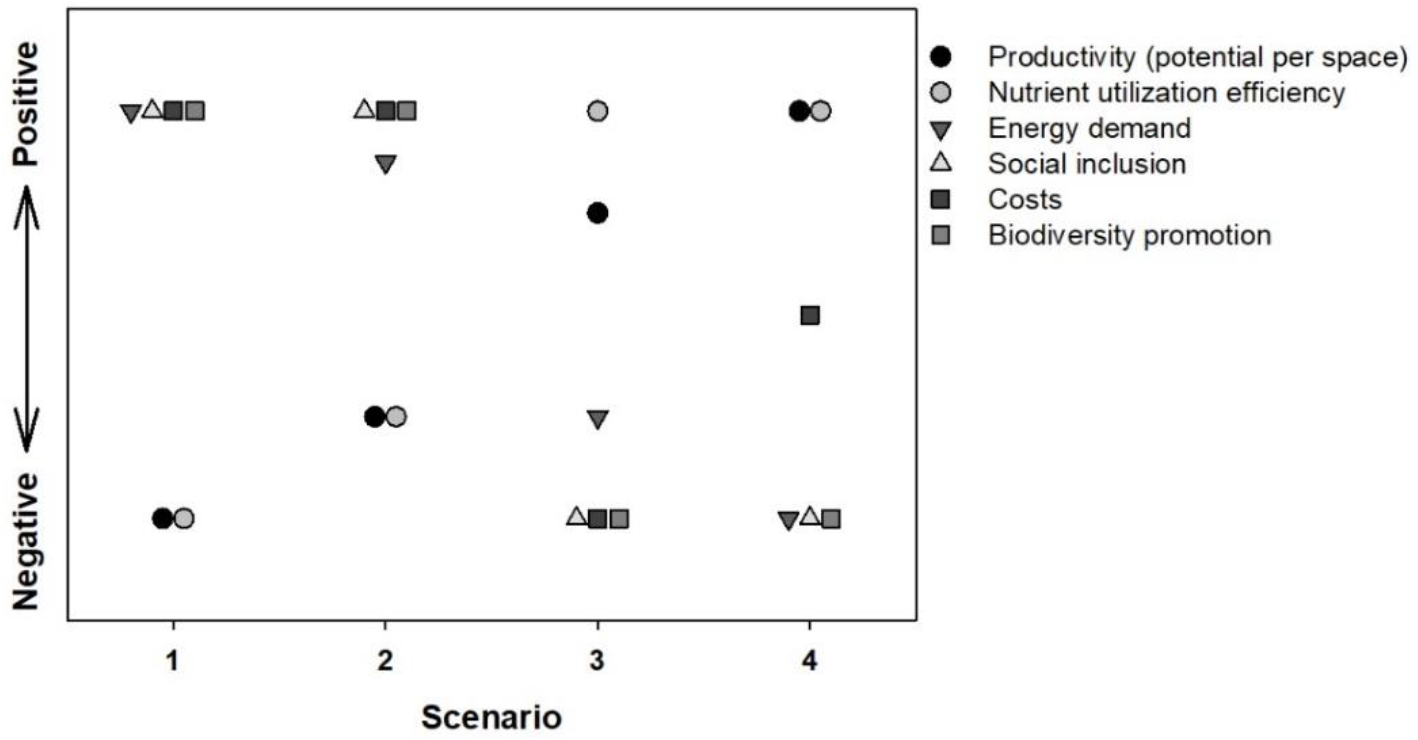
Relative assessment of the cultivation scenarios. Scenario 1: roof-top raised-bed, community residents; Scenario 2: roof-top vertical hydroponic system, community residents; Scenario 3: roof-top greenhouse, professional management; Scenario 4: basement vertical farm, professional management.

**Fig. 9:**
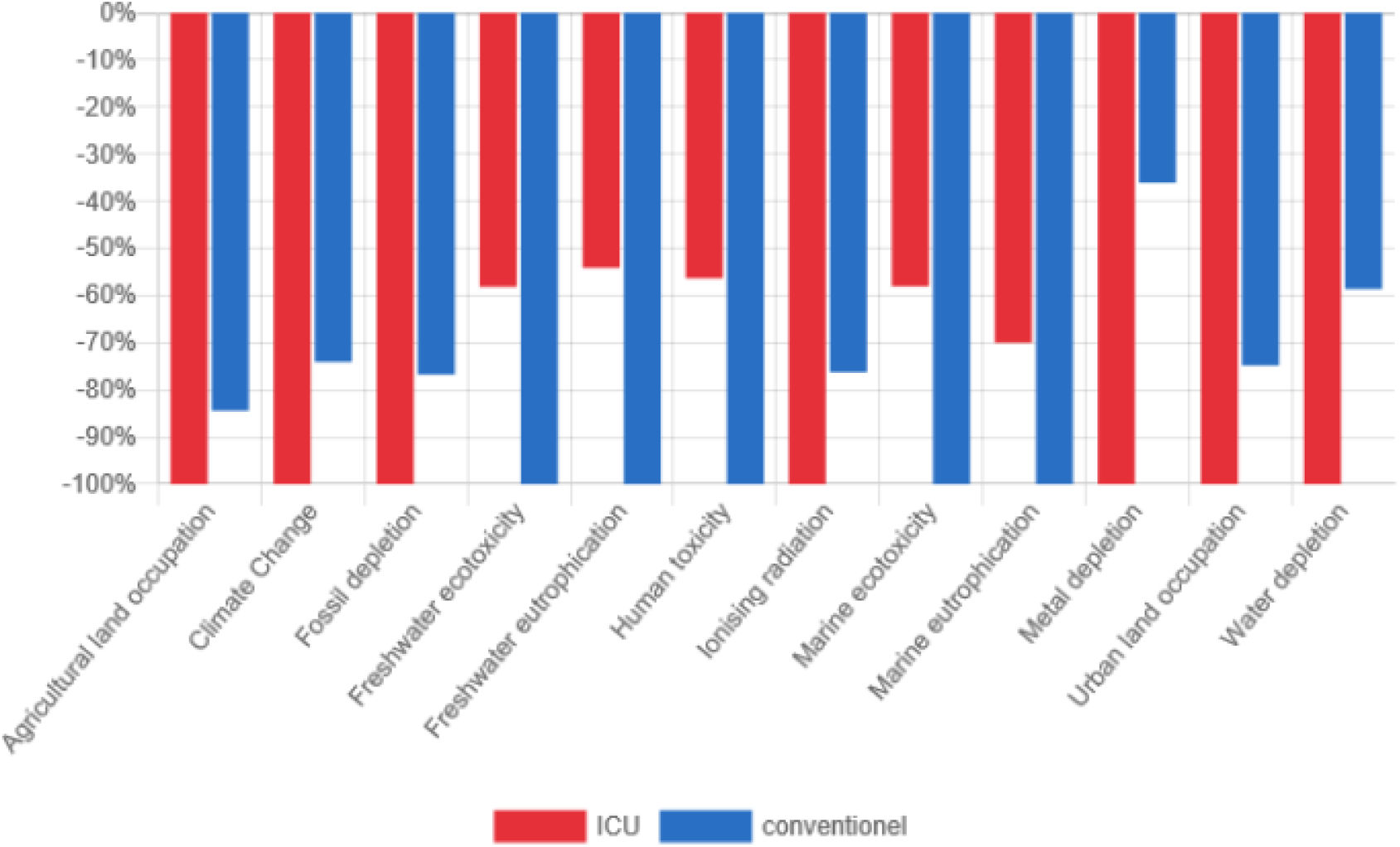
LCA comparison between ICU-concept and conventional building. Diagram shows the impact indicators for ICU and conventional building in %.

In scenario 1, plants are cultivated in raised-beds on the roof-top and maintained by community residents. In this scenario, the residents have a high ecological “feelgood factor”. Maintenance requires only low skills, and the solid fraction of the composted digestate could be utilized, too. While food production is less efficient, a high crop diversity can be attained. With scenario 2, the production efficiency (per m^2^) can be increased. Here, the residents cultivate the plants in vertical hydroponic tiers. Vertical tiers are more space-efficient, while through optimal nutrient availability, hydroponic cultivation systems allow a faster and higher crop production (Rorabaugh et al., 2002). For example, Li et al. (2018) realized in hydroponic systems about twice as much shoot fresh weight of two lettuce cultivars than for the cultivation in culture substrate. However, hydroponic systems require more experience and regular control of the composition of the nutrient solution (Savvas et al., 2008).

Better suited are scenario 3 and 4 because of the controlled or semi-controlled environment as in plant factories or greenhouses. In addition, increased dynamics of the crop water and nutrient demand like in scenario 2 create unfavorable situations with the necessity to discard parts of the nutrient solution (Goddek and Körner, 2019).

In scenario 3, the crop is cultivated in a roof-top greenhouse using hydroponics. This scenario enables very high year-round production but with a higher demand for heating and lighting (see table 3 in section 3.4). The energy consumption is, in particular between autumn and spring, very high. Furthermore a certain dynamic in water and nutrient demand, as well as in yield throughout the year is still present. In scenario 4, this dynamic is more or less completely eliminated. Here, the plant factory is installed in the basement or rooms without natural light and the climate is fully controlled (SharathKumar et al., 2020). Due to the cultivation of crops in layers, even less space than for the roof-top greenhouse is needed. However, since there is no natural light, high amounts of artificial light for plant growth must be provided, and the exhaust heat needs to be removed. Thus, scenario 4 is, despite its higher productivity, a better product quality as well as a stable and constant harvest, the most energy demanding. In addition, it requires maintenance and resources all year round. As the community provides a year-round output in biowaste, continuous use of it can be best achieved with scenario 4. Thus, a decision must be made between the installation of a buffer system for seasonal provision of nutrient-rich irrigation water or a high energy demand. Next to that, non-technical issues also need to be taken into consideration, e.g., the roof-top floor is highly desired by residents and the most expensive floor in the building. In contrast, skyscrapers contain inside rooms without daylight, which must not be used for apartments or offices, at least in Germany.

### 3.2.2 Production potentials and energy demand of simulated scenarios

To evaluate the potential of in-house food production, crop cultivation in protected environments with semi-closed and closed systems, i.e. greenhouses in scenario 3 and plant factories in scenario 4, a model-based simulator tuned to the respective cases was developed. Scenarios 1 and 2 were neglected since their outcome varies, among other factors, highly on the residents’ skills and motivation, which complicates simulations regarding pure bio-technological scenarios significantly. As one of the most common crops in plant factories and greenhouses, lettuce was chosen as a model crop because it is relatively easy to handle during cultivation (Karimaei et al., 2004), and it is suitable to produce lettuce even with wastewater (Sikawa and Yakupitiyage, 2010). Based on the N content in the liquid fraction from the biowaste digestate, it was calculated that the production of 6.3 t fresh mass of lettuce per year would be theoretically possible. A challenge for implementing closed hydroponic systems in the ICU concept is the open question after what time period the fertilizer has to be replaced due to the accumulation of salts (mainly natrium and other deleterious substances. However, in comparison to open systems (the drained nutrient solution is discarded), closed re-circulating hydroponic systems reduce water and fertilizer by about 30 % and 50 %, respectively (Van Os et al., 1999, Grewal et al. 2011).

Using the ICU concept, uncertainties in the dynamic of crop water demand in scenario 3 (through seasonal fluctuations) could result in an unwanted discharge of nutrient solution. This can partly be solved by increasing the aera of the cultivation system. Therefore, the area for the greenhouse in scenario 3 was with 70 m^2^ partly oversized. Due to that, all available N_tot_ will be taken up by the crop. This dynamics is not relevant in scenario 4 and thus a perfect sizing of the plant factory is possible in this case. Accordingly, a total annual yield of 18 000 or 20 500 kg fresh lettuce (i.e. 89.6 kg m^−2^ or 259.37 kg m^−2^; table 1) was predicted for the greenhouse (scenario 3) and the plant factory (scenario 4), respectively. These values correspond to the yields reported in other studies (Becker and Kläring, 2016; Gerbaud and André, 1999; Körner et al., 2018). In these scenarios, a weekly harvest of roughly 2 000 or 3 800 lettuce heads in a 200 m^2^ greenhouse or in a plant factory can be expected. Since 1 kg of lettuce contains about 150 calories and, assuming a person requires about 2 000 calories a day without burden, this would nourish about four people for one year. However, while human nutrition is not covered solely by lettuce, in a real implementation of innovative urban agriculture practitioners a broad mix of vegetables and herbs with various nutritious values should be considered in follow-up studies (Armanda et al. 2019) (Table 2).

**Table 1:**
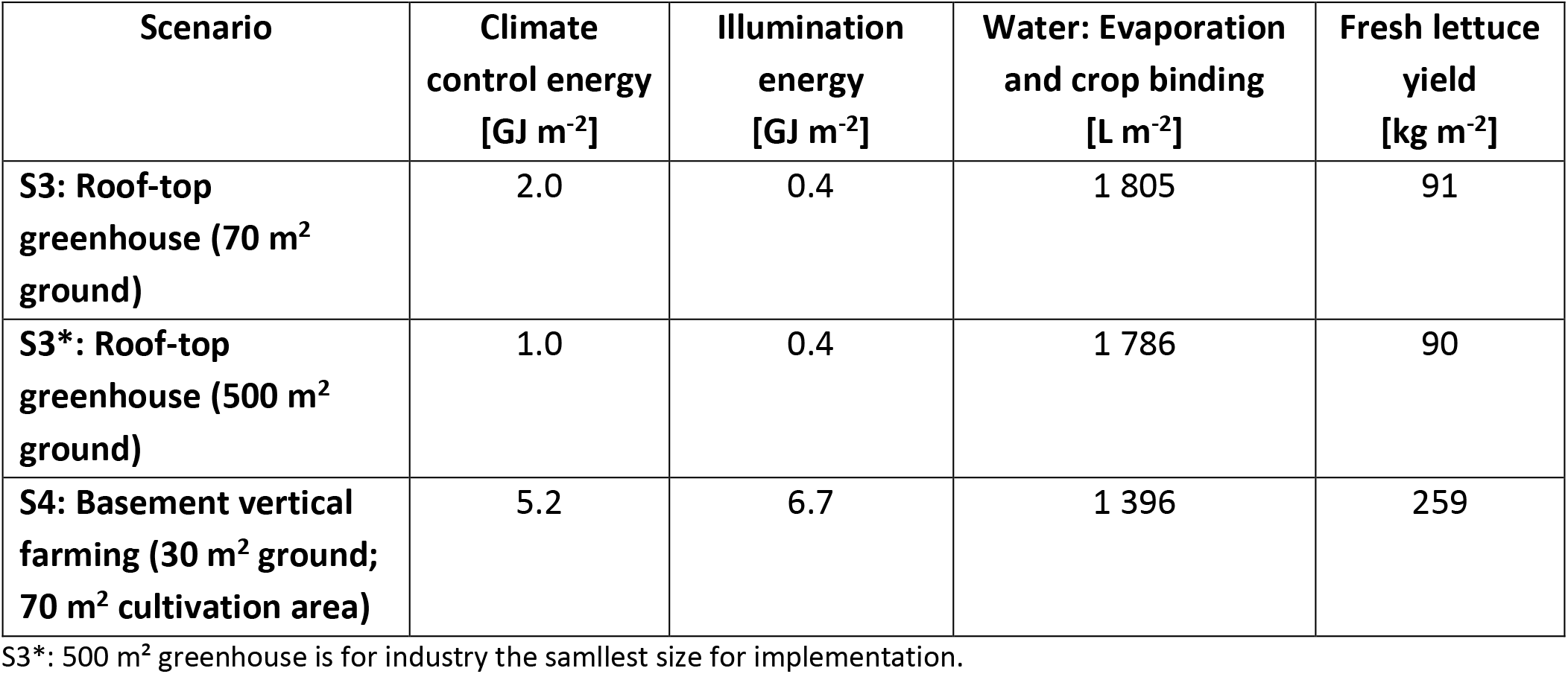
Model output per m^2^ ground area and year

| Scenario | Climate control energy [GJ m<sup>-2</sup>] | Illumination energy [GJ m<sup>-2</sup>] | Water: Evaporation and crop binding [L m<sup>-2</sup>] | Fresh lettuce yield [kg m<sup>-2</sup>] |
| --- | --- | --- | --- | --- |
| **S3: Roof-top greenhouse (70 m<sup>2</sup> ground)** | 2.0 | 0.4 | 1 805 | 91 |
| **S3\*: Roof-top greenhouse (500 m<sup>2</sup> ground)** | 1.0 | 0.4 | 1 786 | 90 |
| **S4: Basement vertical farming (30 m<sup>2</sup> ground; 70 m<sup>2</sup> cultivation area)** | 5.2 | 6.7 | 1 396 | 259 |

S3\*: 500 m<sup>2</sup> greenhouse is for industry the smallest size for implementation.

**Table 2:** Potential biomass production per year for different vegetable crops based on the total available N amount in the liquid effluent and respective calories

| Crop | Potential fresh mass [kg] | Calories [kcal/100 g] |
| --- | --- | --- |
| Broccoli | 3050 | 35* |
| Bush bean | 5490 | 25* |
| Brussel sprouts | 2771 | 43* |
| Lettuce | 7625 | 11** |
| Spinach | 3431 | 23* |

\* https://www.foodspring.ch/magazine/wp-content/uploads/2020/10/FS_Kalorientabelle.pdf
\*\* https://www.wikifit.de/kalorientabelle/gemuese/kopfsalat

**Table 3:**
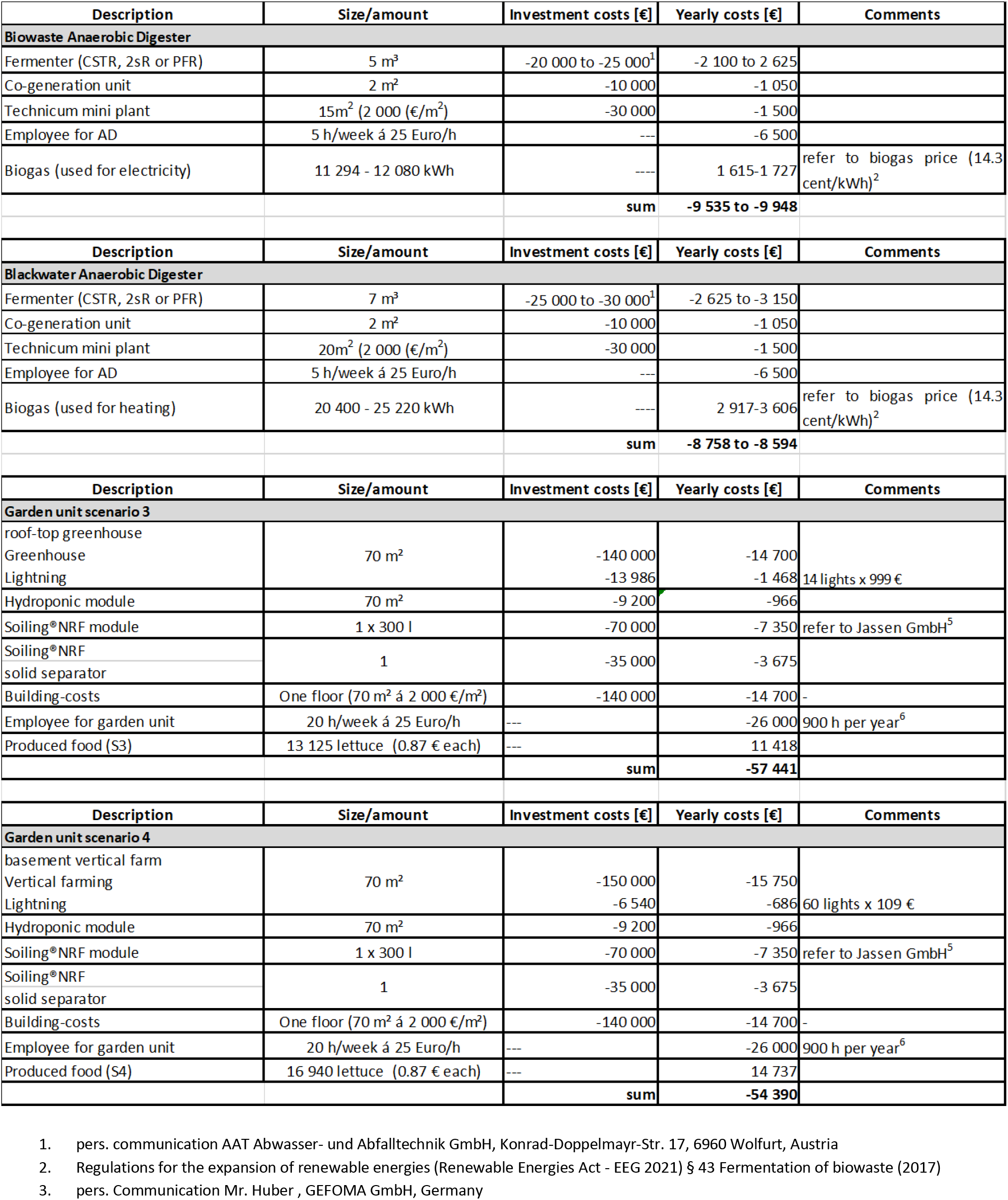
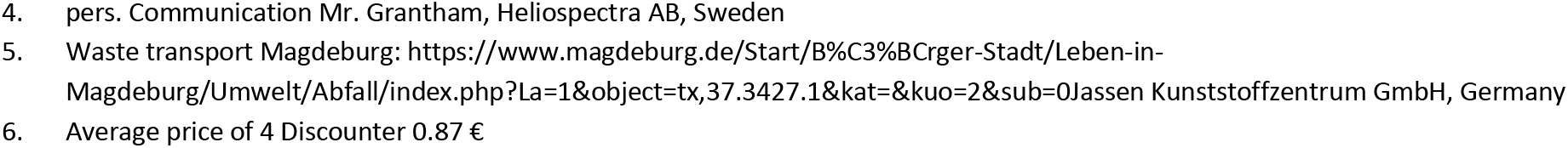
Summary of cost and yields. Yields are considered from the point of the residents using the energy and food for their personal need. References are below table. S3 = scenario 3, S4 = scenario 4.

| Description | Size/amount | Investment costs [€] | Yearly costs [€] | Comments |
| --- | --- | --- | --- | --- |
| **Biowaste Anaerobic Digester** |  |  |  |  |
| Fermenter (CSTR, 2sR or PFR) | 5 m<sup>3</sup> | -20 000 to -25 000<sup>1</sup> | -2 100 to 2 625 |  |
| Co-generation unit | 2 m<sup>2</sup> | -10 000 | -1 050 |  |
| Technicum mini plant | 15m<sup>2</sup> (2 000 (€/m<sup>2</sup>)) | -30 000 | -1 500 |  |
| Employee for AD | 5 h/week á 25 Euro/h | --- | -6 500 |  |
| Biogas (used for electricity) | 11 294 - 12 080 kWh | --- | 1 615-1 727 | refer to biogas price (14.3 cent/kWh)<sup>2</sup> |
|  |  | sum | -9 535 to -9 948 |  |
| **Blackwater Anaerobic Digester** |  |  |  |  |
| Fermenter (CSTR, 2sR or PFR) | 7 m<sup>3</sup> | -25 000 to -30 000<sup>1</sup> | -2 625 to -3 150 |  |
| Co-generation unit | 2 m<sup>2</sup> | -10 000 | -1 050 |  |
| Technicum mini plant | 20m<sup>2</sup> (2 000 (€/m<sup>2</sup>)) | -30 000 | -1 500 |  |
| Employee for AD | 5 h/week á 25 Euro/h | --- | -6 500 |  |
| Biogas (used for heating) | 20 400 - 25 220 kWh | --- | 2 917-3 606 | refer to biogas price (14.3 cent/kWh)<sup>2</sup> |
|  |  | sum | -8 758 to -8 594 |  |
| **Garden unit scenario 3** |  |  |  |  |
| roof-top greenhouse |  |  |  |  |
| Greenhouse | 70 m<sup>2</sup> | -140 000 | -14 700 |  |
| Lightning |  | -13 986 | -1 468 | 14 lights x 999 € |
| Hydroponic module | 70 m<sup>2</sup> | -9 200 | -966 |  |
| Soiling®NRF module | 1 x 300 l | -70 000 | -7 350 | refer to Jassen GmbH<sup>5</sup> |
| Soiling®NRF solid separator | 1 | -35 000 | -3 675 |  |
| Building-costs | One floor (70 m<sup>2</sup> á 2 000 €/m<sup>2</sup>) | -140 000 | -14 700 | - |
| Employee for garden unit | 20 h/week á 25 Euro/h | --- | -26 000 | 900 h per year<sup>6</sup> |
| Produced food (S3) | 13 125 lettuce (0.87 € each) | --- | 11 418 |  |
|  |  | sum | -57 441 |  |
| **Garden unit scenario 4** |  |  |  |  |
| basement vertical farm |  |  |  |  |
| Vertical farming | 70 m<sup>2</sup> | -150 000 | -15 750 |  |
| Lightning |  | -6 540 | -686 | 60 lights x 109 € |
| Hydroponic module | 70 m<sup>2</sup> | -9 200 | -966 |  |
| Soiling®NRF module | 1 x 300 l | -70 000 | -7 350 | refer to Jassen GmbH<sup>5</sup> |
| Soiling®NRF solid separator | 1 | -35 000 | -3 675 |  |
| Building-costs | One floor (70 m<sup>2</sup> á 2 000 €/m<sup>2</sup>) | -140 000 | -14 700 | - |
| Employee for garden unit | 20 h/week á 25 Euro/h | --- | -26 000 | 900 h per year<sup>6</sup> |
| Produced food (S4) | 16 940 lettuce (0.87 € each) | --- | 14 737 |  |
|  |  | sum | -54 390 |  |

- pers. communication AAT Abwasser- und Abfalltechnik GmbH, Konrad-Doppelmayer-Str. 17, 6960 Wolfurt, Austria - Regulations for the expansion of renewable energies (Renewable Energies Act - EEG 2021) § 43 Fermentation of biowaste (2017) - pers. Communication Mr. Huber , GEFOMA GmbH, Germany
4. pers. Communication Mr. Grantham, Heliospectra AB, Sweden 5. Waste transport Magdeburg: https://www.magdeburg.de/Start/B%C3%BCrger-Stadt/Leben-in-Magdeburg/Umwelt/Abfall/index.php?La=1&object=tx,37.3427.1&kat=&sub=0Jassen Kunststoffzentrum GmbH, Germany 6. Average price of 4 Discounter 0.87 €

On the downside: The roof-top greenhouse in scenario 3 (m^2^ ground) requires an annual amount of ~182 GJ (50 550 kWh) energy for heating and 72.5 GJ (20 150 kWh) electrical power for light. While the energy demand is highest during autumn and winter, the yield is highest in summer. Thus, the efficiency of energy use strongly drops in the cold season. Using a cultivation period of March to October requires only about 44 % of the energy for heat and 10 % for light. The latter, however, would required an additional storage solution for crops during that period. Furthermore, power consumption for greenhouse lighting depends on many parameters e.g. geographical location, greenhouse cover light transmission, light source, crop, set points. Comparable results for power demand in greenhouses as in scenario 3 were reported earlier (Aaslyng et al., 2006; Körner et al., 2006; Mortensen and Stromme, 1987; Seginer et al., 2006).

The energy demands for the plant factory case in scenario 4 are even higher. This corresponds to an annual electrical power demand of 201 GJ (56.1 MWh) per year. As each plant factory is unique and large differences exist between implementations, e.g. he number of layers, layer size, empty spacing (room use efficiency), light source, light set point, etc. the power consumption needs to be compared taking into account the net production area and the net installed and used light capacity. Based on our assumptions, the results obtained are in the same range as reported earlier (Kozai, 2013; Tong et al, 2013).

The lettuce crop produced in scenario 4 using a full climate-controlled plant factory consumes 95 % of the fertilizer of the ICU system when a (nearly) closed system is attained. About 36 m^3^ water would be additionally needed if the evaporation water is not reused.

As both scenario 3 and 4 have a high energy demand, these solutions are not the best concerning the CO_2_ footprint. However, to make controlled hydroponic systems more sustainable, regenerative energy could be used more intensively in this culture system, as it is already done for some urban farms (Armanda et al. 2019). Moreover, due to further advantages, like reduced water consumption and space requirements in comparison to a field cultivation, less transport and utilization of non-used spaces in the city, reliable and stable resource demand and crop yield, higher quality and food safety, there is a great potential for controlled and semi-controlled crop production in an urban scenario. Additionally, local food production is highly advantageous. In particular, it represents an alternative for highly-populated mega cities lacking space or for regions lacking agricultural areas.

To stress the full potential of the ICU concept, further extrapolations were performed. The complete fertilizer produced from the biowaste (N_tot_ in liquid and solid fraction neglecting possible N-losses during further processing like composting and/or immobilisation processes) allows for the cultivation of about 19.5 t lettuce. Interestingly, the 720 l of black water produced by the residents already contain 183.95 kg NH_4_, which could be converted to 170.15 kg NO_3_. This amount per year could theoretically allow to produce 1.1 t/ m^2^ lettuce in a roof-top greenhouse, and clearly demonstrates the potential of reusing nutrients from the digestate of biowaste and black water to produce food and nourish urban populations.

### 3.3 Implementation of a ICU concept for a building with 100 residents saves up to 6,468 kg CO_2_-eq

Implementation of the ICU project for a building with 100 residents reduced CO_2_-emission due to reduced transport of 11 tons biowaste by 693 kg CO_2_. The reuse of NH_4_ as fertilizer saved 2 363 kg CO_2_ compared to the new synthesis by the Haber-Bosch process. Usage of the produced 25 855 m^3^ biogas for heating saved 3 412 kg CO_2_ and is comparable to the CO_2_ fingerprint for heating reported in literature (Capponi et al., 2012). In total, the implementation of the ICU concept can save 6,468 kg CO_2_-eq. Based on a CO_2_ emission price of 25 € per ton CO_2_, the CO_2_ saving value is currently 161.7 € (BMU, 2021) (supplementary fig. 9).

### 3.4. ICU concept becomes economically feasible in large buildings and with growing food prices

To estimate the economic feasibility, yearly costs and yields of the ICU concept were estimated. Operation of an AD for biogenic waste (table 3, Biowaste Anaerobic Digester) costs 9 535–9 948 € and yields 1 615-1 727 € annually. In contrast, operation of an AD for blackwater utilization costs 8 594-8 758 € and yields 2 917-3 606 € annually. In consequence, the in-house use of biowaste and black water is not profitable for small buildings (Salerno et al., 2017). For large buildings, however, personal costs remain more or less the same and investment costs for larger fermenters rise only slightly. Therefore, implementation is economically feasible for large buildings or agglomeration of several buildings, favoring the ICU implementation in large cities or at the district level.

Generally, black water utilization is economically more promising than biowaste usage under the premise that the saved cost for the wastewater removal compensates for the black water system’s cost. Combined usage of black water and biowaste would create the synergy that personal costs for the daily lookup and the co-generation unit can be shared.

The annual cost for a 70 m^2^ greenhouse on the roof-top was 57 441 € (scenario 3), and in the basement 54 390 € (scenario 4). The benefit of both solutions is 11 418 – 14 737 €, based on a lettuce price of 0.87 €. Extrapolation showed that the system is profitable with lettuce prices of 1.80 €. This rise is possible when the agricultural space vanishes further, and the population grows. For example, in Singapore, the lettuce price is already 1.00 −2.50 €. In contrast to the AD, the hydroponics’ economic efficiency grows only slightly with larger systems since it scales quite well.

Whether the hydroponic should be integrated on the roof-top, in the basement, or in rooms without light depends heavily on the price. The top floor’s rental price is roughly 30 % more expensive. So, it could bring more profit to use the top floor as a penthouse. In a prestigious skyscraper with around 100 m^2^ of living space, a six-digit amount in major German cities is possible.

### 3.5 High motivation of stakeholders for the implementation of an ICU concept but high legal barriers

#### 3.5.1 High motivation of residents for a sustainable lifestyle

Current studies show that in Germany, a sustainable lifestyle becomes more and more important (Tölkes et al. 2018). Today, 657 urban gardening projects exist in Germany (Winkler et al., 2019). However, the engagement of residents in urbane gardening might decline and require strategies to counteract. In contrast, a professional management of the urban gardening projects is more reliable but often reduces the acceptance of residents (Specht et al. 2016). A compromise could be combined models with hydroponic modules for the residents or a botanic garden with a cafe alongside professionally managed greenhouses.

#### 3.5.2 Real estate owners would implement the ICU project as long as it is profitable

Almost all real estate owners participating in this survey considered sustainable building as important for their business. Nevertheless, closed material cycles or exploitation of options related to the operation of AD or plant cultivation seemed less important for most of them. Some may even lose their tax benefits when they engage in another business field like urbane agriculture or operation of an AD. Therefore, it seems beneficial to outsource the operation of an ICU project to a contracting partner or a cooperative of the residents. Necessary conditions for implementing ICU concepts for real estate owners are high acceptance of the residents, lower maintenance requirements and profitability. Typically, there is no interest in pure flagship projects. Furthermore, especially for the roof-top use of buildings, ICU projects compete with other more established sustainable solutions such as the operation of photovoltaic systems (supplementary file 8).

#### 3.5.3 In the framework of the Paris agreements, government promote a CO_2_-society

In the Paris Agreement, 195 states including the European Union agreed to limit global warming below 2° C (www.unfccc.int/process-and-meetings/the-paris-agreement/the-paris-agreement). Therefore, netto CO_2_ emission has to be reduced to zero by 2040. In addition to the targets for the energy and building sector, new goals and measures are also being set for agriculture. In Germany, 8 % (72 million tons of CO_2_-eq) of the greenhouse gas emission came from agriculture in 2014. Despite the fact that of most agricultural greenhouse gas emissions are caused by natural physiological processes, the ability to reduce them is limited. The largest source with about 25 million tons of CO_2_-eq is the use of N fertilizers. The use of organic nutrients considered in the ICU project, in particular the soiling^®^-products, are one important step towards minimization of the N use. The main goal until 2030 is to significantly reduce the emissions of mineralic fertilizers in agriculture. One option is to use financing instruments under the Common Agricultural Policy. Another option is to increase the percentage of land used for organic farming by circular economy approaches (Klimaschutzplan 2050: https://www.bmu.de/publikation/klimaschutzplan-2050/).

#### 3.5.4 ICU implementations must fulfill high legal standards favoring large projects or tiny ones for personal need

The implementation of ICU projects requires observing the laws of the state. In the following, the most critical regulations for implementation ICU concepts in Germany are exemplarily addressed. One important factor is to fulfill the guidelines for building security (BauGB §29-§38, in particular admissibility of projects §34, BauNVO). Additionally, urban development plans and urban planning law must be complied with. Biogas production using AD for biowaste and black water is critical since biogas is flammable and requires sufficient ventilation. Therefore, the storage of large biogas volumes should be avoided. The produced biogas should be consumed immediately, upgraded to natural gas, fed in the gas grid, or outsourced from the buildings (Schmidt-Eichstaedt, 2019). Nevertheless, fire prevention (§§ 3 and 14 MBO) and explosion control (DGUV Regel 113-001, Frigger et al., 2019) must be considered. For the removal of black water digestate and the solid fraction of the biowaste fermentation, the laws for sludge disposal have to be considered (AbfKlärV, Queitsch et al., 2918). Utilization of the biowaste as fertilizer requires compliance with the German laws for biowaste (Bioabfallverordnung (BioAbfV)) and fertilizer ordinance (DüMV) as well as the EU regulations for fertilizer ordinance (EU-FPR). In particular, base materials must be allowed (DüMV, supplementary 2, table 7). The fertilizer has to be listed in a positive list (DüMV, supplementary 1, table 1) or equals any allowed fertilizer type. Furthermore, emission limits (DüMV, supplementary 2, table 1.4) and minimum hygiene requirements (§ 5 DüMV) have to be fulfilled. Exception exist in the case the biowaste and the produced fertilizer are only used for personal needs. However, it is questionable if biowaste utilization in a cooperative of more than 100 residents accounts as a personal need. Due to the high legal requirementst, implementationof ICU concepts seems only manageable for large projects or tiny implementations for personal needs (supplementary file 3). Another question is liability, which is difficult to address in general and usually depends on the specific case. Therefore no general recommendation can be given here, except to address this issue in a contract between the stakeholders (supplementary file 6).

#### 3.5.5 Communication and participation are important for the acceptance of residents

Implementation of ICU projects require the participation of the residents. Residents have to separate the biowaste accurately, agree to install vacuum toilets and use the urban gardens either as gardeners or as consumers. In general, there is a high acceptance in Germany to waste separation (Walk et al., 2019), vacuum toilets (Poortvliet et al., 2018), and urban gardening (Winkler et al., 2019). However, it is always useful to integrate all stakeholders as early as possible to successfully implement projects (García-Sánchez et al., 2018) and, in addition, to guide their participation by teaching material.

Furthermore, it is recommended to communicate potential risks (Xia et al., 2018). For example, ICU operation has the risk of microbial contamination of the food collected. Pretretment of biomass at 70 °C can ensure the inactivation of harmful microorganisms. Another issue is the produced biogas, which is explosive. However, when immediately consumed, the risk is reduced to the level of a conventual gas heater.

An important cultural aspect is the utilization of black water as fertilizer. Theoretically, animal dung or manure usage and spreading it on fields are quite similar to the use of black water digestate for hydroponics. However, this is neither allowed nor accepted (Gell et al., 2011).

### 3.6 Strategies to extend the ICU concept

The ultimate goal of the ICU project is to close energy and material flow cycles in an urbane building. Additional components could increase yields and productivity and allow for a more robust operation.

For example, hydroponic modules for the balcony could be added, or food production can be elevated by aquaponic (Chia et al., 2018) or algae cultivation (Wongkiew et al., 2017). For an implementation, strategies to combine agriculture with photovoltaic could also be implemented (Navarte et al., 2018, Putri et al., 2018). For hot and dry areas, the ICU concept could be extended to include the water cycle (Liuzzo, 2016). For example, greywater can be reused (Hertel et al., 2015) or rainwater could be used for adiabatic cooling. In order to achieve a more robust operation, a module for cleaning the biowaste, for example, via conveyor belts, can be added (Verma et al., 2002). Also, storage capacities for biowaste, biogas, or fertilizers can be added. However, additional stores in the building are expensive and increase the fire load.

## 4. Conclusions

Integrating biomass cycles into residential buildings, as proposed by the ICU concept, is technically feasible, reduces CO_2_ emission, and is of interest to owners of urband buildings and their residents. It is profitable for implementation in large buildings or agglomeration of buildings and in case food prices further increase. However, to achieve this goal, it will require the implementation of prototypes to perfect technical details and to confirm economic and material calculations. Major challenges for the implementation come from legal aspects relate to the biowaste prescription (the German BioAbfV) and the fertilizer prescription (the German DüMV). In sum, the results of this study should bring us one step closer to a reduction in land use and to a sustainable, CO_2_-neutral society.

## Abbreviations

AD: Anaerobic digestion
ADM1: Anaerobic digestion model number one
ADM1da: Anaerobic digestion model number one da
ADP: Anaerobic digestion plant
BioAbfV: Bioabfallverordnung
CCPP: Combined cycle power plants
CHP: Combined heat and power
CH_4_: Methane
CO_2_: Carbon dioxide
CO_2_-eq: Carbon dioxide equivalent
CSTR: Continuous stirred-tank reactor
DLI: Daily light integral
DüngV: Düngemittelverordnung
EC: Electrical conductivity
ICU: Integrated Cycles for Urban Biomass
K: Potassium
LAT: Latidude
LCA: Life cycle assessment
LCCA: Life cycle cost analysis
LON: Longitude
MT: Microturbines
N: NitrogenNH_4_
NH_3_: Ammonia
NO_3_: Nitrate
NO_3_-N: Nitrate nitrogen
NO_2_: Nitrite
NPK: fertilizer Nitrogen, phosphate and potassium fertilizer
NPV: Net present value
N_tot_: Total nitrogen
P: Phosphorus
PFR: Plug flow reactor

## 6. Supplementary

File 1: Fluctuation of biomass

File 2: Simba parameter

File 3:Matlab parameter for crop model

File 4: openLCA parameters

File 5: Online questionnaire

File 6: Calculation for optimal fermenter size for hydrolysis

File 7: Results online questionnaire

File 8: Legal aspects

## 7. Acknowledgement

The authors acknowledge the financial support by a funding program of the Fachagentur Nachwachsende Rohstoffe e.V. (FNR, FPNR). We additionally thank the GEFOMA GmbH for the estimation of costs of greenhouse construction. Jennifer Uebbing received funding from the EU-program ERDF (European Regional Development Fund) of the German Federal State Saxony Anhalt by Research Center of Dynamic Systems (CDS).

