## Supplementary file 1 for "Integrated cycles for urban biomass as a strategy to promote a CO_2_-neutral society – a feasibility study"

Supplementary Table XX. Parameter values and their units as used in the two crop simulation model scenario in the greenhouse and plant factory set-up

| Description | Parameter | Unit | Value | Scenario |
|---|---|---|---|---|
| **Crop** | | | | |
| Fraction dry matter harvested product | $\Theta_d$ | - | 0.048 | 3,4 |
| Target harvest fresh weight | $FW_t$ | g $plant^{-1}$ | 100 | 3,4 |
| plant density | $\rho_p$ | - | 36 | 3,4 |
| Maximum leaf area index | $LAI_{max}$ | - | 1.6 | 3,4 |
| Superficial chlorophyll density | $\rho_{chl}$ | - | 0.45 | 3,4 |
| Fraction PAR absorbed by non-photosynthetic parts | $\Theta_{PAR}$ | - | 0.1 | 3,4 |
| **Climate** | | | | |
| Setpoint minimum daily light integral greenhouse | $DLI_{set,GH}$ | mol $m^{-2}$ $d^{-1}$ | 12 | 3 |
| Setpoint minimum daily light integral plant factory | $DLI_{set,VF}$ | mol $m^{-2}$ $d^{-1}$ | 20 | 4 |
| Setpoint temperature heating day | $T_{h,d}$ | °C | 18 | 3,4 |
| Setpoint temperature heating night | $T_{h,n}$ | °C | 16 | 3,4 |
| Setpoint temperature ventilation day | $T_{v,d}$ | °C | 22 | 3 |
| Setpoint temperature ventilation night | $T_{v,n}$ | °C | 22 | 3 |
| Setpoint temperature cooling | $T_c$ | °C | 18.5 | 4 |
| Setpoint maximum humidity | $RH_s$ | % | 82.5 | 3,4 |
| Setpoint daytime minimum $CO_2$ | $C_s$ | µmol $mol^{-1}$ | 700 | 3,4 |

| | | | | |
|---|---|---|---|---|
| **Cultivation** | | | | |
| LED light energy conversion | I | mol $J^{-1}$ | 2.4 | 3,4 |
| Fraction diffuse light transmission greenhouse | $\Theta_{I,Tr}$ | - | 0.8 | 3 |
| Net crop production area greenhouse | $A_{crop,GH}$ | $m^2$ | 70 | 3 |
| Net crop production area plant factory | $A_{crop, VF}$ | $m^2$ | 70 | 4 |
| Heights vertical production layer | $h_{VF}$ | m | 0.5 | 4 |
| Number vertical production layers | $N_{VF}$ | - | 4 | 4 |
| Total volume production unit greenhouse | $V_{GH}$ | m3 | 280 | 3 |
| Total volume production unit plant factory | $V_{VF}$ | m3 | 60 | 4 |
