## Supplementary file 2 for "Integrated cycles for urban biomass as a strategy to promote a CO_2_-neutral society – a feasibility study"

### FRAGEBOGEN Akzeptanzanalyse ICU

Hintergrund:

Modelle zur Produktion von Strom/Wärme und Nutzpflanzen:

A) Nutzung von Biomasse aus dem Haushalt/ Garten,
B) Nutzung von Schwarzwasser,
C) Produktion von Nutzpflanzen in einem Gewächshaus?

Zielgruppe: Wohnungsbaugesellschaften/Hauseigentümer

Verteilung per E-Mail über Interessenvertretungen Baubranche / Multiplikatoren

Dauer: 15 Minuten

Ablauf: 31.08. Fragebogen fertig, Partner eine Woche Zeit Feedback, dann ab 07-11.09. für vier Wochen online stellen

Ziel: Interesse und Hindernisse der Wohnungseigentümer für die Installation einer solchen Anlage herausfinden

Fragen zu

- Einstellung/Interesse des Unternehmens allgemein
- Welche Merkmale/Hindernisse ein akzeptables System haben müsste
- Fragen zum Unternehmen

Tool: Microsoft Forms

Arbeitsteilung:

- Anschreiben und Adressaten > Anita
- Fragenkatalog und digitale Umsetzung > Regine

Weitere Beispielprojekt:

- Arbeitsagentur Oberhausen
- Roof Water Farm

FRAGEBOGEN

#### Integration von Biogasnutzung und Pflanzenproduktion in Wohngebäuden

##### A) Bedeutung und Interesse

Bitte schätzen Sie ein, inwiefern die folgenden Aussagen für Ihr Unternehmen und Ihre Gebäude zutreffen:

| | Sehr groß | groß | gering | Sehr gering | Keine Angabe |
|---|---|---|---|---|---|
| Welche Bedeutung hat „nachhaltiges“ bzw. „grünes“ Bauen für Ihr Unternehmen? | | | | | |
| Welche Bedeutung hat in diesem Zusammenhang bisher die Optimierung/das | | | | | |

| |
|---|
| Schließen von Stoffströmen in Ihren Gebäuden? |
| Wie groß ist das Interesse in Ihrem Unternehmen an der Möglichkeit, das eigene Geschäftsfeld um Biogasproduktion und Gartenbau zu erweitern? |

| Kommentar: |
|---|

#### B) Merkmale der Anlage

Bitte schätzen Sie ein, welche Merkmale ein für Ihr Unternehmen akzeptables System aus Ihrer Sicht haben müsste:

| | Sehr wichtig | wichtig | Nicht so wichtig | Gar nicht wichtig | Keine Angabe |
|---|---|---|---|---|---|
| Geringer Wartungsaufwand des Betriebs | | | | | |
| Einfache Handhabung der Anlagen | | | | | |
| Hohe visuelle Sichtbarkeit der Anlage (z.B. großes Gewächshaus) | | | | | |
| Keine betriebsbedingten Beeinträchtigungen der MieterInnen (Geruch, Verkehr,…) | | | | | |
| Akzeptanz der MieterInnen (z.B. Mülltrennung) | | | | | |
| Möglichst kurzer Zeitraum bis zur Amortisierung | | | | | |
| Große Außenwirkung als Vorzeigeprojekt (z.B. Auszeichnungen, internationale Presse) | | | | | |

| Kommentar: |
|---|

#### C) Fragen zum Unternehmen

- Bitte geben Sie an, wieviele Wohneinheiten Ihr Unternehmen besitzt:
- In welchen Bundesländern ist Ihr Unternehmen aktiv? (Mehrfachauswahl + Kategorie Bundesweit + Kategorie International)
- Wo befinden sich Ihre Gebäude in Deutschland mehrheitlich: Innenstadt, Stadtrand oder ländlichen Bereich? /unklar
- Planen Sie für die kommenden 5 Jahre weitere Wohnungsbauprojekte in Deutschland? Ja/nein/unklar

- Spielt die Produktion von erneuerbare Energien in Ihren jetzigen und geplanten Gebäuden bereits eine Rolle? Ja/nein/teilweise
- Haben Sie Interesse, an einem Pilotprojekt zur gebäudegebundenen Produktion von Biogas und Nutzpflanzen teilzunehmen? ja/nein
    - Falls Ja > geben Sie hier bitte Ihren Kontakt ein: (Unternehmen, Ansprechperson, email, Webseite)

| Kommentar: |
| --- |
