## Supplementary file 4 for "Integrated cycles for urban biomass as a strategy to promote a CO_2_-neutral society – a feasibility study"

### Slide 1
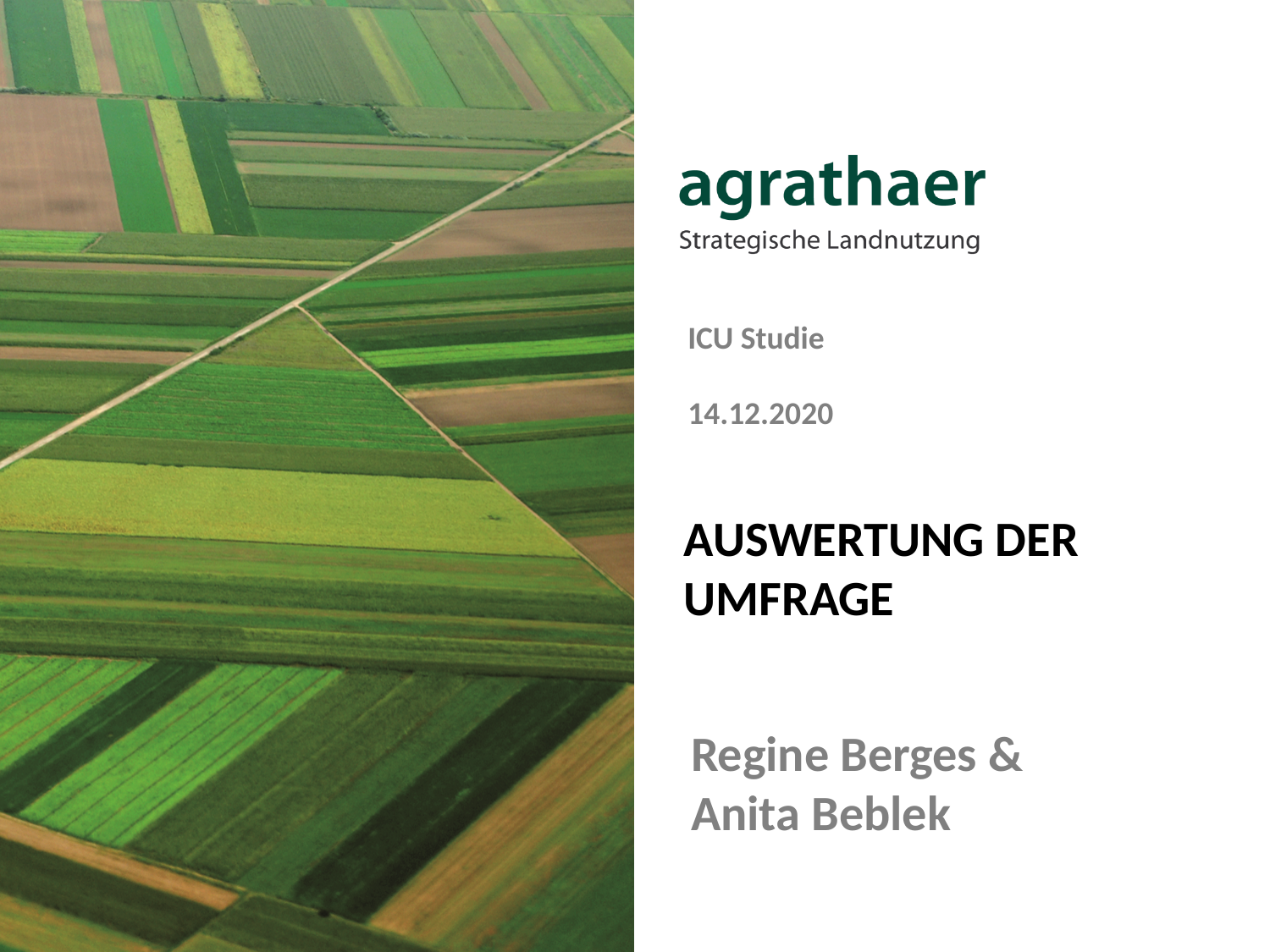

ICU Studie
14.12.2020
#### Auswertung der Umfrage
Regine Berges & Anita Beblek

### Slide 2
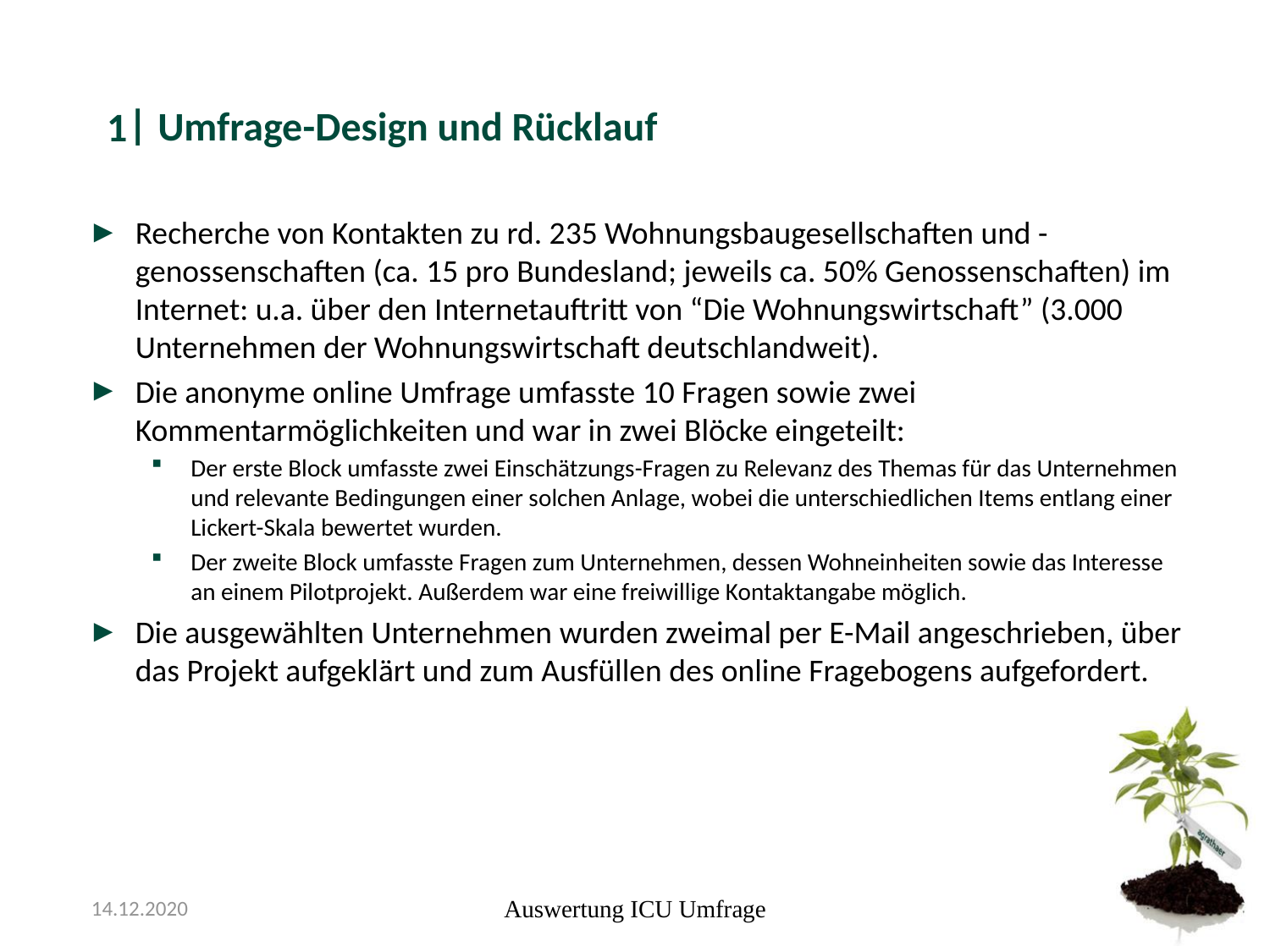

#### Umfrage-Design und Rücklauf
1
Recherche von Kontakten zu rd. 235 Wohnungsbaugesellschaften und -genossenschaften (ca. 15 pro Bundesland; jeweils ca. 50% Genossenschaften) im Internet: u.a. über den Internetauftritt von “Die Wohnungswirtschaft” (3.000 Unternehmen der Wohnungswirtschaft deutschlandweit).
Die anonyme online Umfrage umfasste 10 Fragen sowie zwei Kommentarmöglichkeiten und war in zwei Blöcke eingeteilt:
Der erste Block umfasste zwei Einschätzungs-Fragen zu Relevanz des Themas für das Unternehmen und relevante Bedingungen einer solchen Anlage, wobei die unterschiedlichen Items entlang einer Lickert-Skala bewertet wurden.
Der zweite Block umfasste Fragen zum Unternehmen, dessen Wohneinheiten sowie das Interesse an einem Pilotprojekt. Außerdem war eine freiwillige Kontaktangabe möglich.
Die ausgewählten Unternehmen wurden zweimal per E-Mail angeschrieben, über das Projekt aufgeklärt und zum Ausfüllen des online Fragebogens aufgefordert.
14.12.2020
Auswertung ICU Umfrage

### Slide 3
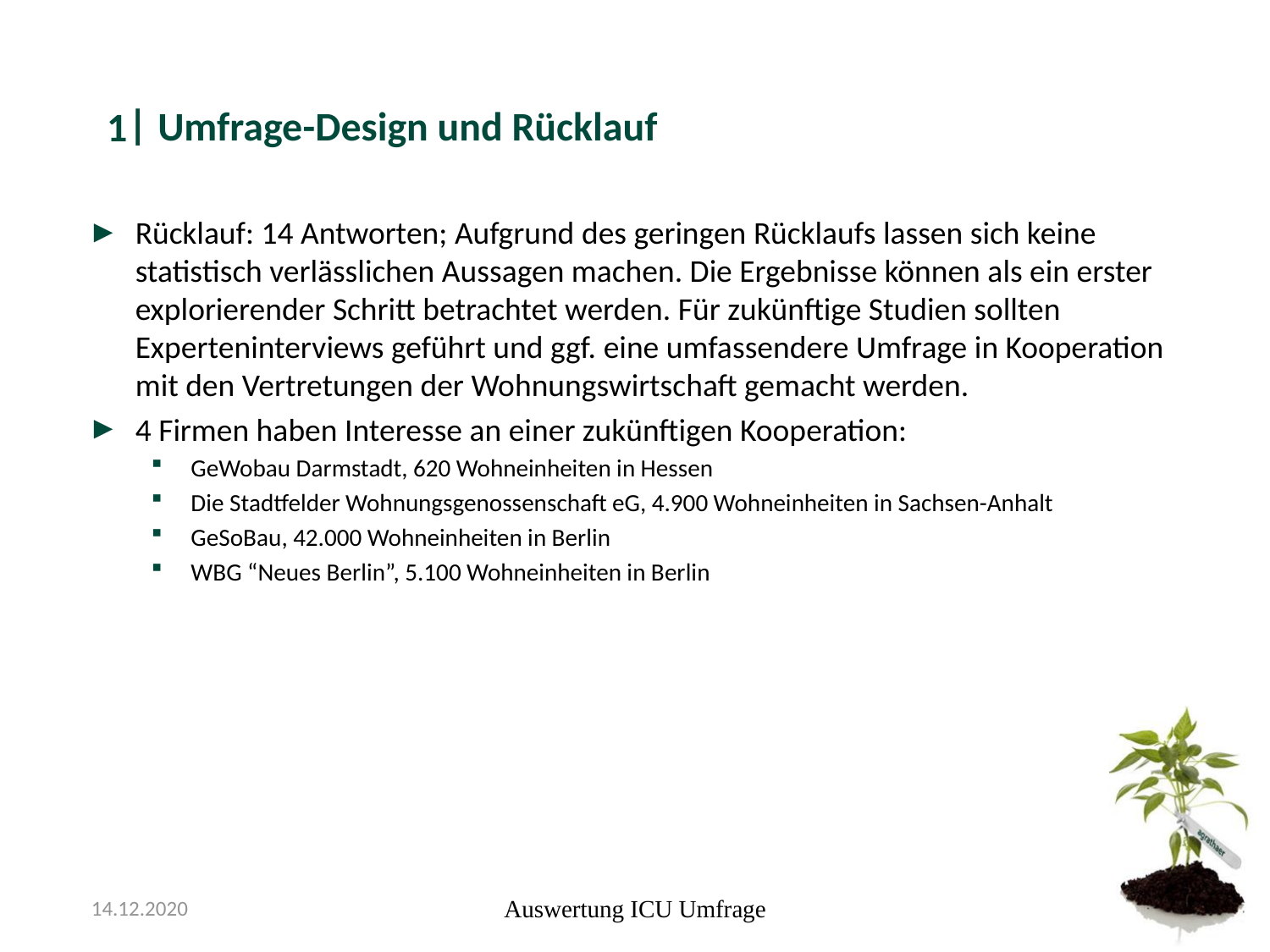

#### Umfrage-Design und Rücklauf
1
Rücklauf: 14 Antworten; Aufgrund des geringen Rücklaufs lassen sich keine statistisch verlässlichen Aussagen machen. Die Ergebnisse können als ein erster explorierender Schritt betrachtet werden. Für zukünftige Studien sollten Experteninterviews geführt und ggf. eine umfassendere Umfrage in Kooperation mit den Vertretungen der Wohnungswirtschaft gemacht werden.
4 Firmen haben Interesse an einer zukünftigen Kooperation:
GeWobau Darmstadt, 620 Wohneinheiten in Hessen
Die Stadtfelder Wohnungsgenossenschaft eG, 4.900 Wohneinheiten in Sachsen-Anhalt
GeSoBau, 42.000 Wohneinheiten in Berlin
WBG “Neues Berlin”, 5.100 Wohneinheiten in Berlin
14.12.2020
Auswertung ICU Umfrage

### Slide 4
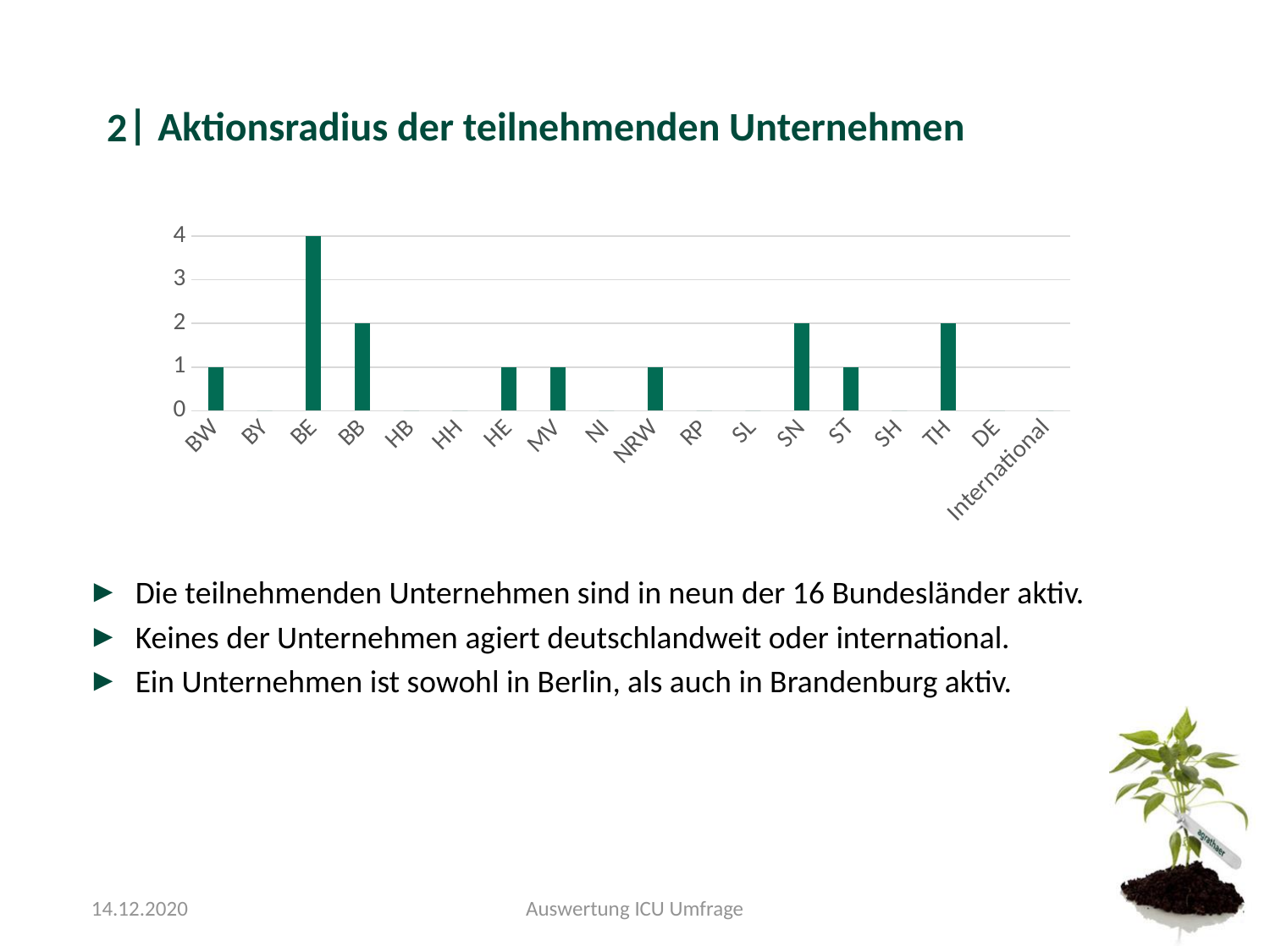

#### Aktionsradius der teilnehmenden Unternehmen
2
###### Chart
| Category | |
|---|---|
| BW | 1.0 |
| BY | 0.0 |
| BE | 4.0 |
| BB | 2.0 |
| HB | 0.0 |
| HH | 0.0 |
| HE | 1.0 |
| MV | 1.0 |
| NI | 0.0 |
| NRW | 1.0 |
| RP | 0.0 |
| SL | 0.0 |
| SN | 2.0 |
| ST | 1.0 |
| SH | 0.0 |
| TH | 2.0 |
| DE | 0.0 |
| International | 0.0 |Die teilnehmenden Unternehmen sind in neun der 16 Bundesländer aktiv.
Keines der Unternehmen agiert deutschlandweit oder international.
Ein Unternehmen ist sowohl in Berlin, als auch in Brandenburg aktiv.
14.12.2020
Auswertung ICU Umfrage

### Slide 5
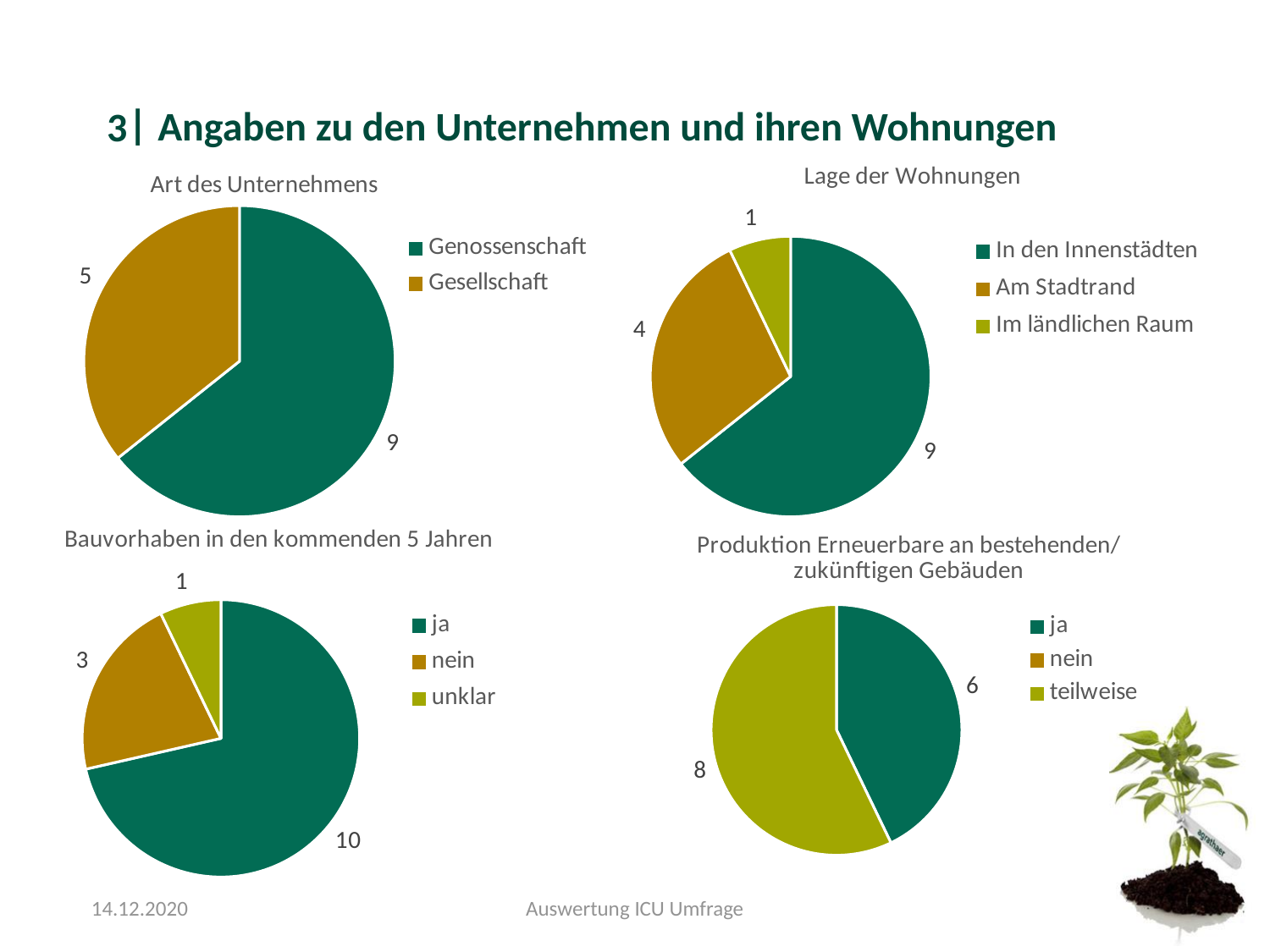

#### Angaben zu den Unternehmen und ihren Wohnungen
3
###### Chart: Art des Unternehmens
| Category | |
|---|---|
| Genossenschaft | 9.0 |
| Gesellschaft | 5.0 |
###### Chart: Lage der Wohnungen
| Category | |
|---|---|
| In den Innenstädten | 9.0 |
| Am Stadtrand | 4.0 |
| Im ländlichen Raum | 1.0 |
###### Chart: Bauvorhaben in den kommenden 5 Jahren
| Category | |
|---|---|
| ja | 10.0 |
| nein | 3.0 |
| unklar | 1.0 |
###### Chart: Produktion Erneuerbare an bestehenden/ zukünftigen Gebäuden
| Category | |
|---|---|
| ja | 6.0 |
| nein | 0.0 |
| teilweise | 8.0 |14.12.2020
Auswertung ICU Umfrage

### Slide 6
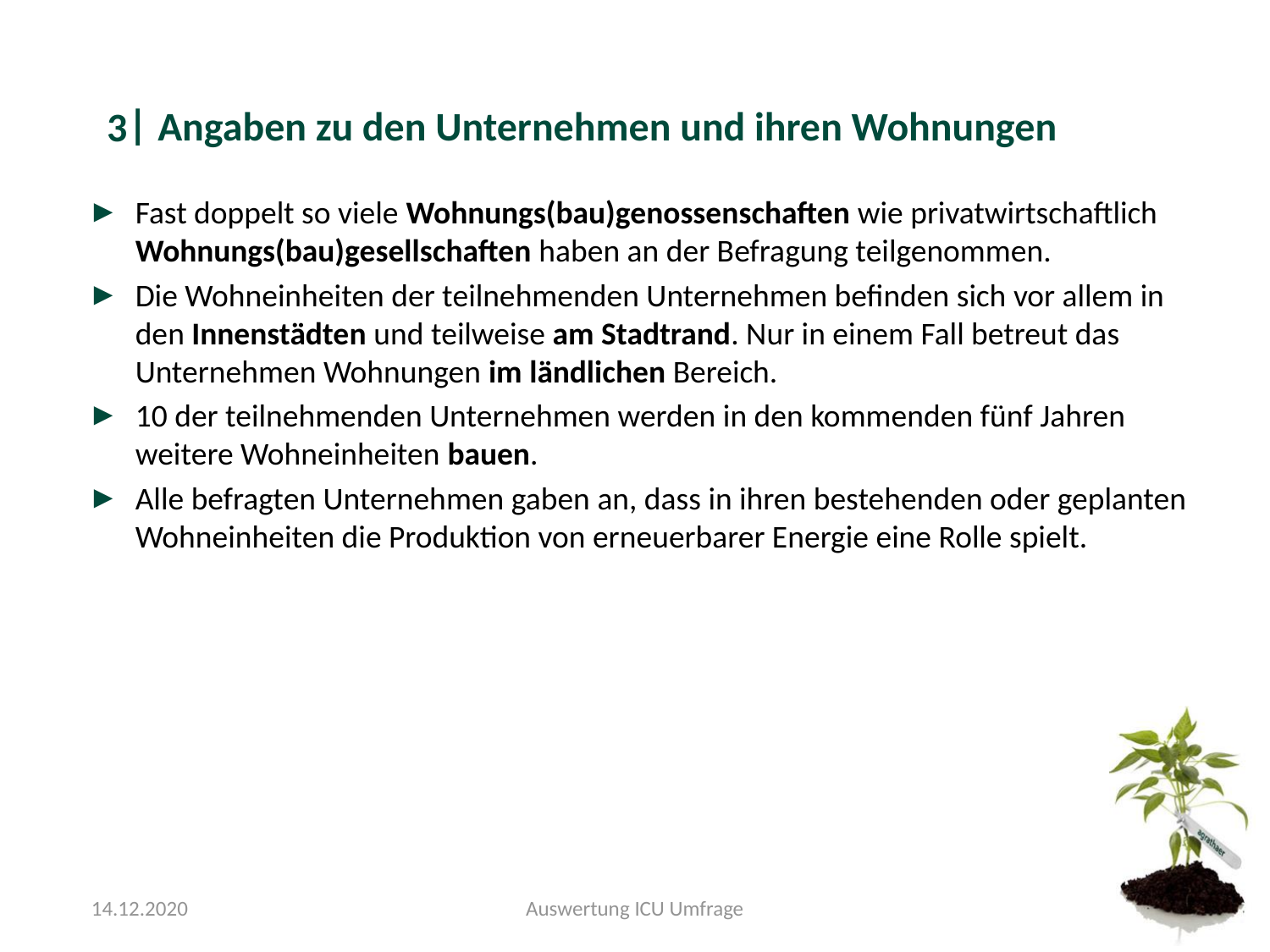

#### Angaben zu den Unternehmen und ihren Wohnungen
3
Fast doppelt so viele Wohnungs(bau)genossenschaften wie privatwirtschaftlich Wohnungs(bau)gesellschaften haben an der Befragung teilgenommen.
Die Wohneinheiten der teilnehmenden Unternehmen befinden sich vor allem in den Innenstädten und teilweise am Stadtrand. Nur in einem Fall betreut das Unternehmen Wohnungen im ländlichen Bereich.
10 der teilnehmenden Unternehmen werden in den kommenden fünf Jahren weitere Wohneinheiten bauen.
Alle befragten Unternehmen gaben an, dass in ihren bestehenden oder geplanten Wohneinheiten die Produktion von erneuerbarer Energie eine Rolle spielt.
14.12.2020
Auswertung ICU Umfrage

### Slide 7
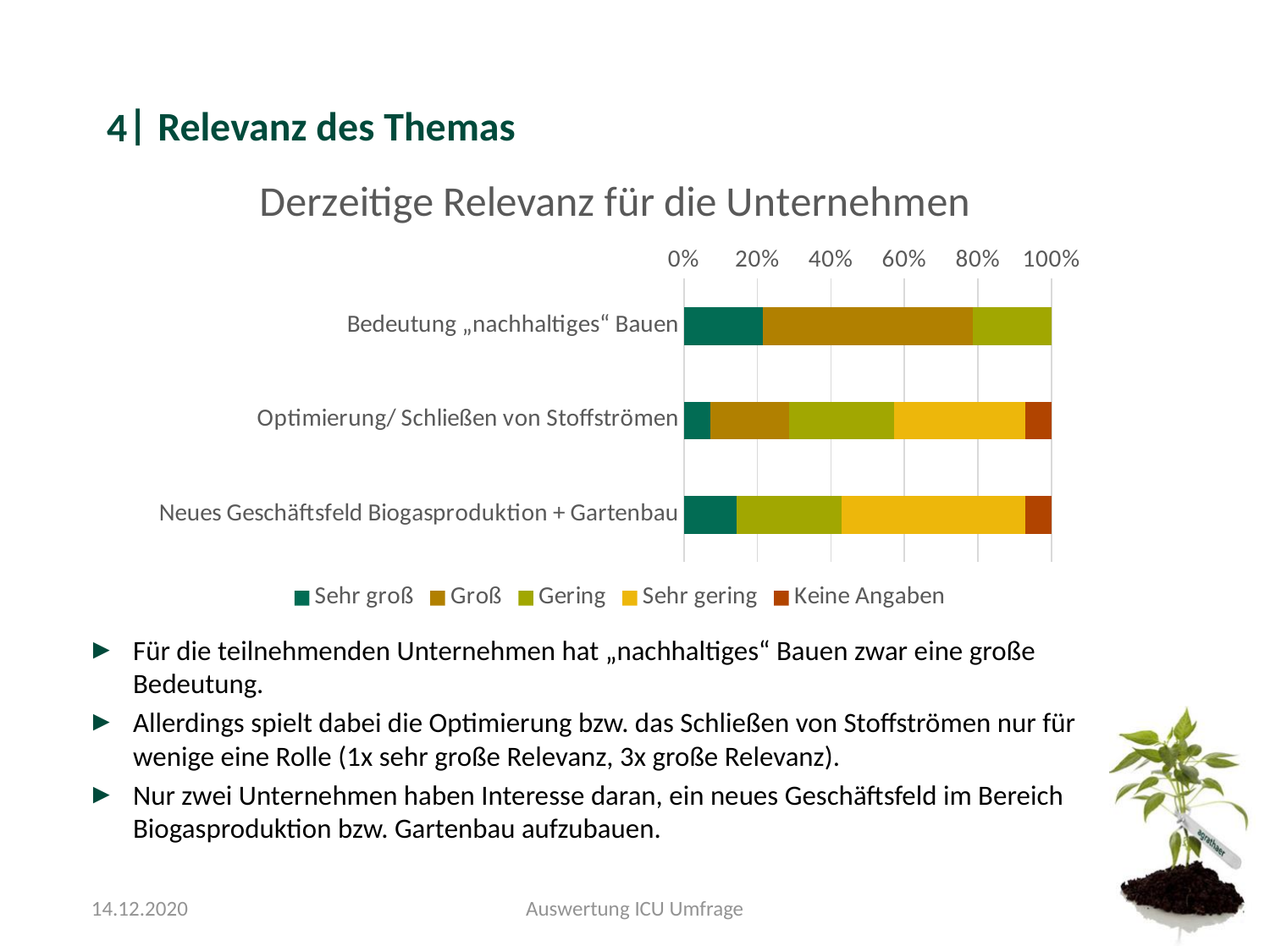

#### Relevanz des Themas
4
###### Chart: Derzeitige Relevanz für die Unternehmen
| Category | Sehr groß | Groß | Gering | Sehr gering | Keine Angaben |
|---|---|---|---|---|---|
| Bedeutung „nachhaltiges“ Bauen | 3.0 | 8.0 | 3.0 | 0.0 | 0.0 |
| Optimierung/ Schließen von Stoffströmen | 1.0 | 3.0 | 4.0 | 5.0 | 1.0 |
| Neues Geschäftsfeld Biogasproduktion + Gartenbau | 2.0 | 0.0 | 4.0 | 7.0 | 1.0 |Für die teilnehmenden Unternehmen hat „nachhaltiges“ Bauen zwar eine große Bedeutung.
Allerdings spielt dabei die Optimierung bzw. das Schließen von Stoffströmen nur für wenige eine Rolle (1x sehr große Relevanz, 3x große Relevanz).
Nur zwei Unternehmen haben Interesse daran, ein neues Geschäftsfeld im Bereich Biogasproduktion bzw. Gartenbau aufzubauen.
14.12.2020
Auswertung ICU Umfrage

### Slide 8
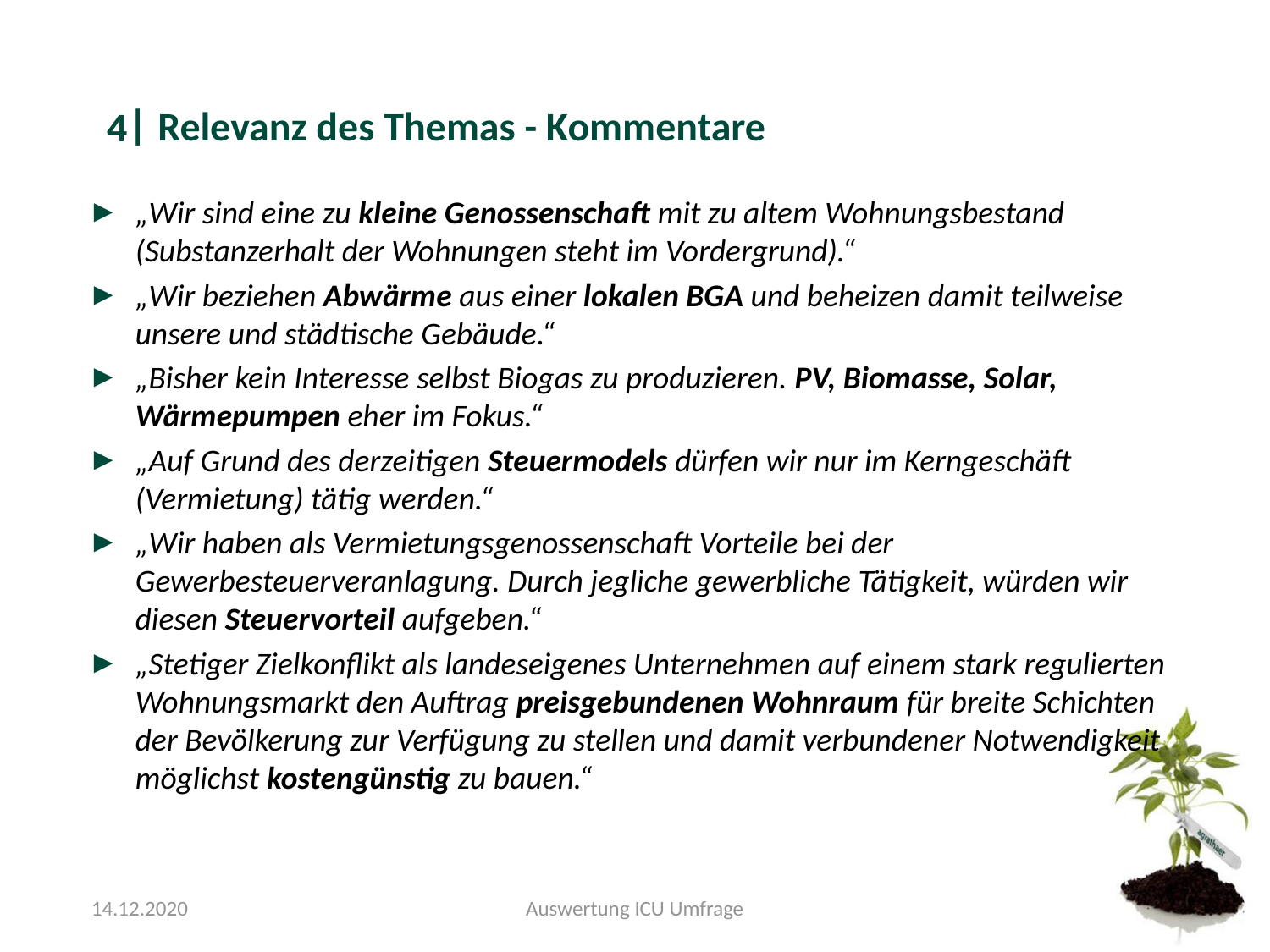

#### Relevanz des Themas - Kommentare
4
„Wir sind eine zu kleine Genossenschaft mit zu altem Wohnungsbestand (Substanzerhalt der Wohnungen steht im Vordergrund).“
„Wir beziehen Abwärme aus einer lokalen BGA und beheizen damit teilweise unsere und städtische Gebäude.“
„Bisher kein Interesse selbst Biogas zu produzieren. PV, Biomasse, Solar, Wärmepumpen eher im Fokus.“
„Auf Grund des derzeitigen Steuermodels dürfen wir nur im Kerngeschäft (Vermietung) tätig werden.“
„Wir haben als Vermietungsgenossenschaft Vorteile bei der Gewerbesteuerveranlagung. Durch jegliche gewerbliche Tätigkeit, würden wir diesen Steuervorteil aufgeben.“
„Stetiger Zielkonflikt als landeseigenes Unternehmen auf einem stark regulierten Wohnungsmarkt den Auftrag preisgebundenen Wohnraum für breite Schichten der Bevölkerung zur Verfügung zu stellen und damit verbundener Notwendigkeit möglichst kostengünstig zu bauen.“
14.12.2020
Auswertung ICU Umfrage

### Slide 9
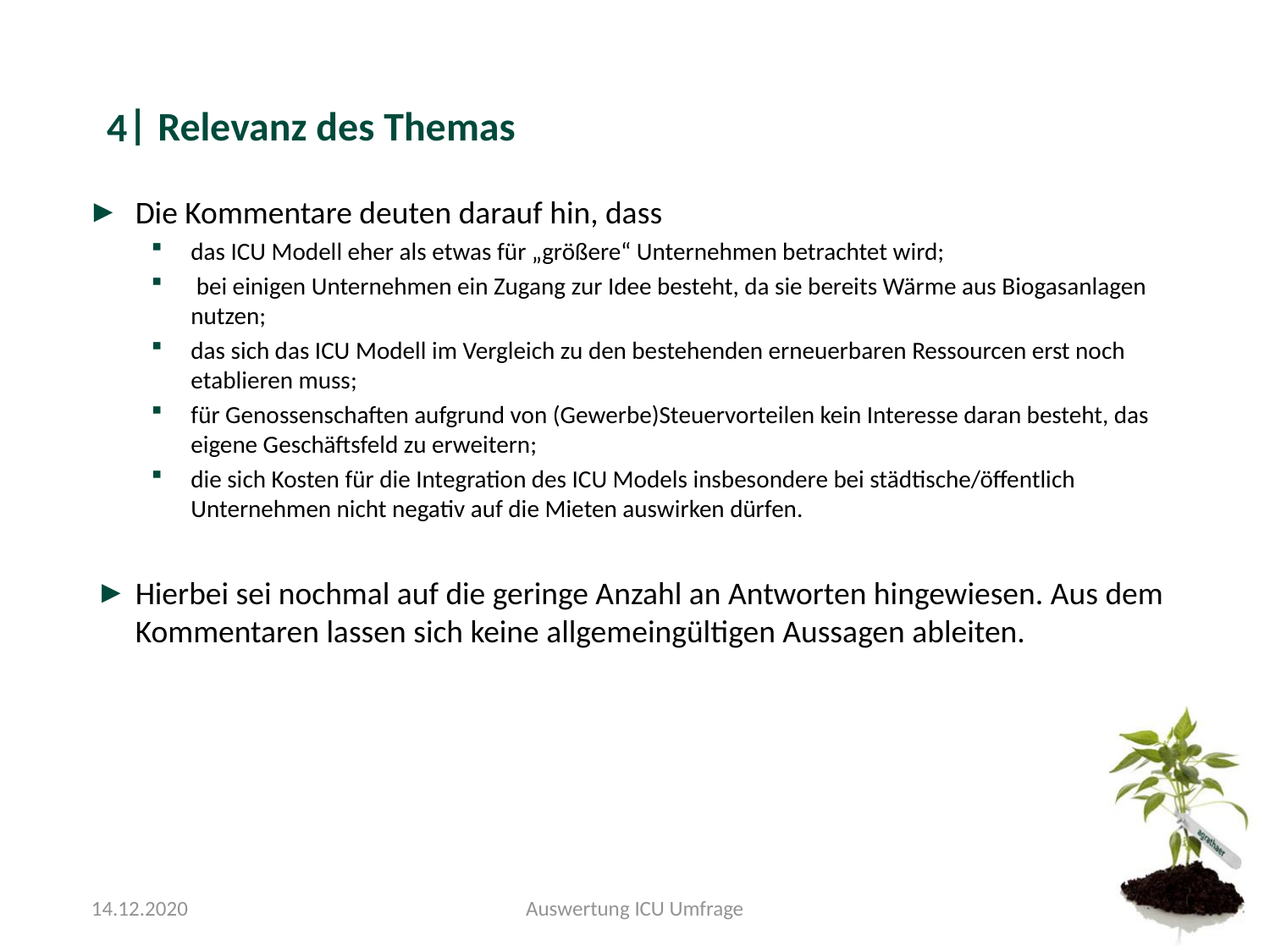

#### Relevanz des Themas
4
Die Kommentare deuten darauf hin, dass
das ICU Modell eher als etwas für „größere“ Unternehmen betrachtet wird;
 bei einigen Unternehmen ein Zugang zur Idee besteht, da sie bereits Wärme aus Biogasanlagen nutzen;
das sich das ICU Modell im Vergleich zu den bestehenden erneuerbaren Ressourcen erst noch etablieren muss;
für Genossenschaften aufgrund von (Gewerbe)Steuervorteilen kein Interesse daran besteht, das eigene Geschäftsfeld zu erweitern;
die sich Kosten für die Integration des ICU Models insbesondere bei städtische/öffentlich Unternehmen nicht negativ auf die Mieten auswirken dürfen.
Hierbei sei nochmal auf die geringe Anzahl an Antworten hingewiesen. Aus dem Kommentaren lassen sich keine allgemeingültigen Aussagen ableiten.
14.12.2020
Auswertung ICU Umfrage

### Slide 10
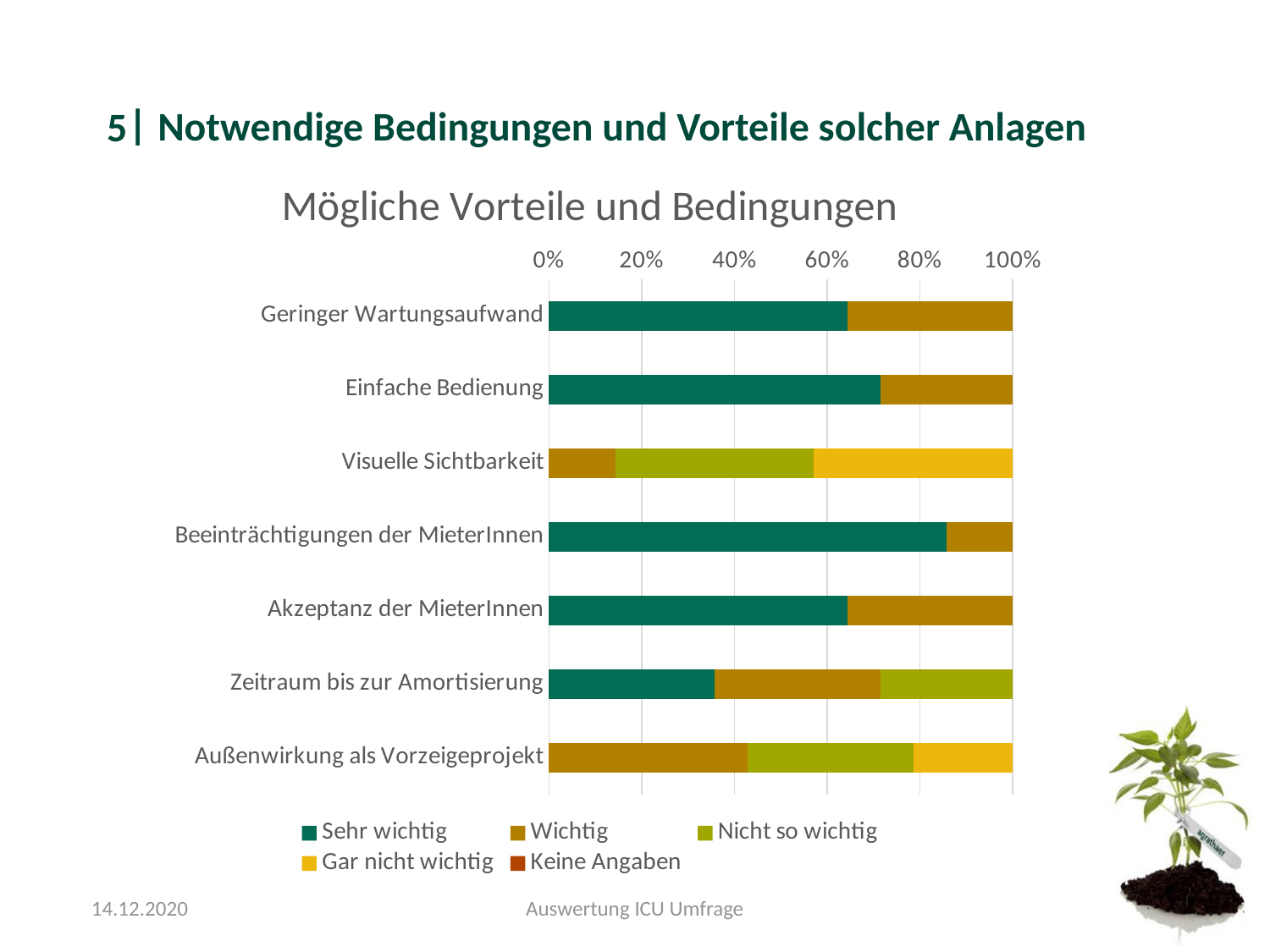

#### Notwendige Bedingungen und Vorteile solcher Anlagen
5
###### Chart: Mögliche Vorteile und Bedingungen
| Category | Sehr wichtig | Wichtig | Nicht so wichtig | Gar nicht wichtig | Keine Angaben |
|---|---|---|---|---|---|
| Geringer Wartungsaufwand | 9.0 | 5.0 | 0.0 | 0.0 | 0.0 |
| Einfache Bedienung | 10.0 | 4.0 | 0.0 | 0.0 | 0.0 |
| Visuelle Sichtbarkeit | 0.0 | 2.0 | 6.0 | 6.0 | 0.0 |
| Beeinträchtigungen der MieterInnen | 12.0 | 2.0 | 0.0 | 0.0 | 0.0 |
| Akzeptanz der MieterInnen | 9.0 | 5.0 | 0.0 | 0.0 | 0.0 |
| Zeitraum bis zur Amortisierung | 5.0 | 5.0 | 4.0 | 0.0 | 0.0 |
| Außenwirkung als Vorzeigeprojekt | 0.0 | 6.0 | 5.0 | 3.0 | 0.0 |14.12.2020
Auswertung ICU Umfrage

### Slide 11
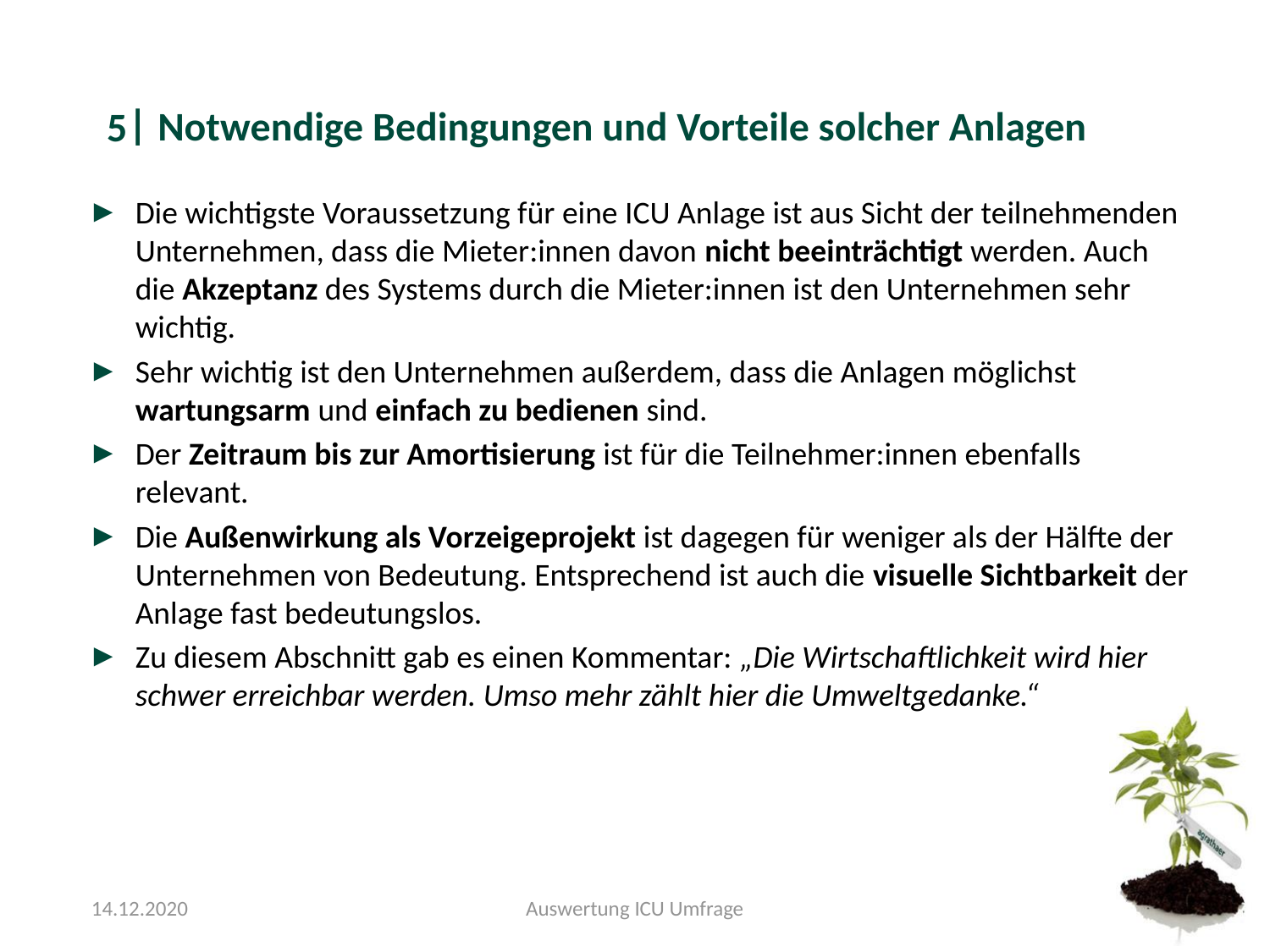

#### Notwendige Bedingungen und Vorteile solcher Anlagen
5
Die wichtigste Voraussetzung für eine ICU Anlage ist aus Sicht der teilnehmenden Unternehmen, dass die Mieter:innen davon nicht beeinträchtigt werden. Auch die Akzeptanz des Systems durch die Mieter:innen ist den Unternehmen sehr wichtig.
Sehr wichtig ist den Unternehmen außerdem, dass die Anlagen möglichst wartungsarm und einfach zu bedienen sind.
Der Zeitraum bis zur Amortisierung ist für die Teilnehmer:innen ebenfalls relevant.
Die Außenwirkung als Vorzeigeprojekt ist dagegen für weniger als der Hälfte der Unternehmen von Bedeutung. Entsprechend ist auch die visuelle Sichtbarkeit der Anlage fast bedeutungslos.
Zu diesem Abschnitt gab es einen Kommentar: „Die Wirtschaftlichkeit wird hier schwer erreichbar werden. Umso mehr zählt hier die Umweltgedanke.“
14.12.2020
Auswertung ICU Umfrage

### Slide 12
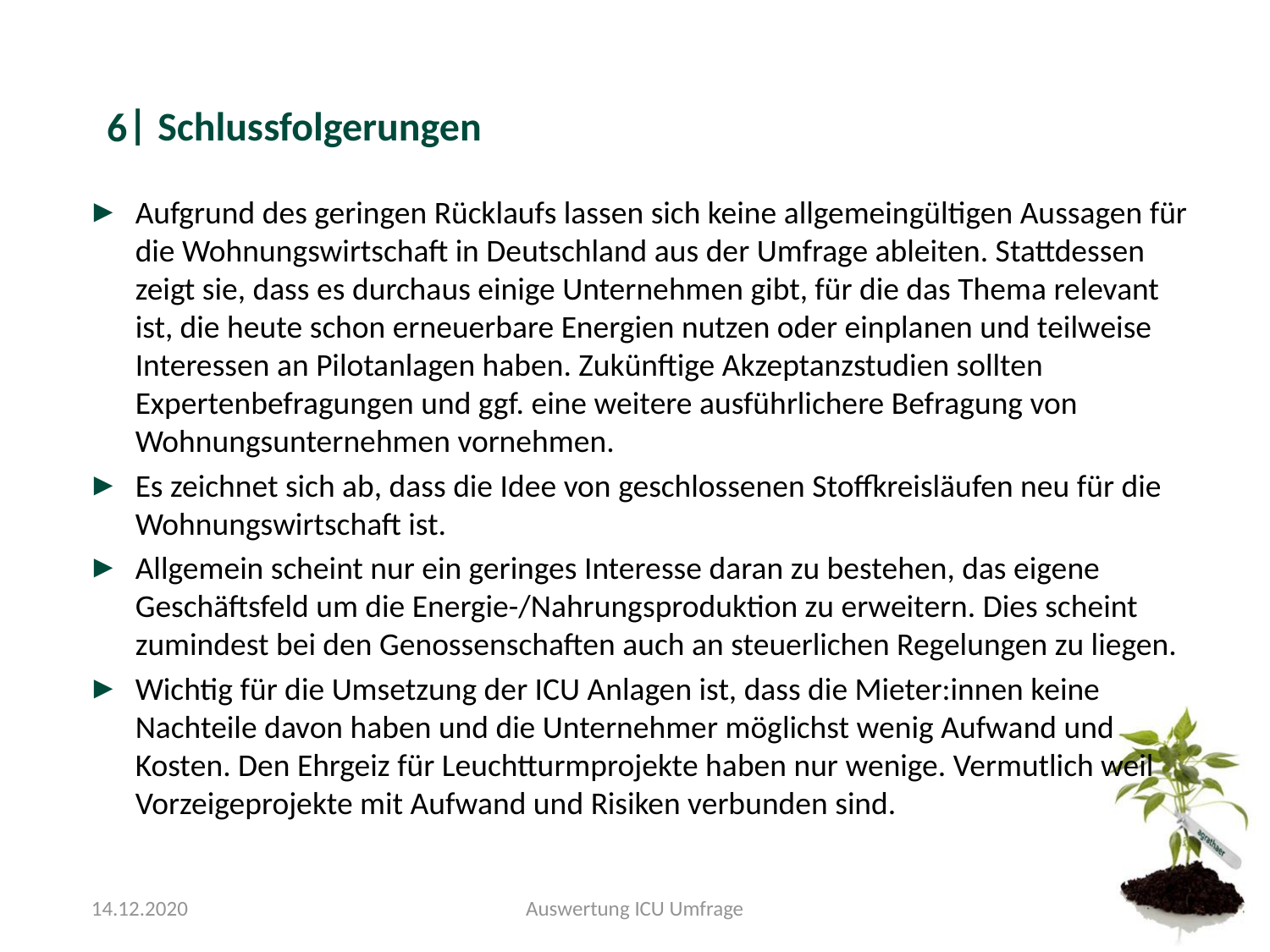

#### Schlussfolgerungen
6
Aufgrund des geringen Rücklaufs lassen sich keine allgemeingültigen Aussagen für die Wohnungswirtschaft in Deutschland aus der Umfrage ableiten. Stattdessen zeigt sie, dass es durchaus einige Unternehmen gibt, für die das Thema relevant ist, die heute schon erneuerbare Energien nutzen oder einplanen und teilweise Interessen an Pilotanlagen haben. Zukünftige Akzeptanzstudien sollten Expertenbefragungen und ggf. eine weitere ausführlichere Befragung von Wohnungsunternehmen vornehmen.
Es zeichnet sich ab, dass die Idee von geschlossenen Stoffkreisläufen neu für die Wohnungswirtschaft ist.
Allgemein scheint nur ein geringes Interesse daran zu bestehen, das eigene Geschäftsfeld um die Energie-/Nahrungsproduktion zu erweitern. Dies scheint zumindest bei den Genossenschaften auch an steuerlichen Regelungen zu liegen.
Wichtig für die Umsetzung der ICU Anlagen ist, dass die Mieter:innen keine Nachteile davon haben und die Unternehmer möglichst wenig Aufwand und Kosten. Den Ehrgeiz für Leuchtturmprojekte haben nur wenige. Vermutlich weil Vorzeigeprojekte mit Aufwand und Risiken verbunden sind.
14.12.2020
Auswertung ICU Umfrage

### Slide 13
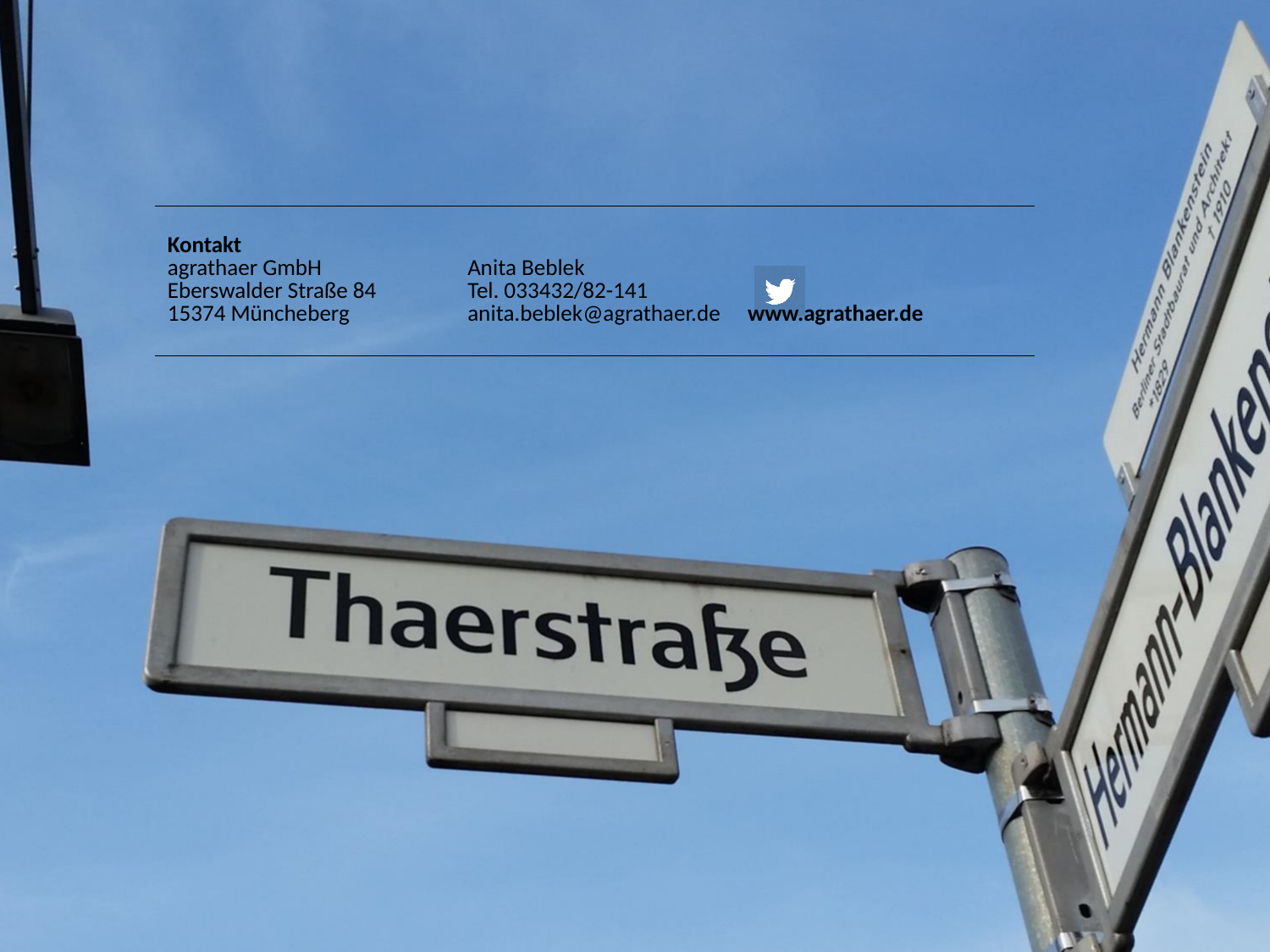

| Kontakt agrathaer GmbH Eberswalder Straße 84 15374 Müncheberg | Anita Beblek Tel. 033432/82-141 | www.agrathaer.de |
| --- | --- | --- |
