## Supplementary file 5 for "Integrated cycles for urban biomass as a strategy to promote a CO_2_-neutral society – a feasibility study"

### Slide 1
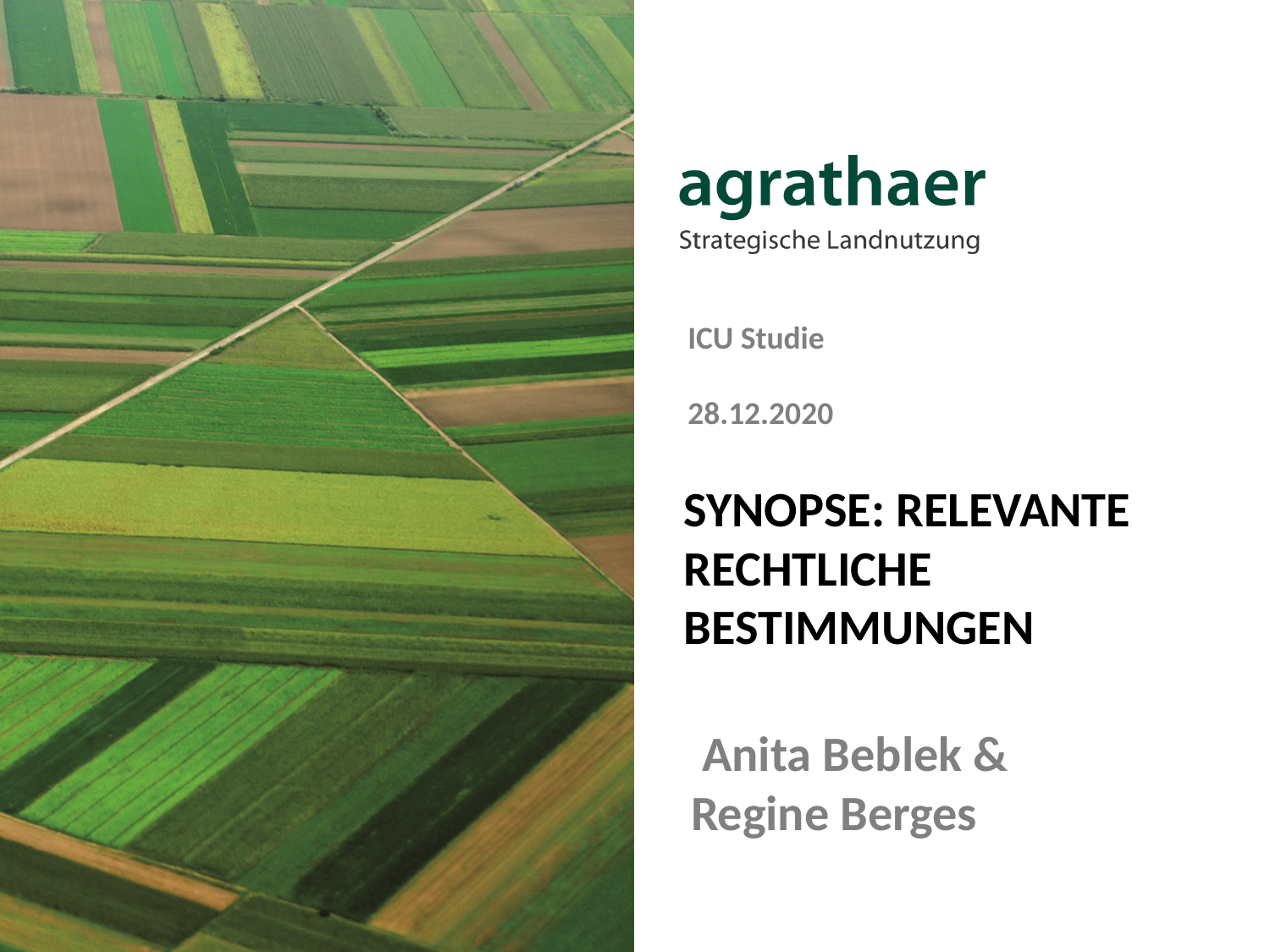

ICU Studie
28.12.2020
#### Synopse: relevante rechtliche Bestimmungen
 Anita Beblek & Regine Berges

### Slide 2
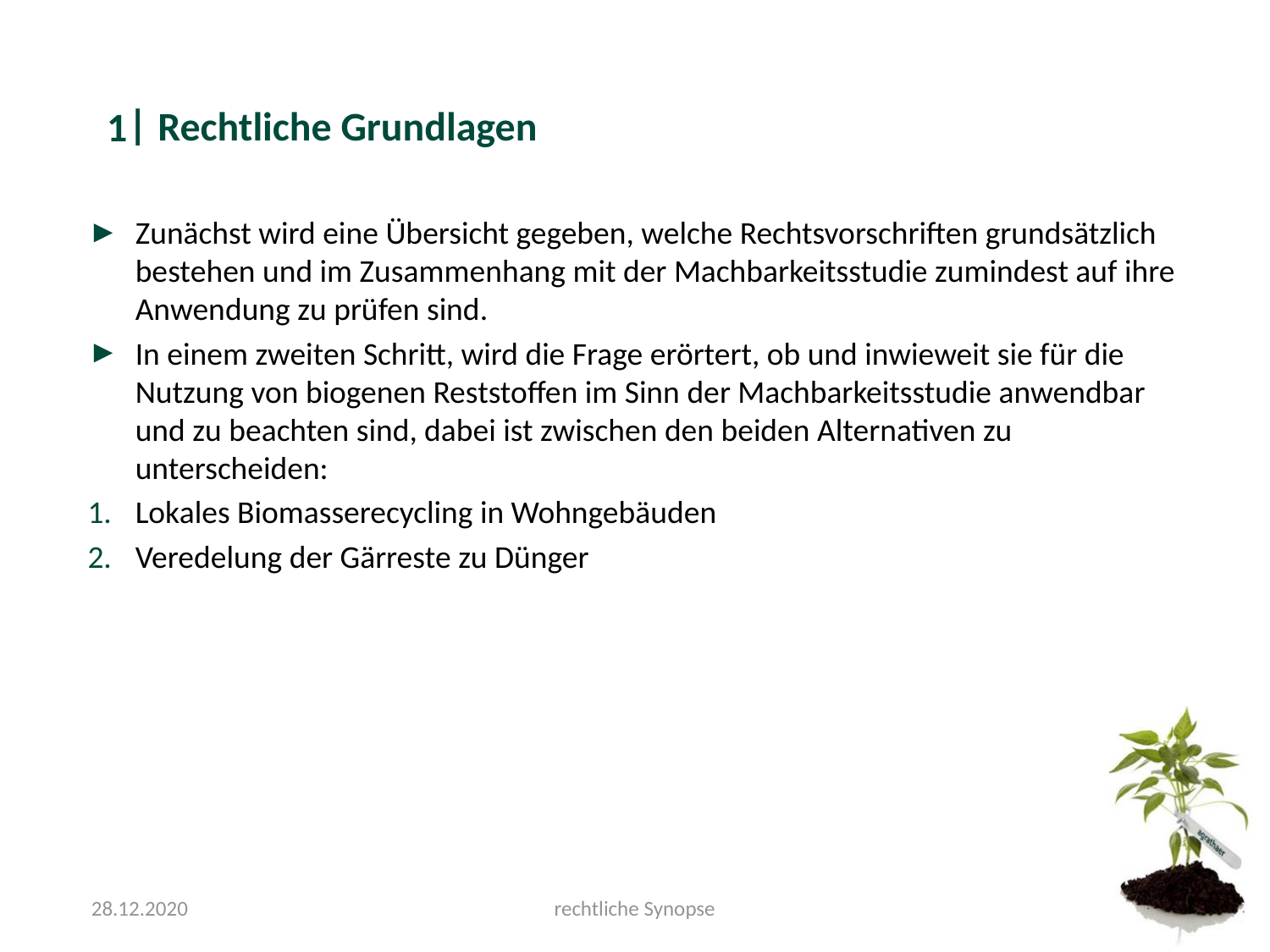

#### Rechtliche Grundlagen
1
Zunächst wird eine Übersicht gegeben, welche Rechtsvorschriften grundsätzlich bestehen und im Zusammenhang mit der Machbarkeitsstudie zumindest auf ihre Anwendung zu prüfen sind.
In einem zweiten Schritt, wird die Frage erörtert, ob und inwieweit sie für die Nutzung von biogenen Reststoffen im Sinn der Machbarkeitsstudie anwendbar und zu beachten sind, dabei ist zwischen den beiden Alternativen zu unterscheiden:
Lokales Biomasserecycling in Wohngebäuden
Veredelung der Gärreste zu Dünger
28.12.2020
rechtliche Synopse

### Slide 3
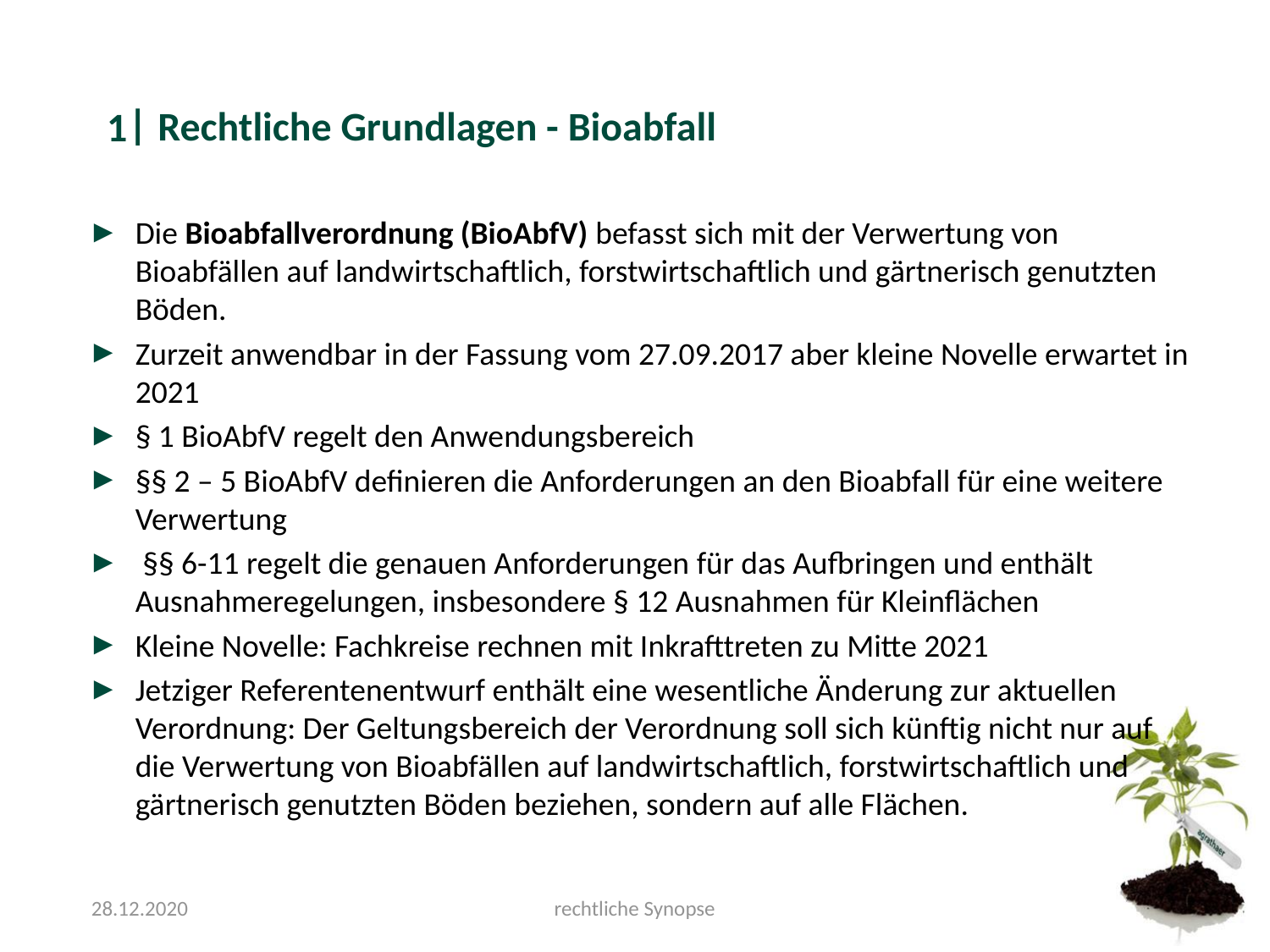

#### Rechtliche Grundlagen - Bioabfall
1
Die Bioabfallverordnung (BioAbfV) befasst sich mit der Verwertung von Bioabfällen auf landwirtschaftlich, forstwirtschaftlich und gärtnerisch genutzten Böden.
Zurzeit anwendbar in der Fassung vom 27.09.2017 aber kleine Novelle erwartet in 2021
§ 1 BioAbfV regelt den Anwendungsbereich
§§ 2 – 5 BioAbfV definieren die Anforderungen an den Bioabfall für eine weitere Verwertung
 §§ 6-11 regelt die genauen Anforderungen für das Aufbringen und enthält Ausnahmeregelungen, insbesondere § 12 Ausnahmen für Kleinflächen
Kleine Novelle: Fachkreise rechnen mit Inkrafttreten zu Mitte 2021
Jetziger Referentenentwurf enthält eine wesentliche Änderung zur aktuellen Verordnung: Der Geltungsbereich der Verordnung soll sich künftig nicht nur auf die Verwertung von Bioabfällen auf landwirtschaftlich, forstwirtschaftlich und gärtnerisch genutzten Böden beziehen, sondern auf alle Flächen.
28.12.2020
rechtliche Synopse

### Slide 4
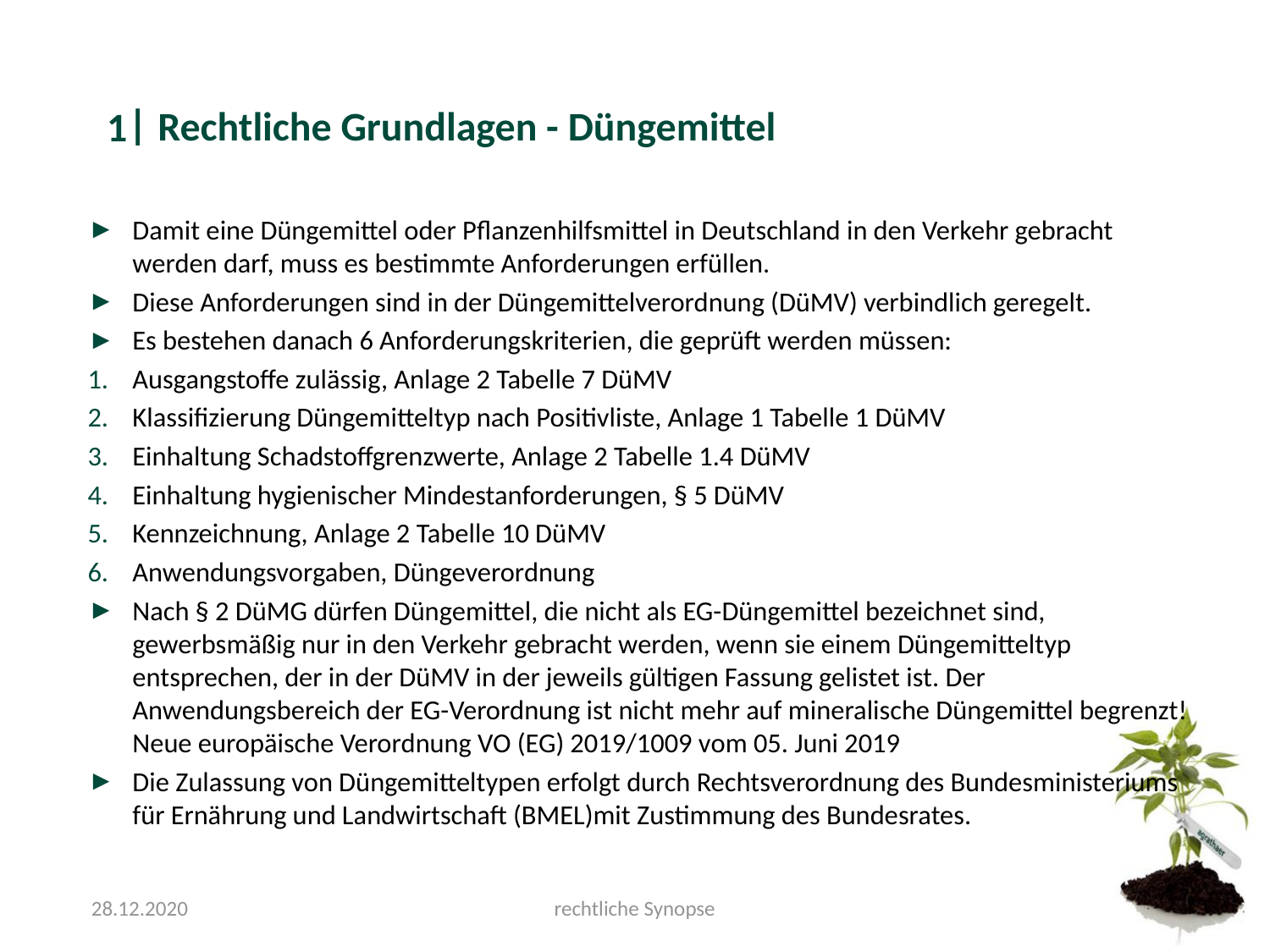

#### Rechtliche Grundlagen - Düngemittel
1
Damit eine Düngemittel oder Pflanzenhilfsmittel in Deutschland in den Verkehr gebracht werden darf, muss es bestimmte Anforderungen erfüllen.
Diese Anforderungen sind in der Düngemittelverordnung (DüMV) verbindlich geregelt.
Es bestehen danach 6 Anforderungskriterien, die geprüft werden müssen:
Ausgangstoffe zulässig, Anlage 2 Tabelle 7 DüMV
Klassifizierung Düngemitteltyp nach Positivliste, Anlage 1 Tabelle 1 DüMV
Einhaltung Schadstoffgrenzwerte, Anlage 2 Tabelle 1.4 DüMV
Einhaltung hygienischer Mindestanforderungen, § 5 DüMV
Kennzeichnung, Anlage 2 Tabelle 10 DüMV
Anwendungsvorgaben, Düngeverordnung
Nach § 2 DüMG dürfen Düngemittel, die nicht als EG-Düngemittel bezeichnet sind, gewerbsmäßig nur in den Verkehr gebracht werden, wenn sie einem Düngemitteltyp entsprechen, der in der DüMV in der jeweils gültigen Fassung gelistet ist. Der Anwendungsbereich der EG-Verordnung ist nicht mehr auf mineralische Düngemittel begrenzt! Neue europäische Verordnung VO (EG) 2019/1009 vom 05. Juni 2019
Die Zulassung von Düngemitteltypen erfolgt durch Rechtsverordnung des Bundesministeriums für Ernährung und Landwirtschaft (BMEL)mit Zustimmung des Bundesrates.
28.12.2020
rechtliche Synopse

### Slide 5
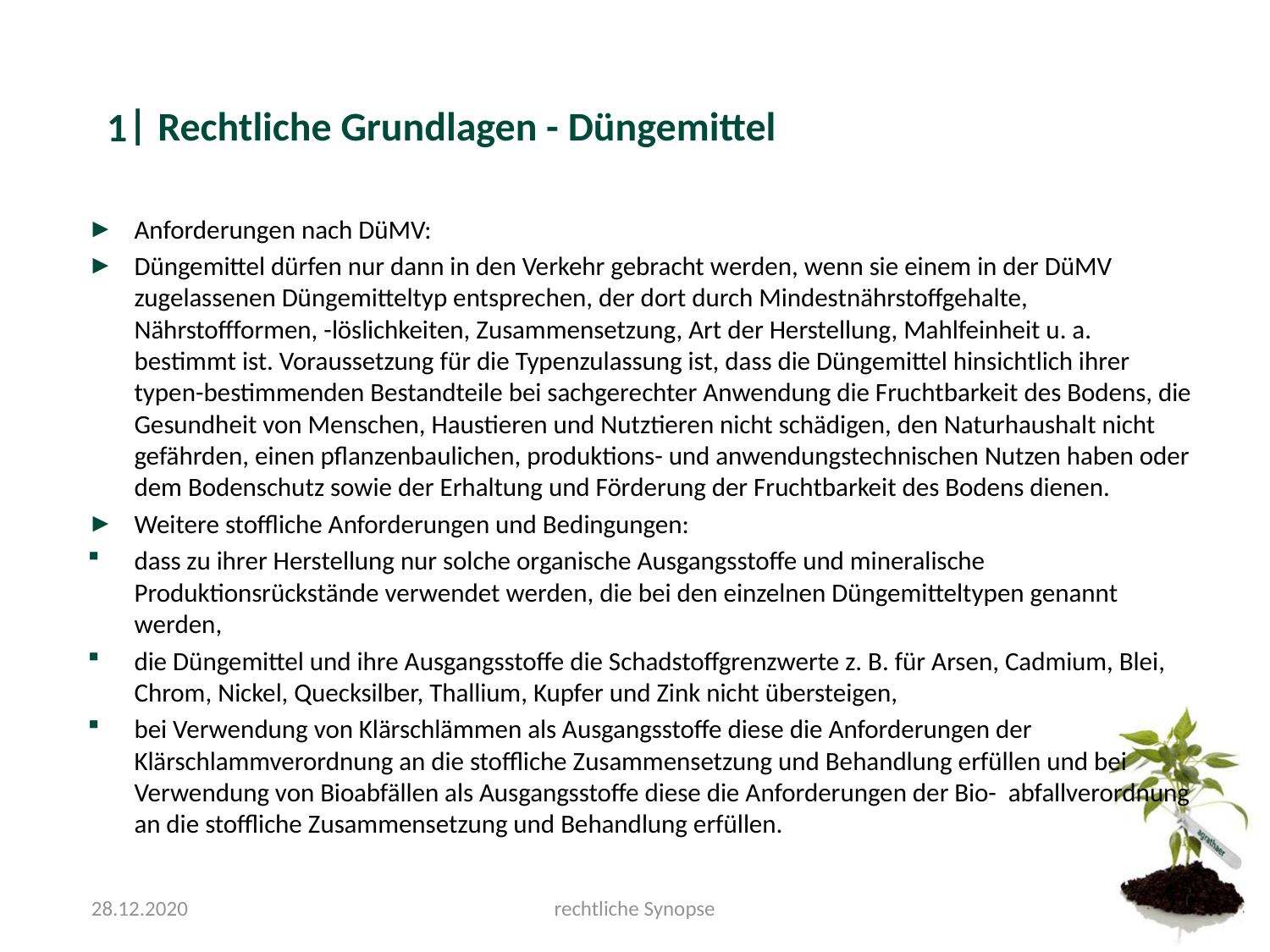

#### Rechtliche Grundlagen - Düngemittel
1
Anforderungen nach DüMV:
Düngemittel dürfen nur dann in den Verkehr gebracht werden, wenn sie einem in der DüMV zugelassenen Düngemitteltyp entsprechen, der dort durch Mindestnährstoffgehalte, Nährstoffformen, -löslichkeiten, Zusammensetzung, Art der Herstellung, Mahlfeinheit u. a. bestimmt ist. Voraussetzung für die Typenzulassung ist, dass die Düngemittel hinsichtlich ihrer typen-bestimmenden Bestandteile bei sachgerechter Anwendung die Fruchtbarkeit des Bodens, die Gesundheit von Menschen, Haustieren und Nutztieren nicht schädigen, den Naturhaushalt nicht gefährden, einen pflanzenbaulichen, produktions- und anwendungstechnischen Nutzen haben oder dem Bodenschutz sowie der Erhaltung und Förderung der Fruchtbarkeit des Bodens dienen.
Weitere stoffliche Anforderungen und Bedingungen:
dass zu ihrer Herstellung nur solche organische Ausgangsstoffe und mineralische Produktionsrückstände verwendet werden, die bei den einzelnen Düngemitteltypen genannt werden,
die Düngemittel und ihre Ausgangsstoffe die Schadstoffgrenzwerte z. B. für Arsen, Cadmium, Blei, Chrom, Nickel, Quecksilber, Thallium, Kupfer und Zink nicht übersteigen,
bei Verwendung von Klärschlämmen als Ausgangsstoffe diese die Anforderungen der Klärschlammverordnung an die stoffliche Zusammensetzung und Behandlung erfüllen und bei Verwendung von Bioabfällen als Ausgangsstoffe diese die Anforderungen der Bio- abfallverordnung an die stoffliche Zusammensetzung und Behandlung erfüllen.
28.12.2020
rechtliche Synopse

### Slide 6
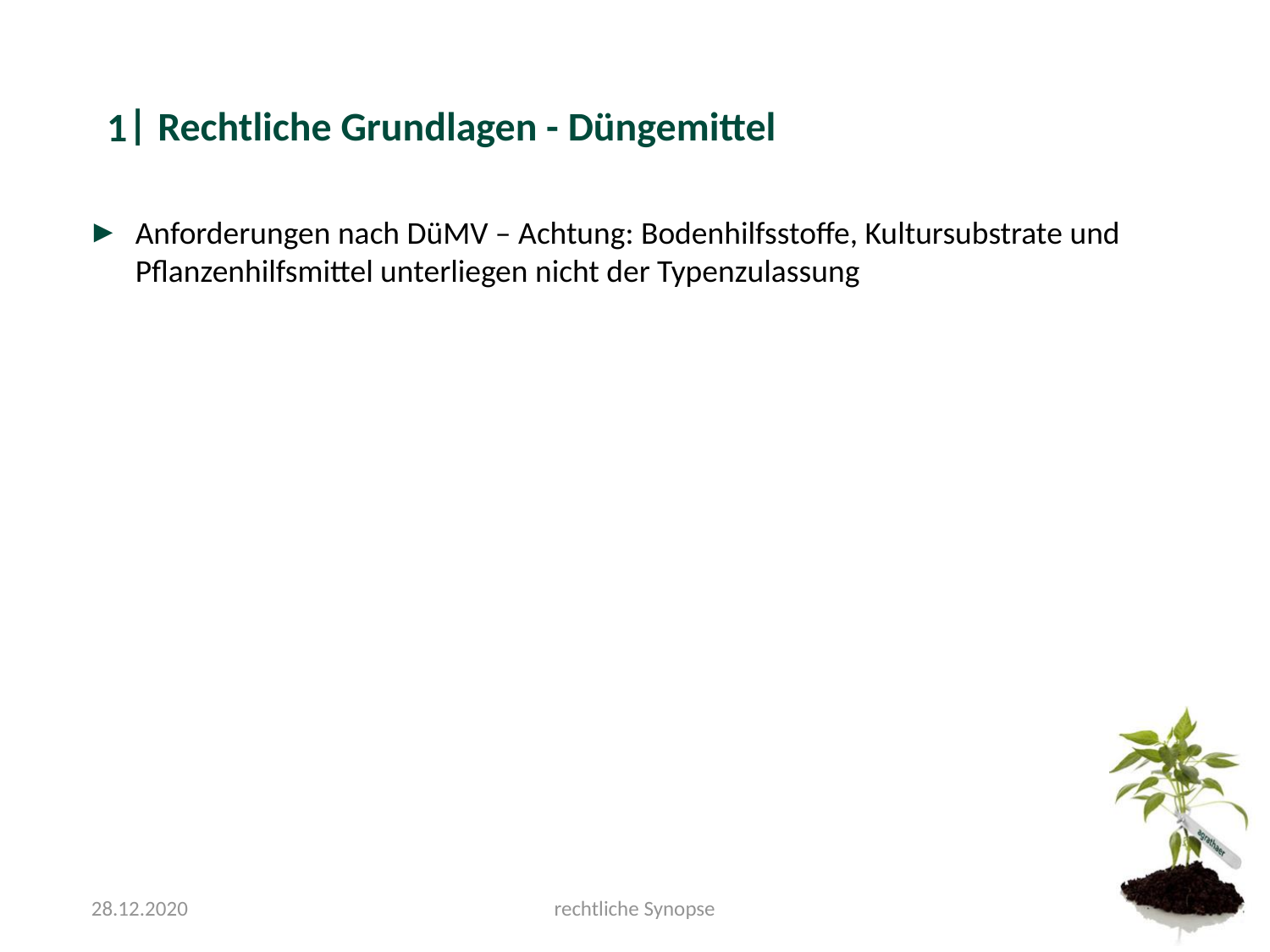

#### Rechtliche Grundlagen - Düngemittel
1
Anforderungen nach DüMV – Achtung: Bodenhilfsstoffe, Kultursubstrate und Pflanzenhilfsmittel unterliegen nicht der Typenzulassung
28.12.2020
rechtliche Synopse

### Slide 7
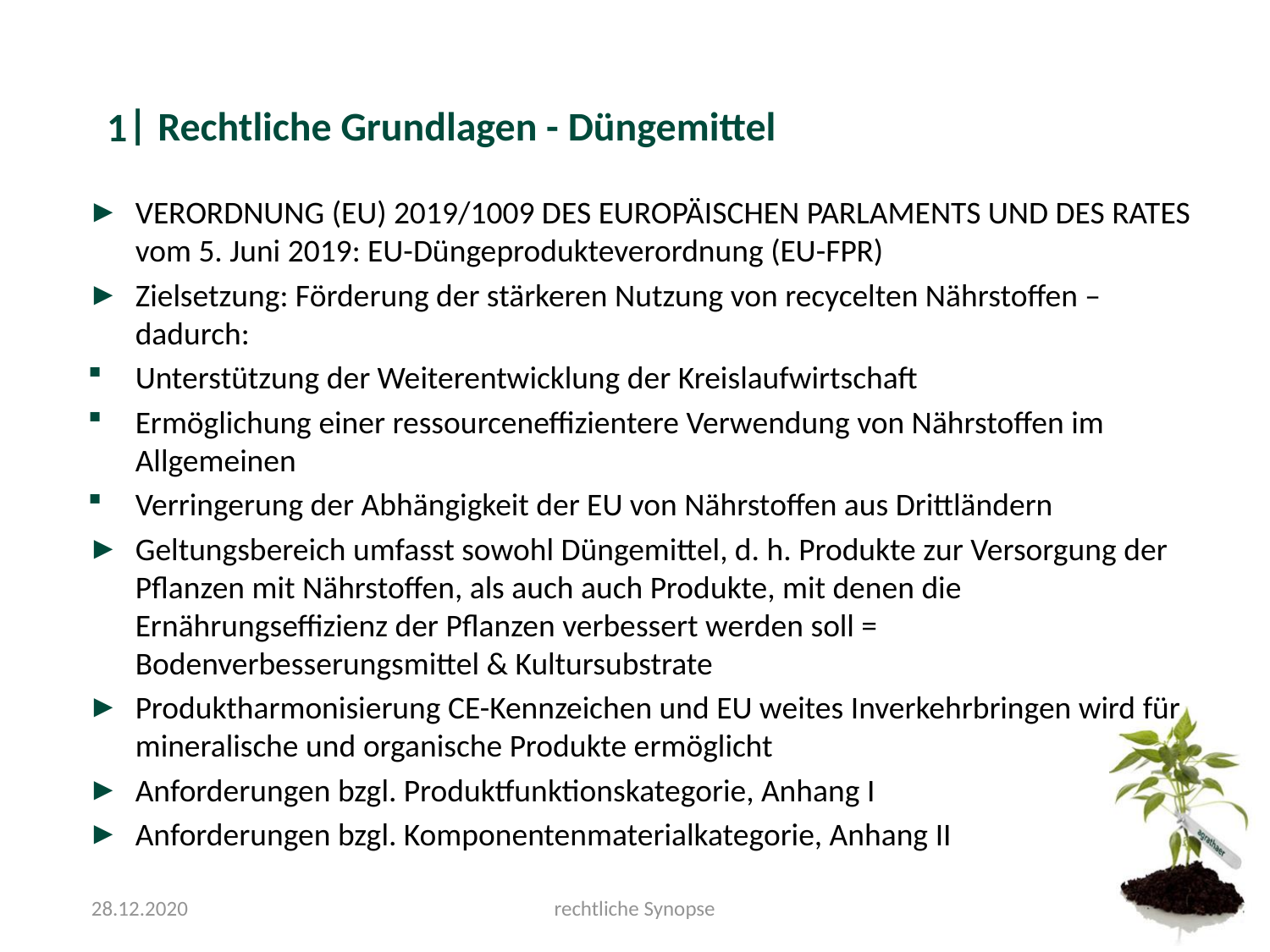

#### Rechtliche Grundlagen - Düngemittel
1
VERORDNUNG (EU) 2019/1009 DES EUROPÄISCHEN PARLAMENTS UND DES RATES vom 5. Juni 2019: EU-Düngeprodukteverordnung (EU-FPR)
Zielsetzung: Förderung der stärkeren Nutzung von recycelten Nährstoffen – dadurch:
Unterstützung der Weiterentwicklung der Kreislaufwirtschaft
Ermöglichung einer ressourceneffizientere Verwendung von Nährstoffen im Allgemeinen
Verringerung der Abhängigkeit der EU von Nährstoffen aus Drittländern
Geltungsbereich umfasst sowohl Düngemittel, d. h. Produkte zur Versorgung der Pflanzen mit Nährstoffen, als auch auch Produkte, mit denen die Ernährungseffizienz der Pflanzen verbessert werden soll = Bodenverbesserungsmittel & Kultursubstrate
Produktharmonisierung CE-Kennzeichen und EU weites Inverkehrbringen wird für mineralische und organische Produkte ermöglicht
Anforderungen bzgl. Produktfunktionskategorie, Anhang I
Anforderungen bzgl. Komponentenmaterialkategorie, Anhang II
28.12.2020
rechtliche Synopse

### Slide 8
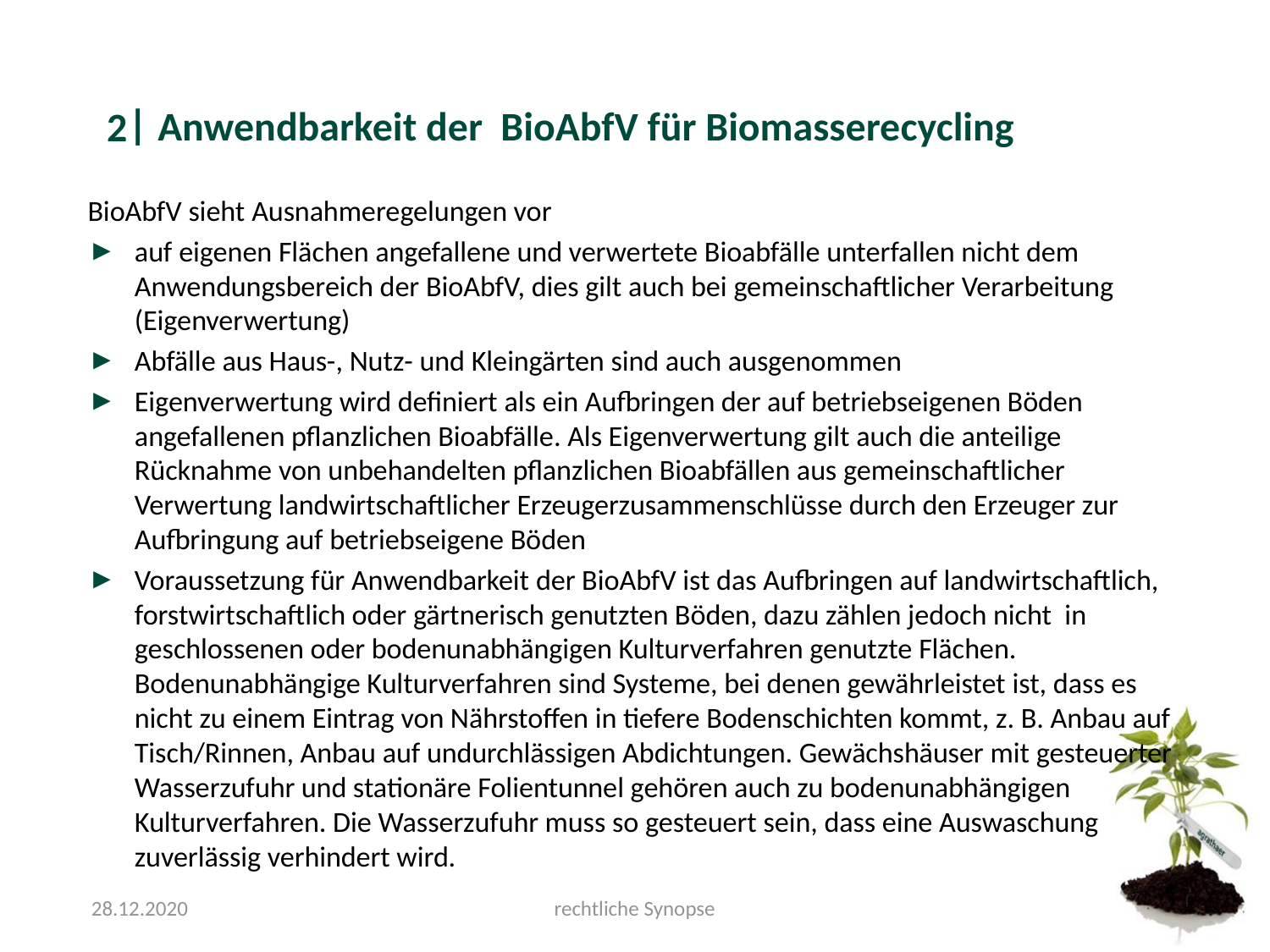

#### Anwendbarkeit der BioAbfV für Biomasserecycling
2
BioAbfV sieht Ausnahmeregelungen vor
auf eigenen Flächen angefallene und verwertete Bioabfälle unterfallen nicht dem Anwendungsbereich der BioAbfV, dies gilt auch bei gemeinschaftlicher Verarbeitung (Eigenverwertung)
Abfälle aus Haus-, Nutz- und Kleingärten sind auch ausgenommen
Eigenverwertung wird definiert als ein Aufbringen der auf betriebseigenen Böden angefallenen pflanzlichen Bioabfälle. Als Eigenverwertung gilt auch die anteilige Rücknahme von unbehandelten pflanzlichen Bioabfällen aus gemeinschaftlicher Verwertung landwirtschaftlicher Erzeugerzusammenschlüsse durch den Erzeuger zur Aufbringung auf betriebseigene Böden
Voraussetzung für Anwendbarkeit der BioAbfV ist das Aufbringen auf landwirtschaftlich, forstwirtschaftlich oder gärtnerisch genutzten Böden, dazu zählen jedoch nicht in geschlossenen oder bodenunabhängigen Kulturverfahren genutzte Flächen. Bodenunabhängige Kulturverfahren sind Systeme, bei denen gewährleistet ist, dass es nicht zu einem Eintrag von Nährstoffen in tiefere Bodenschichten kommt, z. B. Anbau auf Tisch/Rinnen, Anbau auf undurchlässigen Abdichtungen. Gewächshäuser mit gesteuerter Wasserzufuhr und stationäre Folientunnel gehören auch zu bodenunabhängigen Kulturverfahren. Die Wasserzufuhr muss so gesteuert sein, dass eine Auswaschung zuverlässig verhindert wird.
28.12.2020
rechtliche Synopse

### Slide 9
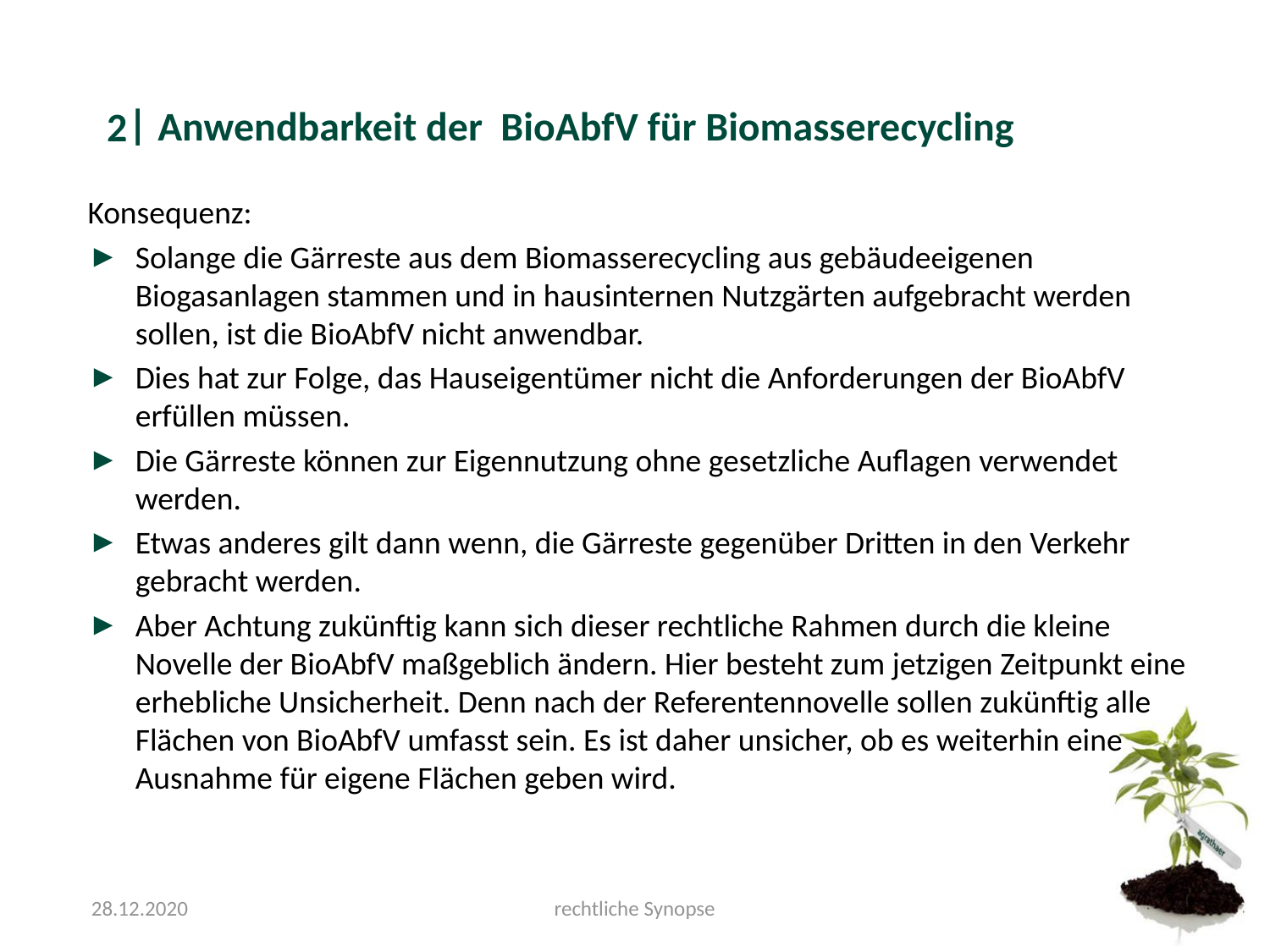

#### Anwendbarkeit der BioAbfV für Biomasserecycling
2
Konsequenz:
Solange die Gärreste aus dem Biomasserecycling aus gebäudeeigenen Biogasanlagen stammen und in hausinternen Nutzgärten aufgebracht werden sollen, ist die BioAbfV nicht anwendbar.
Dies hat zur Folge, das Hauseigentümer nicht die Anforderungen der BioAbfV erfüllen müssen.
Die Gärreste können zur Eigennutzung ohne gesetzliche Auflagen verwendet werden.
Etwas anderes gilt dann wenn, die Gärreste gegenüber Dritten in den Verkehr gebracht werden.
Aber Achtung zukünftig kann sich dieser rechtliche Rahmen durch die kleine Novelle der BioAbfV maßgeblich ändern. Hier besteht zum jetzigen Zeitpunkt eine erhebliche Unsicherheit. Denn nach der Referentennovelle sollen zukünftig alle Flächen von BioAbfV umfasst sein. Es ist daher unsicher, ob es weiterhin eine Ausnahme für eigene Flächen geben wird.
28.12.2020
rechtliche Synopse

### Slide 10
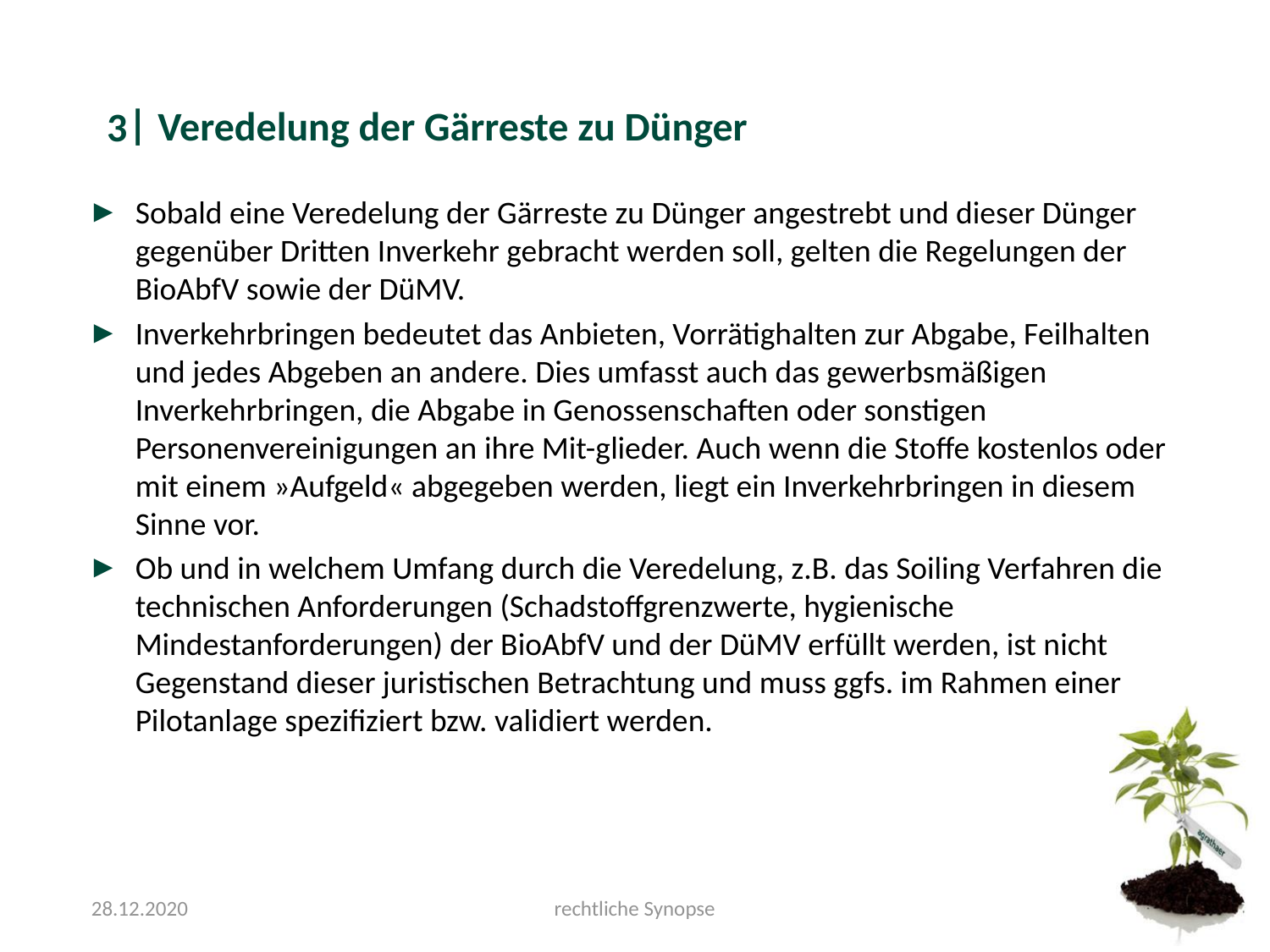

#### Veredelung der Gärreste zu Dünger
3
Sobald eine Veredelung der Gärreste zu Dünger angestrebt und dieser Dünger gegenüber Dritten Inverkehr gebracht werden soll, gelten die Regelungen der BioAbfV sowie der DüMV.
Inverkehrbringen bedeutet das Anbieten, Vorrätighalten zur Abgabe, Feilhalten und jedes Abgeben an andere. Dies umfasst auch das gewerbsmäßigen Inverkehrbringen, die Abgabe in Genossenschaften oder sonstigen Personenvereinigungen an ihre Mit-glieder. Auch wenn die Stoffe kostenlos oder mit einem »Aufgeld« abgegeben werden, liegt ein Inverkehrbringen in diesem Sinne vor.
Ob und in welchem Umfang durch die Veredelung, z.B. das Soiling Verfahren die technischen Anforderungen (Schadstoffgrenzwerte, hygienische Mindestanforderungen) der BioAbfV und der DüMV erfüllt werden, ist nicht Gegenstand dieser juristischen Betrachtung und muss ggfs. im Rahmen einer Pilotanlage spezifiziert bzw. validiert werden.
28.12.2020
rechtliche Synopse

### Slide 11
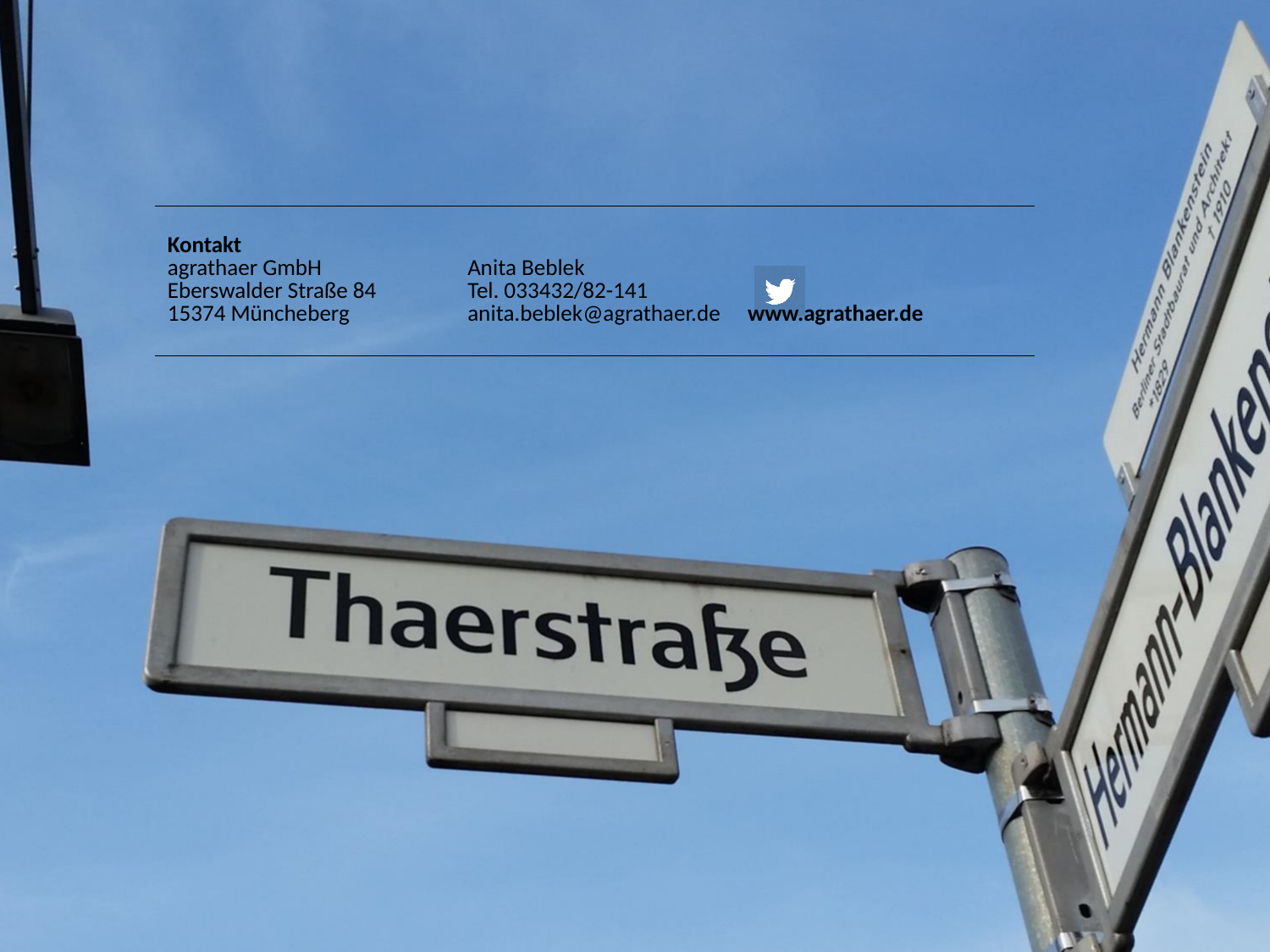

| Kontakt agrathaer GmbH Eberswalder Straße 84 15374 Müncheberg | Anita Beblek Tel. 033432/82-141 | www.agrathaer.de |
| --- | --- | --- |
