## Supplementary file 9 for "Integrated cycles for urban biomass as a strategy to promote a CO_2_-neutral society – a feasibility study"

Parameter for ICU-models in Simba#Biogas


```
1.  <block name="main fermenter" blockType="sisiBlock" x="625" y="230" w="125" h="95"
    drawFrame="False" help="simba#_biogas_reactor_vffg_h.pdf">
2.      <extra blockID="badmLib.adm_vffg_h" HlpIdx="">
3.        <Module id="main fermenter"
    subtype="adm1da|gash|adm1da2gash|adm1da2meas|{Default State}"
    Assambly="badmLib.dll" Class="badmLib.adm_vffg_h">
4.          <parameters>
5.            <par id="SeparatorMA" val="Model approach" />
6.            <par id="SludgeModel" val="adm1da" />
7.            <par id="ConverterModel" val="adm1da2gash" />
8.            <par id="ConverterMeasModel" val="adm1da2meas" />
9.            <par id="HasBioProcesses" val="true" />
10.           <par id="HasExtBioPar" val="false" />
11.           <par id="HasSludgeOutput" val="true" />
12.           <par id="HasSludgeRatesOutput" val="false" />
13.           <par id="NumberOfGasPorts" val="1" />
14.           <par id="SeparatorRP" val="Reactor parameters" />
15.           <par id="MaxTankVolume" val="1100*5" />
16.           <par id="MaxSludgeVolume" val="1000*5" />
17.           <par id="InitSludgeVolume" val="1000*5" />
18.           <par id="MaxSludgeHeigth" val="8" />
19.           <par id="ExternTemperature" val="false" />
20.           <par id="Temperature" val="55" />
21.           <par id="AutomaticV2COD" val="true" />
22.           <par id="Volume2COD" val="0.0006" />
23.           <par id="adm1da2gashkLa_CO2" val="200.0" />
24.           <par id="adm1da2gashkLa_CH4" val="200.0" />
25.           <par id="adm1da2gashkLa_H2" val="200.0" />
26.           <par id="adm1da2gashkLa_NH3" val="200.0" />
27.           <par id="StateFile" val="{Default State}" />
28.           <par id="StateBlock" val="" />
29.         </parameters>
30.         <Input id="SludgeControl" vis="True" Link="Rezi" LinkIdx="0" />
31.         <Input id="SludgeInflow" vis="True" Link="1st fermenter" LinkIdx="0" />
32.         <Input id="GasFlow0" vis="True" Link="Pipe" LinkIdx="0" />
33.       </Module>
34.     </extra>
35.     <icon rotate="False" flip="True" />
36.   </block>
37.   <block name="FM2M1" blockType="sisiBlock" x="185" y="195" w="25" h="25">
38.     <extra blockID="basmLib.asm_convBlock" HlpIdx="">
39.       <Module id="FM2M1" subtype="fm2substratem1_v2" Assambly="basmLib.dll"
    Class="basmLib.asm_convBlock">
40.         <parameters>
41.           <par id="TS" val="330" />
42.           <par id="NH4" val="0.7" />
43.         </parameters>
```

```
        <Input id="u1" vis="True" Link="From Excel" LinkIdx="0" />
      </Module>
    </extra>
    <icon rotate="False" flip="True" />
  </block>
  <block name="Pipe" blockType="sisiBlock" x="760" y="180" w="35" h="10" bgCol="#64CCFF34" help="simba#_air_pipe.pdf">
    <extra blockID="airlibBlocks.Air_SimpPipeBi" HlpIdx="">
      <Module id="Pipe" subtype="gash" Assambly="airlibBlocks.dll" Class="airlibBlocks.Air_SimpPipeBi">
        <parameters>
          <par id="Model" val="gash" />
          <par id="zeta_e" val="2" />
          <par id="k" val="0.02" />
          <par id="l" val="10" />
          <par id="d" val="0.075" />
          <par id="oneWay" val="false" />
          <par id="simple" val="true" />
          <par id="designVs" val="3.3337947101344607" />
          <par id="nbarrel" val="1" />
          <par id="animate" val="true" />
        </parameters>
        <Output id="y" vis="False" Link="" />
      </Module>
    </extra>
    <defShape typ="Rectangle" lWidth="2" fcol="#FF0020B8" />
    <icon rotate="False" flip="True" />
  </block>
  <block name="setEnvTemp" blockType="sisiBlock" x="200" y="15" w="50" h="20" help="simba#_air_setEnvTemperatureGain.pdf">
    <extra blockID="airlibBlocks.Air_setEnvTemperatureGain" HlpIdx="">
      <Module id="setEnvTemp" subtype="" Assambly="airlibBlocks.dll" Class="airlibBlocks.Air_setEnvTemperatureGain">
        <parameters>
          <par id="dP_min_pipe" val="100" />
          <par id="Zeta_min" val="1" />
        </parameters>
        <Input id="u" vis="True" Link="temperature air" LinkIdx="0" />
        <Output id="y" vis="False" Link="temperature_air" />
      </Module>
    </extra>
    <defShape typ="Arrow" fcol="#FF9B8500" bgcol="#3BEFCD00" var1="0.2" shadow="True" />
    <icon rotate="False" flip="True" />
  </block>
  <block name="temperature air" blockType="sisiBlock" x="95" y="15" w="50" h="20" bgCol="#FF5BA1C4">
    <extra blockID="basmLib.asm_convBlock" HlpIdx="">
```

```xml
        <Module id="temperature air" subtype="setairtemperature" Assambly="basmLib.dll" Class="basmLib.asm_convBlock">
          <parameters>
            <par id="theta" val="20" />
          </parameters>
        </Module>
      </extra>
      <defShape typ="Rectangle" fcol="#FF9B8500" bgcol="#3BEFCD00" shadow="True" />
      <icon rotate="False" flip="True" name=" " x="0" y="0" w="1" h="1" scale="True" />
    </block>
    <block name="1st fermenter" blockType="sisiBlock" x="435" y="230" w="60" h="55" drawFrame="False" help="simba#_biogas_reactor_vfvg_h_en.pdf">
      <extra blockID="badmLib.adm_vfvg_h" HlpIdx="">
        <Module id="1st fermenter" subtype="" Assambly="badmLib.dll" Class="badmLib.adm_vfvg_h">
          <parameters>
            <par id="SeparatorMA" val="Model approach" />
            <par id="SludgeModel" val="adm1da" />
            <par id="ConverterModel" val="adm1da2gash" />
            <par id="ConverterMeasModel" val="adm1da2meas" />
            <par id="HasBioProcesses" val="true" />
            <par id="HasExtBioPar" val="false" />
            <par id="HasSludgeOutput" val="false" />
            <par id="HasSludgeRatesOutput" val="false" />
            <par id="NumberOfGasPorts" val="1" />
            <par id="SeparatorRP" val="Reactor parameters" />
            <par id="MaxTankVolume" val="110" />
            <par id="MaxSludgeVolume" val="100" />
            <par id="InitSludgeVolume" val="100" />
            <par id="MaxSludgeHeigth" val="8" />
            <par id="ExcessPressure" val="2" />
            <par id="ExternTemperature" val="false" />
            <par id="Temperature" val="20" />
            <par id="AutomaticV2COD" val="true" />
            <par id="Volume2COD" val="0.0006" />
            <par id="adm1da2gashkLa_CO2" val="200.0" />
            <par id="adm1da2gashkLa_CH4" val="200.0" />
            <par id="adm1da2gashkLa_H2" val="200.0" />
            <par id="adm1da2gashkLa_NH3" val="200.0" />
            <par id="StateFile" val="{Default State}" />
            <par id="StateBlock" val="" />
          </parameters>
          <Input id="SludgeInflow" vis="True" Link="Mix" LinkIdx="0" />
          <Input id="GasFlow0" vis="True" Link="Pipe1" LinkIdx="0" />
        </Module>
      </extra>
      <icon rotate="False" flip="True" />
    </block>
```

```
      <block name="Pipe1" blockType="sisiBlock" x="755" y="120" w="35" h="10" bgCol="#64CCFF34" help="simba#_air_pipe.pdf">
        <extra blockID="airlibBlocks.Air_SimpPipeBi" HlpIdx="">
          <Module id="Pipe1" subtype="gash" Assambly="airlibBlocks.dll" Class="airlibBlocks.Air_SimpPipeBi">
            <parameters>
              <par id="Model" val="gash" />
              <par id="zeta_e" val="2" />
              <par id="k" val="0.02" />
              <par id="l" val="10" />
              <par id="d" val="0.075" />
              <par id="oneWay" val="false" />
              <par id="simple" val="true" />
              <par id="designVs" val="3.3337947101344607" />
              <par id="nbarrel" val="1" />
              <par id="animate" val="true" />
            </parameters>
            <Output id="y" vis="False" Link="" />
          </Module>
        </extra>
        <defShape typ="Rectangle" lWidth="2" fcol="#FF0020B8" />
        <icon rotate="False" flip="True" />
      </block>
      <block name="Annotation1" blockType="aBlockAnnotation" x="20" y="55" w="110" h="65" drawName="False" help="simba#_wwtp_block_annotation.pdf" page="35">
        <extra anno="&lt;b&gt;Annotation&lt;/b&gt;&#xD;&#xA;&lt;br/&gt; &#xD;&#xA;&lt;br/&gt; &#xD;&#xA;&lt;br/&gt;&#xD;&#xA;Averrage biowaste per 100 person/day&#xD;&#xA;&lt;br/&gt;&#xD;&#xA;&lt;br/&gt;&#xD;&#xA;&#xD;&#xA;" imScale="1" mleft="8" />
        <defShape typ="Rectangle" fcol="#76000000" bgcol="#12000000" shadow="True" />
        <icon rotate="True" flip="False" />
      </block>
      <block name="Level" blockType="sisiBlock" x="870" y="175" w="15" h="20" help="simba#_air_level.pdf">
        <extra blockID="airlibBlocks.Air_LevelUni" HlpIdx="">
          <Module id="Level" subtype="gash" Assambly="airlibBlocks.dll" Class="airlibBlocks.Air_LevelUni">
            <parameters>
              <par id="Model" val="gash" />
              <par id="n_N2" val="0.779713" />
              <par id="n_O2" val="0.209919" />
              <par id="n_CO2" val="0.000381" />
              <par id="n_CH4" val="0" />
              <par id="n_H2" val="0" />
              <par id="n_NH3" val="0" />
              <par id="n_N2O" val="0" />
              <par id="n_SO2" val="0" />
```

```xml
          <par id="n_H2S" val="0" />
          <par id="RH" val="70" />
          <par id="calcP" val="false" />
          <par id="h0" val="0" />
          <par id="hw" val="4" />
          <par id="extP" val="false" />
          <par id="P" val="1013.25+5" />
          <par id="Pmin" val="1000" />
          <par id="Pmax" val="1500" />
          <par id="isSource" val="false" />
          <par id="animate" val="false" />
          <par id="T1" val="2.0/(24.0*60.0)" />
        </parameters>
        <Input id="Tair" vis="False" Link="temperature_air" />
        <Input id="i" vis="True" Link="Pipe" LinkIdx="1" />
      </Module>
    </extra>
    <defShape typ="Rectangle" lWidth="2" fcol="#FF0020B8" var1="0.5" />
    <icon rotate="False" flip="True" />
  </block>
  <block name="Mix" blockType="sisiBlock" x="320" y="210" w="45" h="95" drawFrame="False" help="SIMBA#_Global_Block_MIx.pdf" page="28">
    <extra blockID="basmLib.asm_mixBlock" HlpIdx="">
      <Module id="Mix" subtype="adm1da" Assambly="basmLib.dll" Class="basmLib.asm_mixBlock">
        <parameters>
          <par id="Model" val="adm1da" />
          <par id="type" val="classic" />
          <par id="nin" val="3" />
        </parameters>
        <Input id="u0" vis="True" Link="Silages" LinkIdx="0" />
        <Input id="u1" vis="True" Link="Silages1" LinkIdx="0" />
        <Input id="u2" vis="True" Link="Separator RF  " LinkIdx="0" />
      </Module>
    </extra>
    <icon rotate="False" flip="True" />
  </block>
  <block name="Silages1" blockType="sisiBlock" x="240" y="260" w="25" h="25" help="simba#_biogas_inflow_silages_v2_en.pdf">
    <extra blockID="basmLib.asm_convBlock" HlpIdx="">
      <Module id="Silages1" subtype="tg_adm1da_v_silages_v2" Assambly="basmLib.dll" Class="basmLib.asm_convBlock" resObjSetName="Grass silage - blooming">
        <parameters>
          <par id="COD_deg_type" val="Indicate biogas potential" />
          <par id="V_m" val="0.022413" />
          <par id="CH4_cod_2_mol" val="64" />
          <par id="BGP" val="560" />
          <par id="BMP" val="302.5" />
```

```xml
          <par id="aXI" val="0.2869" />
          <par id="fOTSrf" val="0.9" />
          <par id="fsOTS" val="0" />
          <par id="ffOTS" val="1" />
          <par id="aSi" val="0" />
          <par id="fRF" val="0.295" />
          <par id="fRP" val="0.07" />
          <par id="fRFe" val="0.08" />
          <par id="fRA" val="0.105" />
          <par id="Temp" val="20" />
          <par id="ph" val="4" />
          <par id="KS43" val="0" />
          <par id="FFS" val="5" />
          <par id="roh_CH" val="s_fp('adm1da','roh_CH')" />
          <par id="roh_PR" val="s_fp('adm1da','roh_PR')" />
          <par id="roh_LI" val="s_fp('adm1da','roh_LI')" />
          <par id="roh_MI" val="s_fp('adm1da','roh_MI')" />
          <par id="roh_AC" val="s_fp('adm1da','roh_AC')" />
          <par id="roh_H2O" val="1000" />
          <par id="MP_CH" val="s_fp('adm1da','MP_CH')" />
          <par id="MP_PR" val="s_fp('adm1da','MP_PR')" />
          <par id="MP_LI" val="s_fp('adm1da','MP_LI')" />
          <par id="MP_AC" val="s_fp('adm1da','MP_AC')" />
          <par id="Kw_35" val="s_fp('adm1da','Kw_35')" />
          <par id="M_Sva" val="s_fp('adm1da','M_Sva')" />
          <par id="M_Sbu" val="s_fp('adm1da','M_Sbu')" />
          <par id="M_Spro" val="s_fp('adm1da','M_Spro')" />
          <par id="M_Sac" val="s_fp('adm1da','M_Sac')" />
          <par id="M_Sh2" val="s_fp('adm1da','M_Sh2')" />
          <par id="M_XB" val="s_fp('adm1da','M_XB')" />
          <par id="M_Xch" val="s_fp('adm1da','M_Xch')" />
          <par id="M_Xpr" val="s_fp('adm1da','M_Xpr')" />
          <par id="M_Xli" val="s_fp('adm1da','M_Xli')" />
          <par id="Kava" val="s_fp('adm1da','Kava')" />
          <par id="Kabu" val="s_fp('adm1da','Kabu')" />
          <par id="Kapro" val="s_fp('adm1da','Kapro')" />
          <par id="Kaac" val="s_fp('adm1da','Kaac')" />
          <par id="Kaco2_35" val="s_fp('adm1da','Kaco2_35')" />
          <par id="Kain_35" val="s_fp('adm1da','Kain_35')" />
          <par id="N_aa" val="s_fp('adm1da','N_aa')" />
        </parameters>
        <Input id="u1" vis="True" Link="FM2M2" LinkIdx="0" />
      </Module>
    </extra>
    <icon rotate="False" flip="True" />
  </block>
  <block name="FM2M2" blockType="sisiBlock" x="185" y="260" w="25" h="25">
    <extra blockID="basmLib.asm_convBlock" HlpIdx="">
```

```
        <Module id="FM2M2" subtype="fm2substratem1_v2" Assambly="basmLib.dll" Class="basmLib.asm_convBlock">
          <parameters>
            <par id="TS" val="330" />
            <par id="NH4" val="0.7" />
          </parameters>
          <Input id="u1" vis="True" Link="Switch" LinkIdx="0" />
        </Module>
      </extra>
      <icon rotate="False" flip="True" />
    </block>
    <block name="plant" blockType="sisiBlock" x="20" y="325" w="30" h="30" bgCol="#F2EFEFEF" help="SIMBA#_Global_Block_Temperature_Zero.pdf" page="32">
      <extra blockID="basmLib.asm_constBlock" HlpIdx="">
        <Module id="plant" subtype="Signal,1" Assambly="basmLib.dll" Class="basmLib.asm_constBlock">
          <parameters>
            <par id="Model" val="Signal" />
            <par id="Dimmension" val="1" />
            <par id="Constant_0" val="0.2" />
            <par id="unit_0" val="-" />
            <par id="sel_0" val="User defined" />
            <par id="displayUnits" val="false" />
          </parameters>
        </Module>
      </extra>
      <defShape typ="Rectangle" fcol="#FF606060" bgcol="#26606060" shadow="True" />
      <icon rotate="False" flip="True" />
    </block>
    <block name="Switch" blockType="sisiBlock" x="135" y="290" w="15" h="65" drawFrame="False" fgCol="#FFB7B7B7" help="simba#_manual_referenz.xps" page="58">
      <extra blockID="signalLib.switchBlock" HlpIdx="">
        <Module id="Switch" subtype="2" Assambly="signalLib.dll" Class="signalLib.switchBlock">
          <parameters>
            <par id="inputIdx" val="Input 2" />
            <par id="switchByInput" val="false" />
            <par id="numberOfInputs" val="2" />
            <par id="n" val="1" />
          </parameters>
          <Input id="u0" vis="True" Link="plant" LinkIdx="0" />
          <Input id="u1" vis="True" Link="plant1" LinkIdx="0" />
        </Module>
      </extra>
      <icon rotate="False" flip="True" />
    </block>
    <block name="Gasproduktion" blockType="sisiBlock" x="1285" y="200" w="55" h="15" help="SIMBA#_Global_Block_Sink.pdf" page="30">
```

```xml
      <extra blockID="basmLib.asm_sinkBlock" HlpIdx="">
        <Anno>DQo=</Anno>
        <Module id="Gasproduktion" subtype="gash" Assambly="basmLib.dll" Class="basmLib.asm_sinkBlock">
          <parameters>
            <par id="Model" val="gash" />
            <par id="size" val="12" />
            <par id="sinktype" val="Other" />
            <par id="display" val="false" />
            <par id="feedThrough" val="false" />
          </parameters>
          <Input id="u" vis="True" Link="meas1" LinkIdx="0" />
        </Module>
      </extra>
      <defShape typ="Arrow" lWidth="1.5" fcol="#FF0049C3" bgcol="#180049C3" var1="0.2" shadow="True" />
      <icon rotate="True" flip="False" />
    </block>
    <block name="Gasanalyse" blockType="sisiBlock" x="1065" y="90" w="65" h="100" help="simba#_manual_refad_en.xps" page="10">
      <extra blockID="basmLib.asm_convBlock" HlpIdx="">
        <Module id="Gasanalyse" subtype="gasanalyseV2a_gash" Assambly="basmLib.dll" Class="basmLib.asm_convBlock">
          <Input id="Gas_in" vis="True" Link="Mix1" LinkIdx="0" />
        </Module>
      </extra>
      <icon rotate="False" flip="True" />
    </block>
    <block name="CH4" blockType="sisiBlock" x="1240" y="100" w="20" h="10" help="SIMBA#_Global_Block_Sink.pdf" page="30">
      <extra blockID="basmLib.asm_sinkBlock" HlpIdx="">
        <Anno>DQo=</Anno>
        <Module id="CH4" subtype="Signal,1" Assambly="basmLib.dll" Class="basmLib.asm_sinkBlock">
          <parameters>
            <par id="Model" val="Signal" />
            <par id="size" val="1" />
            <par id="sinktype" val="Effluent" />
            <par id="display" val="false" />
            <par id="feedThrough" val="false" />
            <par id="unit_0" val="-" />
          </parameters>
          <Input id="u" vis="True" Link="Gasanalyse" LinkIdx="0" />
        </Module>
      </extra>
      <defShape typ="Arrow" lWidth="1.5" fcol="#FF0049C3" bgcol="#180049C3" var1="0.2" shadow="True" />
      <icon rotate="True" flip="False" />
    </block>
```

```xml
<block name="O2" blockType="sisiBlock" x="1240" y="120" w="20" h="10" help="SIMBA#_Global_Block_Sink.pdf" page="30">
  <extra blockID="basmLib.asm_sinkBlock" HlpIdx="">
    <Anno>DQo=</Anno>
    <Module id="O2" subtype="Signal,1" Assambly="basmLib.dll" Class="basmLib.asm_sinkBlock">
      <parameters>
        <par id="Model" val="Signal" />
        <par id="size" val="1" />
        <par id="sinktype" val="Effluent" />
        <par id="display" val="false" />
        <par id="feedThrough" val="false" />
        <par id="unit_0" val="-" />
      </parameters>
      <Input id="u" vis="True" Link="Gasanalyse" LinkIdx="2" />
    </Module>
  </extra>
  <defShape typ="Arrow" lWidth="1.5" fcol="#FF0049C3" bgcol="#180049C3" var1="0.2" shadow="True" />
  <icon rotate="True" flip="False" />
</block>
<block name="H2S2" blockType="sisiBlock" x="1240" y="140" w="20" h="10" help="SIMBA#_Global_Block_Sink.pdf" page="30">
  <extra blockID="basmLib.asm_sinkBlock" HlpIdx="">
    <Anno>DQo=</Anno>
    <Module id="H2S2" subtype="Signal,1" Assambly="basmLib.dll" Class="basmLib.asm_sinkBlock">
      <parameters>
        <par id="Model" val="Signal" />
        <par id="size" val="1" />
        <par id="sinktype" val="Effluent" />
        <par id="display" val="false" />
        <par id="feedThrough" val="false" />
        <par id="unit_0" val="-" />
      </parameters>
      <Input id="u" vis="True" Link="Gasanalyse" LinkIdx="4" />
    </Module>
  </extra>
  <defShape typ="Arrow" lWidth="1.5" fcol="#FF0049C3" bgcol="#180049C3" var1="0.2" shadow="True" />
  <icon rotate="True" flip="False" />
</block>
<block name="H2" blockType="sisiBlock" x="1195" y="130" w="20" h="10" help="SIMBA#_Global_Block_Sink.pdf" page="30">
  <extra blockID="basmLib.asm_sinkBlock" HlpIdx="">
    <Anno>DQo=</Anno>
    <Module id="H2" subtype="Signal,1" Assambly="basmLib.dll" Class="basmLib.asm_sinkBlock">
      <parameters>
```

```
              <par id="Model" val="Signal" />
              <par id="size" val="1" />
              <par id="sinktype" val="Effluent" />
              <par id="display" val="false" />
              <par id="feedThrough" val="false" />
              <par id="unit_0" val="-" />
            </parameters>
            <Input id="u" vis="True" Link="Gasanalyse" LinkIdx="3" />
          </Module>
        </extra>
        <defShape typ="Arrow" lWidth="1.5" fcol="#FF0049C3" bgcol="#180049C3" var1="0.2" shadow="True" />
        <icon rotate="True" flip="False" />
      </block>
      <block name="CH4_tr" blockType="sisiBlock" x="1240" y="170" w="20" h="10" help="SIMBA#_Global_Block_Sink.pdf" page="30">
        <extra blockID="basmLib.asm_sinkBlock" HlpIdx="">
          <Anno>DQo=</Anno>
          <Module id="CH4_tr" subtype="Signal,1" Assambly="basmLib.dll" Class="basmLib.asm_sinkBlock">
            <parameters>
              <par id="Model" val="Signal" />
              <par id="size" val="1" />
              <par id="sinktype" val="Effluent" />
              <par id="display" val="false" />
              <par id="feedThrough" val="false" />
              <par id="unit_0" val="-" />
            </parameters>
            <Input id="u" vis="True" Link="Gasanalyse" LinkIdx="7" />
          </Module>
        </extra>
        <defShape typ="Arrow" lWidth="1.5" fcol="#FF0049C3" bgcol="#180049C3" var1="0.2" shadow="True" />
        <icon rotate="True" flip="False" />
      </block>
      <block name="CO2" blockType="sisiBlock" x="1195" y="110" w="20" h="10" help="SIMBA#_Global_Block_Sink.pdf" page="30">
        <extra blockID="basmLib.asm_sinkBlock" HlpIdx="">
          <Anno>DQo=</Anno>
          <Module id="CO2" subtype="Signal,1" Assambly="basmLib.dll" Class="basmLib.asm_sinkBlock">
            <parameters>
              <par id="Model" val="Signal" />
              <par id="size" val="1" />
              <par id="sinktype" val="Effluent" />
              <par id="display" val="false" />
              <par id="feedThrough" val="false" />
              <par id="unit_0" val="-" />
            </parameters>
```

```
          <Input id="u" vis="True" Link="Gasanalyse" LinkIdx="1" />
        </Module>
      </extra>
      <defShape typ="Arrow" lWidth="1.5" fcol="#FF0049C3" bgcol="#180049C3" var1="0.2" shadow="True" />
      <icon rotate="True" flip="False" />
    </block>
    <block name="Qch4" blockType="sisiBlock" x="1380" y="245" w="20" h="40" help="simba#_manual_referenz.xps" page="48">
      <extra blockID="signalLib.MULn" HlpIdx="">
        <Module id="Qch4" subtype="" Assambly="signalLib.dll" Class="signalLib.MULn">
          <parameters>
            <par id="n" val="1" />
            <par id="Input signs" val="**/" />
          </parameters>
          <Input id="u0" vis="True" Link="Gasanalyse" LinkIdx="0" />
          <Input id="u1" vis="True" Link="meas1" LinkIdx="1" />
          <Input id="u2" vis="True" Link="const1" LinkIdx="0" />
        </Module>
      </extra>
      <defShape typ="Rectangle" fcol="#FF606060" bgcol="#26606060" shadow="True" />
      <icon rotate="False" flip="False" />
    </block>
    <block name="meas1" blockType="sisiBlock" x="1240" y="225" w="25" h="55" drawFrame="False" drawName="False" bgCol="#D3F0F3F5" help="SIMBA#_Global_Block_Meas.pdf" page="33">
      <extra blockID="basmLib.asm_measBlock" HlpIdx="">
        <Anno>DQo=</Anno>
        <Module id="meas1" subtype="gash" Assambly="basmLib.dll" Class="basmLib.asm_measBlock">
          <parameters>
            <par id="Model" val="gash" />
            <par id="horz" val="false" />
            <par id="label" val="false" />
            <par id="n_N2" val="false" />
            <par id="n_O2" val="false" />
            <par id="n_CO2" val="false" />
            <par id="n_CH4" val="false" />
            <par id="n_H2" val="false" />
            <par id="n_NH3" val="false" />
            <par id="n_N2O" val="false" />
            <par id="n_SO2" val="false" />
            <par id="n_H2S" val="false" />
            <par id="n_H2O" val="false" />
            <par id="theta" val="false" />
            <par id="Q" val="true" />
            <par id="P: R" val="false" />
```

```
                <par id="P: Roh" val="false" />
                <par id="P: COD" val="false" />
                <par id="P: N" val="false" />
                <par id="P: P" val="false" />
                <par id="P: C" val="false" />
                <par id="P: S" val="false" />
                <par id="P: M" val="false" />
                <par id="P: a" val="false" />
                <par id="P: b" val="false" />
                <par id="P: c" val="false" />
                <par id="P: d" val="false" />
                <par id="isDyn" val="false" />
                <par id="T" val="1/(24*60*60)" />
              </parameters>
              <Input id="u" vis="True" Link="Gasanalyse" LinkIdx="5" />
            </Module>
          </extra>
          <icon rotate="False" flip="True" />
        </block>
        <block name="n" blockType="aBranchBlock" x="1220" y="100" w="10" h="10" drawFrame="False" drawName="False">
          <extra btype="1" lineType="Signal" lineExtra="1" />
          <icon rotate="False" flip="False" />
        </block>
        <block name="const1" blockType="sisiBlock" x="1310" y="285" w="35" h="20" drawName="False" bgCol="#F2EFEFEF" help="SIMBA#_Global_Block_Temperature_Zero.pdf" page="32">
          <extra blockID="basmLib.asm_constBlock" HlpIdx="">
            <Module id="const1" subtype="Signal,1" Assambly="basmLib.dll" Class="basmLib.asm_constBlock">
              <parameters>
                <par id="Model" val="Signal" />
                <par id="Dimmension" val="1" />
                <par id="Constant_0" val="100" />
                <par id="unit_0" val="-" />
                <par id="sel_0" val="User defined" />
                <par id="displayUnits" val="false" />
              </parameters>
            </Module>
          </extra>
          <defShape typ="Rectangle" fcol="#FF606060" bgcol="#26606060" shadow="True" />
          <icon rotate="False" flip="True" />
        </block>
        <block name="Annotation3" blockType="aBlockAnnotation" x="1265" y="335" w="250" h="50" drawName="False" help="simba#_wwtp_block_annotation.pdf" page="35">
          <extra anno="&lt;b&gt;Annotation&lt;/b&gt;&#xD;&#xA;&lt;br/&gt;&#xD;&#xA;&lt;br/&gt;
```

```
&#xD;&#xA;&lt;br/&gt;&#xD;&#xA;Gasanalyse&#xD;&#xA;&lt;br/&gt;&#xD;&#xA;Calculation of gas amounts (CH4, CO2, H2, O2)&#xD;&#xA;&#xD;&#xA;" imScale="1" mleft="8" />
        <defShape typ="Rectangle" fcol="#76000000" bgcol="#12000000" shadow="True" />
        <icon rotate="True" flip="False" />
      </block>
      <block name="Biogas into kWh1" blockType="sisiBlock" x="1445" y="250" w="35" h="30" help="simba#_manual_referenz.xps" page="47">
        <extra blockID="signalLib.Gain" HlpIdx="">
          <Module id="Biogas into kWh1" subtype="" Assambly="signalLib.dll" Class="signalLib.Gain">
            <parameters>
              <par id="n" val="1" />
              <par id="K" val="10.5" />
              <par id="inunit" val="-" />
              <par id="outunit" val="-" />
            </parameters>
            <Input id="u" vis="True" Link="Qch4" LinkIdx="0" />
          </Module>
        </extra>
        <defShape typ="Arrow" fcol="#FF606060" bgcol="#26606060" var1="0.2" shadow="True" />
        <icon rotate="True" flip="False" />
      </block>
      <block name="kWh1" blockType="sisiBlock" x="1505" y="260" w="50" h="15" help="SIMBA#_Global_Block_Sink.pdf" page="30">
        <extra blockID="basmLib.asm_sinkBlock" HlpIdx="">
          <Anno>DQo=</Anno>
          <Module id="kWh1" subtype="Signal,1" Assambly="basmLib.dll" Class="basmLib.asm_sinkBlock">
            <parameters>
              <par id="Model" val="Signal" />
              <par id="size" val="1" />
              <par id="sinktype" val="Other" />
              <par id="display" val="true" />
              <par id="feedThrough" val="false" />
              <par id="unit_0" val="-" />
            </parameters>
            <Input id="u" vis="True" Link="Biogas into kWh1" LinkIdx="0" />
          </Module>
        </extra>
        <defShape typ="Arrow" lWidth="1.5" fcol="#FF0049C3" bgcol="#180049C3" var1="0.2" shadow="True" />
        <icon rotate="True" flip="False" />
      </block>
      <block name="n1" blockType="aBranchBlock" x="1415" y="260" w="10" h="10" drawFrame="False" drawName="False">
        <extra btype="1" lineType="Signal" lineExtra="1" />
        <icon rotate="False" flip="False" />
```

```
      </block>
      <block name="CH4_volume" blockType="sisiBlock" x="1505" y="315" w="50" h="15" help="SIMBA#_Global_Block_Sink.pdf" page="30">
        <extra blockID="basmLib.asm_sinkBlock" HlpIdx="">
          <Anno>DQo=</Anno>
          <Module id="CH4_volume" subtype="Signal,1" Assambly="basmLib.dll" Class="basmLib.asm_sinkBlock">
            <parameters>
              <par id="Model" val="Signal" />
              <par id="size" val="1" />
              <par id="sinktype" val="Other" />
              <par id="display" val="true" />
              <par id="feedThrough" val="false" />
              <par id="unit_0" val="-" />
            </parameters>
            <Input id="u" vis="True" Link="Qch4" LinkIdx="0" />
          </Module>
        </extra>
        <defShape typ="Arrow" lWidth="1.5" fcol="#FF0049C3" bgcol="#180049C3" var1="0.2" shadow="True" />
        <icon rotate="True" flip="False" />
      </block>
      <block name="Silages" blockType="sisiBlock" x="235" y="195" w="25" h="25" help="simba#_biogas_inflow_silages_v2_en.pdf">
        <extra blockID="basmLib.asm_convBlock" HlpIdx="">
          <Module id="Silages" subtype="tg_adm1da_v_silages_v2" Assambly="basmLib.dll" Class="basmLib.asm_convBlock" resObjSetName="Maize silage - milk ripeness">
            <parameters>
              <par id="COD_deg_type" val="Indicate biogas potential" />
              <par id="V_m" val="0.022413" />
              <par id="CH4_cod_2_mol" val="64" />
              <par id="BGP" val="615" />
              <par id="BMP" val="337.4" />
              <par id="aXI" val="0.2182" />
              <par id="fOTSrf" val="0.95" />
              <par id="fsOTS" val="0" />
              <par id="ffOTS" val="1" />
              <par id="aSi" val="0" />
              <par id="fRF" val="0.235" />
              <par id="fRP" val="0.07" />
              <par id="fRFe" val="0.08" />
              <par id="fRA" val="0.105" />
              <par id="Temp" val="20" />
              <par id="ph" val="7" />
              <par id="KS43" val="20" />
              <par id="FFS" val="0" />
              <par id="roh_CH" val="s_fp('adm1da','roh_CH')" />
              <par id="roh_PR" val="s_fp('adm1da','roh_PR')" />
```

```
            <par id="roh_LI" val="s_fp('adm1da','roh_LI')" />
            <par id="roh_MI" val="s_fp('adm1da','roh_MI')" />
            <par id="roh_AC" val="s_fp('adm1da','roh_AC')" />
            <par id="roh_H2O" val="1000" />
            <par id="MP_CH" val="s_fp('adm1da','MP_CH')" />
            <par id="MP_PR" val="s_fp('adm1da','MP_PR')" />
            <par id="MP_LI" val="s_fp('adm1da','MP_LI')" />
            <par id="MP_AC" val="s_fp('adm1da','MP_AC')" />
            <par id="Kw_35" val="s_fp('adm1da','Kw_35')" />
            <par id="M_Sva" val="s_fp('adm1da','M_Sva')" />
            <par id="M_Sbu" val="s_fp('adm1da','M_Sbu')" />
            <par id="M_Spro" val="s_fp('adm1da','M_Spro')" />
            <par id="M_Sac" val="s_fp('adm1da','M_Sac')" />
            <par id="M_Sh2" val="s_fp('adm1da','M_Sh2')" />
            <par id="M_XB" val="s_fp('adm1da','M_XB')" />
            <par id="M_Xch" val="s_fp('adm1da','M_Xch')" />
            <par id="M_Xpr" val="s_fp('adm1da','M_Xpr')" />
            <par id="M_Xli" val="s_fp('adm1da','M_Xli')" />
            <par id="Kava" val="s_fp('adm1da','Kava')" />
            <par id="Kabu" val="s_fp('adm1da','Kabu')" />
            <par id="Kapro" val="s_fp('adm1da','Kapro')" />
            <par id="Kaac" val="s_fp('adm1da','Kaac')" />
            <par id="Kaco2_35" val="s_fp('adm1da','Kaco2_35')" />
            <par id="Kain_35" val="s_fp('adm1da','Kain_35')" />
            <par id="N_aa" val="s_fp('adm1da','N_aa')" />
          </parameters>
          <Input id="u1" vis="True" Link="FM2M1" LinkIdx="0" />
        </Module>
      </extra>
      <icon rotate="False" flip="True" />
    </block>
    <block name="From Excel" blockType="sisiBlock" x="15" y="195" w="55" h="30" bgCol="#CFFFEC7A" help="SIMBA#_Global_Block_FromXLSX.pdf" page="8">
      <extra blockID="basmLib.gen_infileExcelBlock" HlpIdx="">
        <Module id="From Excel" subtype="Signal" Assambly="basmLib.dll" Class="basmLib.gen_infileExcelBlock">
          <parameters>
            <par id="Model" val="Signal" />
            <par id="sigsize" val="1" />
            <par id="qunit" val="m^3/d" />
            <par id="exfname" val="Table1_Fluctuation_of_Biomass.xlsx" />
            <par id="tabname" val="SimbaFlu" />
            <par id="interpol" val="First Order (linear)" />
            <par id="afterlast" val="cyclic repitition" />
            <par id="average" val="false" />
            <par id="isexact" val="true" />
            <par id="showicon" val="true" />
            <par id="showlab" val="false" />
            <par id="Test" val="0.0" />
```

```
        <par id="Open.." val="0.0" />
        <par id="Copy file" val="0.0" />
        <par id="Buffer" val="1000" />
        <par id="labs" val="Biomasse;" />
      </parameters>
    </Module>
  </extra>
  <defShape typ="Arrow" lWidth="1.5" fcol="#FF9B8500" bgcol="#3BEFCD00" var1="0.2" shadow="True" />
  <icon rotate="False" flip="True" />
</block>
<block name="Level1" blockType="sisiBlock" x="870" y="115" w="15" h="20" help="simba#_air_level.pdf">
  <extra blockID="airlibBlocks.Air_LevelUni" HlpIdx="">
    <Module id="Level1" subtype="gash" Assambly="airlibBlocks.dll" Class="airlibBlocks.Air_LevelUni">
      <parameters>
        <par id="Model" val="gash" />
        <par id="n_N2" val="0.779713" />
        <par id="n_O2" val="0.209919" />
        <par id="n_CO2" val="0.000381" />
        <par id="n_CH4" val="0" />
        <par id="n_H2" val="0" />
        <par id="n_NH3" val="0" />
        <par id="n_N2O" val="0" />
        <par id="n_SO2" val="0" />
        <par id="n_H2S" val="0" />
        <par id="RH" val="70" />
        <par id="calcP" val="false" />
        <par id="h0" val="0" />
        <par id="hw" val="4" />
        <par id="extP" val="false" />
        <par id="P" val="1013.25+5" />
        <par id="Pmin" val="1000" />
        <par id="Pmax" val="1500" />
        <par id="isSource" val="false" />
        <par id="animate" val="false" />
        <par id="T1" val="2.0/(24.0*60.0)" />
      </parameters>
      <Input id="Tair" vis="False" Link="temperature_air" />
      <Input id="i" vis="True" Link="Pipe1" LinkIdx="1" />
    </Module>
  </extra>
  <defShape typ="Rectangle" lWidth="2" fcol="#FF0020B8" var1="0.5" />
  <icon rotate="False" flip="True" />
</block>
<block name="Mix1" blockType="sisiBlock" x="1005" y="130" w="25" h="50" drawFrame="False" help="SIMBA#_Global_Block_MIx.pdf" page="28">
  <extra blockID="basmLib.asm_mixBlock" HlpIdx="">
```

```
          <Module id="Mix1" subtype="gash" Assambly="basmLib.dll" Class="basmLib.asm_mixBlock">
            <parameters>
              <par id="Model" val="gash" />
              <par id="type" val="classic" />
              <par id="nin" val="2" />
            </parameters>
            <Input id="u0" vis="True" Link="Level1" LinkIdx="0" />
            <Input id="u1" vis="True" Link="Level" LinkIdx="0" />
          </Module>
        </extra>
        <icon rotate="False" flip="True" />
      </block>
      <block name="plant1" blockType="sisiBlock" x="20" y="380" w="30" h="30" bgCol="#F2EFEFEF" help="SIMBA#_Global_Block_Temperature_Zero.pdf" page="32">
        <extra blockID="basmLib.asm_constBlock" HlpIdx="">
          <Module id="plant1" subtype="Signal,1" Assambly="basmLib.dll" Class="basmLib.asm_constBlock">
            <parameters>
              <par id="Model" val="Signal" />
              <par id="Dimmension" val="1" />
              <par id="Constant_0" val="1" />
              <par id="unit_0" val="-" />
              <par id="sel_0" val="User defined" />
              <par id="displayUnits" val="false" />
            </parameters>
          </Module>
        </extra>
        <defShape typ="Rectangle" fcol="#FF606060" bgcol="#26606060" shadow="True" />
        <icon rotate="False" flip="True" />
      </block>
      <block name="Sludge_overflow1" blockType="sisiBlock" x="945" y="235" w="20" h="10" help="SIMBA#_Global_Block_Sink.pdf" page="30">
        <extra blockID="basmLib.asm_sinkBlock" HlpIdx="">
          <Anno>DQo=</Anno>
          <Module id="Sludge_overflow1" subtype="adm1da" Assambly="basmLib.dll" Class="basmLib.asm_sinkBlock">
            <parameters>
              <par id="Model" val="adm1da" />
              <par id="size" val="43" />
              <par id="sinktype" val="Sludge" />
              <par id="display" val="true" />
              <par id="feedThrough" val="false" />
            </parameters>
            <Input id="u" vis="True" Link="main fermenter" LinkIdx="0" />
          </Module>
        </extra>
```

```
729. <defShape typ="Arrow" lWidth="1.5" fcol="#FF0049C3" bgcol="#180049C3" var1="0.2" shadow="True" />
730. <icon rotate="True" flip="False" />
731. </block>
732. <block name="Separator RF  " blockType="sisiBlock" x="880" y="300" w="45" h="30" drawFrame="False" help="simba#_biogas_separator_rf_en.pdf">
733. <extra blockID="badmLib.adm_sep_as1" HlpIdx="">
734. <Module id="Separator RF  " subtype="adm1da" Assambly="badmLib.dll" Class="badmLib.adm_sep_as1">
735. <parameters>
736. <par id="SeparatorMA" val="Model approach" />
737. <par id="SludgeModel" val="adm1da" />
738. <par id="ControlPortPosition" val="Top" />
739. <par id="SeparatorDP" val="Device parameter" />
740. <par id="ControlTypeList" val="Flow of thick separate" />
741. <par id="TSIndicationList" val="Specify TSS" />
742. <par id="FinalTSContent" val="230" />
743. <par id="FinalOTSContent" val="150" />
744. </parameters>
745. <Input id="us1" vis="True" Link="main fermenter" LinkIdx="1" />
746. <Input id="uc1" vis="True" Link="VolStromDick" LinkIdx="0" />
747. </Module>
748. </extra>
749. <icon rotate="True" flip="False" />
750. </block>
751. <block name="Dickseparat" blockType="sisiBlock" x="1075" y="380" w="20" h="10" help="SIMBA#_Global_Block_Sink.pdf" page="30">
752. <extra blockID="basmLib.asm_sinkBlock" HlpIdx="">
753. <Anno>DQo=</Anno>
754. <Module id="Dickseparat" subtype="adm1da" Assambly="basmLib.dll" Class="basmLib.asm_sinkBlock">
755. <parameters>
756. <par id="Model" val="adm1da" />
757. <par id="size" val="43" />
758. <par id="sinktype" val="Sludge" />
759. <par id="display" val="true" />
760. <par id="feedThrough" val="false" />
761. </parameters>
762. <Input id="u" vis="True" Link="Separator RF  " LinkIdx="1" />
763. </Module>
764. </extra>
765. <defShape typ="Arrow" lWidth="1.5" fcol="#FF0049C3" bgcol="#180049C3" var1="0.2" shadow="True" />
766. <icon rotate="True" flip="False" />
767. </block>
768. <block name="VolStromDick" blockType="sisiBlock" x="850" y="260" w="30" h="30" bgCol="#F2EFEFEF" help="SIMBA#_Global_Block_Temperature_Zero.pdf" page="32">
769. <extra blockID="basmLib.asm_constBlock" HlpIdx="">
```

```
        <Module id="VolStromDick" subtype="Signal,1" Assambly="basmLib.dll" Class="basmLib.asm_constBlock">
          <parameters>
            <par id="Model" val="Signal" />
            <par id="Dimmension" val="1" />
            <par id="Constant_0" val="15" />
            <par id="unit_0" val="-" />
            <par id="sel_0" val="User defined" />
            <par id="displayUnits" val="false" />
          </parameters>
        </Module>
      </extra>
      <defShape typ="Rectangle" fcol="#FF606060" bgcol="#26606060" shadow="True" />
      <icon rotate="False" flip="True" />
    </block>
    <block name="Rezi" blockType="sisiBlock" x="530" y="305" w="30" h="30" bgCol="#F2EFEFEF" help="SIMBA#_Global_Block_Temperature_Zero.pdf" page="32">
      <extra blockID="basmLib.asm_constBlock" HlpIdx="">
        <Module id="Rezi" subtype="Signal,1" Assambly="basmLib.dll" Class="basmLib.asm_constBlock">
          <parameters>
            <par id="Model" val="Signal" />
            <par id="Dimmension" val="1" />
            <par id="Constant_0" val="35" />
            <par id="unit_0" val="-" />
            <par id="sel_0" val="User defined" />
            <par id="displayUnits" val="false" />
          </parameters>
        </Module>
      </extra>
      <defShape typ="Rectangle" fcol="#FF606060" bgcol="#26606060" shadow="True" />
      <icon rotate="False" flip="True" />
```
